# Endosymbiotic algal photosynthesis shapes diel transcriptome architecture in its ciliate host *Paramecium bursaria*

**DOI:** 10.64898/2026.03.31.715701

**Authors:** Md Mostafa Kamal, Yu-Hsuan Cheng, Chia-Ling Yang, Chien-Fu Jeff Liu, Chuan Ku, Jun-Yi Leu

## Abstract

How photosynthetic endosymbionts reorganize host daily regulation remains unclear. *Paramecium bursaria* displays pronounced day-night behaviors, but whether its algal symbionts drive host temporal programs has been unresolved. We compared host gene expressions across a 24-hour light-dark cycle in symbiotic and aposymbiotic cells. Symbiotic cells exhibit an expanded and highly temporally ordered diel transcriptome compared with aposymbiotic cells. These rhythmic programs encompass motility, signaling, metabolism, and growth regulation, consistent with observed behaviors. Symbiosis-associated rhythmic programs recruit gene families encoding post-translational regulatory domains, including kinases, ubiquitin-related factors, WD40 scaffolds, and calcium-binding proteins, despite lacking recognizable canonical clock genes. Disrupting photosynthesis with paraquat altered these temporal profiles, shifting them toward an aposymbiotic-like state. A distantly related ciliate, *Tetrahymena utriculariae*, with an independently evolved symbiosis, showed similar symbiosis-associated daily programs, suggesting that photosynthetic endosymbionts can act as important organizers of host daily gene regulation in endosymbiotic protists.

## Introduction

Time-of-day transcriptional programs are widespread among unicellular eukaryotes, including algae and diverse protists (*1–3*). Under light-dark (LD) cycles, a diel pattern can emerge from interactions between environmental cues and endogenous factors. These integrated programs enable cells to coordinate different cellular processes, including energy metabolism, redox balance, transcription, translation, and protein turnover (*4*). Metabolic flux often acts as an entraining input for gene expression cycles, including in lineages where canonical circadian regulators are not readily identifiable (*5*). For example, non-photosynthetic protists can exhibit diel reprogramming when exposed to rhythmic metabolic environments (*3*). In photosynthetic organisms, rhythmic redox and sugar production act as strong time-of-day signals that synchronize cellular functions to the photoperiod (*4, 6*). Importantly, diel patterns measured under LD entrainment do not by themselves demonstrate free-running circadian rhythmicity. Instead, they reveal how predictable daily inputs can impose genome-wide temporal organization, as shown in systems where LD-driven transcriptional rhythms persist even when endogenous circadian oscillations are disrupted (*7*).

Symbiotic interactions between eukaryotic hosts and photosynthetic microbes have repeatedly shaped ecological productivity and cellular evolution (*8*). The intracellular association between the freshwater ciliate *Paramecium bursaria* and *Chlorella* spp. offers a tractable system for testing whether symbiont-associated metabolic inputs are associated with broad reorganization of host physiology (*9*). Each *P. bursaria* cell harbors hundreds of algal symbionts within host-derived perialgal vacuoles and, under light, can receive photosynthetically fixed carbon while supplying nitrogen, CO₂, and an intracellular niche (*10*). This exchange improves performance under nutrient-limited conditions (*11*). Although nutrient transfer and stress responses in this symbiosis have been studied extensively, a key gap remains regarding how symbionts influence host gene expression and the diel cycle (*1, 12, 13*).

*P. bursaria* cells provide a compelling context for studying how diel temporal programs are organized in a photosymbiotic ciliate. They display distinct behaviors under light versus dark conditions. For instance, they adjust their bacterial ingestion rate, as well as their swimming speed and direction, in response to light intensity and exhibit phototaxis by accumulating in lit regions (*14–16*). In addition, *P. bursaria* exhibits circadian rhythms in mating behavior, photoaccumulation, and swimming velocity (*1, 17, 18*). Some of its cellular activities are also correlated with circadian periods (*19*). However, whether the transcriptional architecture underlying such behavioral rhythms relies on canonical circadian clock genes based on transcription-translation feedback loop (TTFL) components or instead reflects metabolic inputs remains unknown. Moreover, it is unclear to what extent the presence of algal symbionts impacts diel transcriptional organization under LD cycles. Symbiotic Chlorella photosynthetic products have been shown to be required for the expression of the mating-reactivity rhythm in *P. bursaria* (*20, 21*), suggesting that intracellular algae create a day-night metabolic environment that shapes host temporal programs at the behavioral level. *P. bursaria* enables direct comparison of host temporal organization between symbiotic and algae-free aposymbiotic states, allowing us to dissect the contribution of symbiont-associated inputs and the intrinsic host diel scaffold to transcriptional programs.

Diel coordination between hosts and microbial partners has been documented in other systems. In coral-algal symbioses, host gene expression has a pronounced day-night structure (*22, 23*). Comparisons between symbiotic and aposymbiotic coral morphs have indicated that symbiont presence alters both the number and timing of rhythmically expressed host genes (*23*). However, a shared limitation of these photosymbiotic systems is the difficulty of suppressing photosynthesis independently of light. Constant darkness reduces symbiont photosynthetic activity and removes light as the primary entraining cue, making it difficult to attribute temporal reorganization specifically to photosynthesis-linked signals rather than to light removal itself.

Here, we integrate time-series transcriptomics with targeted disruption of photosynthetic electron transport to examine whether and how algal metabolism shapes diel gene expression in the host, *P. bursaria*. By profiling symbiotic and aposymbiotic cells across a 24-h LD cycle and applying rhythmicity analysis, we identify expanded symbiosis-associated temporal modules, primarily driven by endosymbiont photosynthesis. Furthermore, we assess whether regulatory domain-hosting genes are accumulated in rhythmic sets, revealing potential post-translational mechanisms by which photosynthesis reshapes the host diel transcriptional architecture.

## Results

### Endosymbiosis reconfigures the host transcriptome and temporal organization

A stable endosymbiotic relationship requires intimate interactions between the host and endosymbionts. To characterize how these interactions reconfigure host gene expression across the diel cycle, we profiled symbiotic and aposymbiotic *P. bursaria* cells across eight diel time points spanning a 24-h LD cycle (Fig. 1A). Differential expression analysis identified 7,284 genes that were differentially expressed (DEGs) between symbiotic and aposymbiotic cells in at least one time-point comparison (FDR ≤ 0.01, |log₂FC| ≥ 1). A larger proportion of DEGs presented lower expression in symbiotic cells than aposymbiotic ones across the cycle (Fig. 1B; data file S1A-S1D), indicating that the presence of endosymbiotic algae is associated with a broad shift in host state rather than a narrow pathway adjustment. Gaussian mixture model (GMM) clustering of these DEGs resolved five temporal/state clusters (fig. S1A; fig. S2; data file S2A). At baseline, symbiotic cells showed elevated expression of biosynthetic and mitochondria-associated genes (C1), whereas aposymbiotic cells displayed elevated intracellular recycling and lysosome-linked turnover (C2) (fig. S1B; data file S2B). These patterns are consistent with differential resource utilization between symbiotic and algae-free states. Two modules presented state-specific daily timing between symbiotic and aposymbiotic cells. C4 revealed dusk-centered suppression of ciliary and intracellular transport genes in symbiotic cells, implying reduced motility and trafficking capacity as photosynthesis ceases. C5 showed a sharp dawn peak in nutrient-sensing and quality-control programs (MAPK/mTOR/FoxO signaling, autophagy) for symbiotic cells, with the opposite timing in aposymbiotic cells, suggesting that symbiotic cells engage these programs as photosynthetic activity resumes (figs. S3-S4; data files S2B-S2C). To assess the global structure of this transcriptomic divergence, we performed a principal component analysis of genome-wide expression, which confirmed endosymbiont presence as the dominant axis of variation (PC1, 50.7%), with symbiotic samples showing clear time-of-day structure along PC2 (22.0%) and aposymbiotic samples clustering more tightly (Fig. 1C). This global separation is consistent with the transcriptome-wide reconfigurations observed in other photosymbiotic systems (*22–24*). Consequently, our diel sampling design, the PCA-resolved temporal structure, and the state-specific timing in modules C4 and C5 motivated us to perform a systematic evaluation of 24-h rhythmicity across the host transcriptome.

**Fig. 1.**
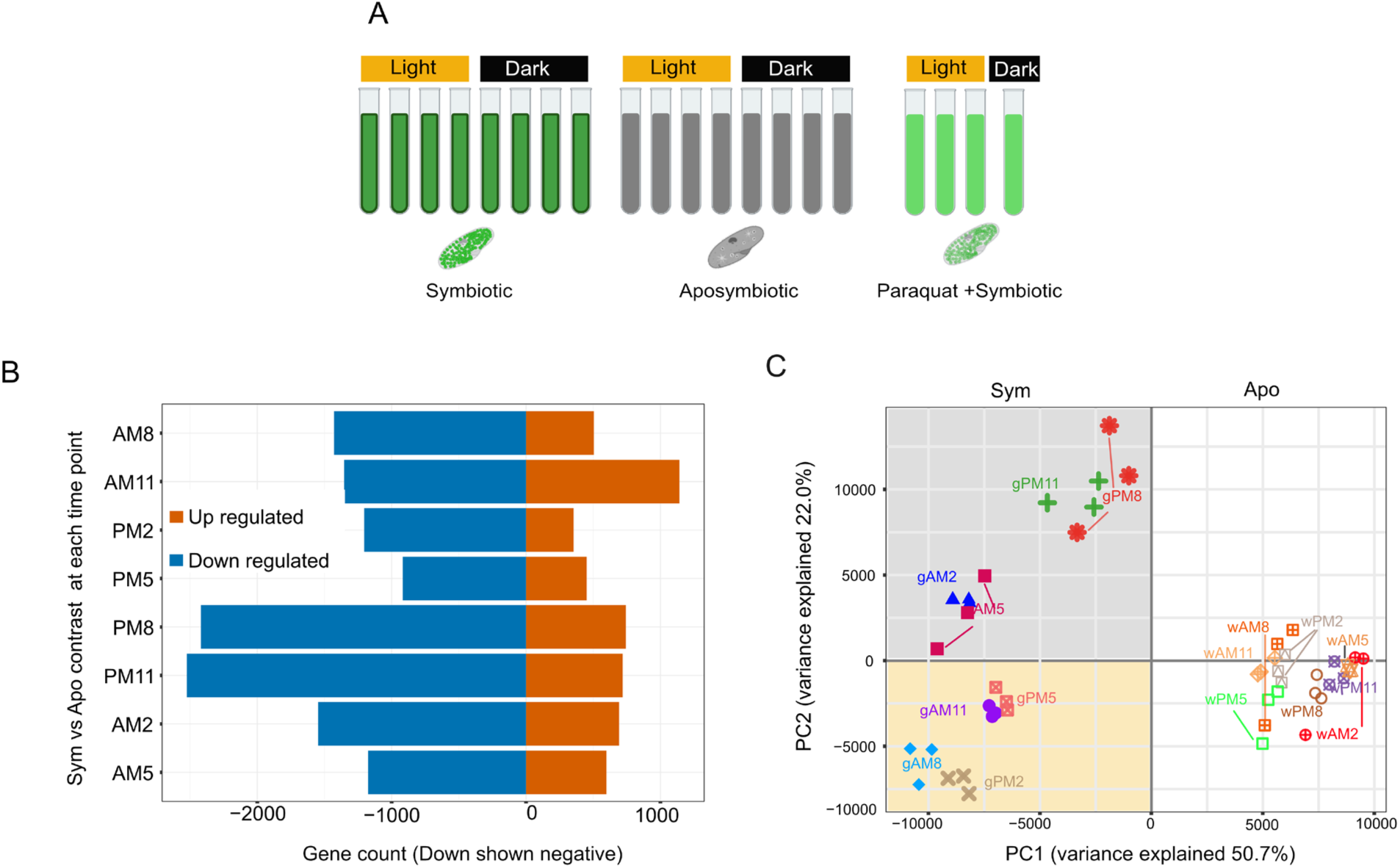
Experimental design, differential expression overview, and global transcriptomic divergence between symbiotic and aposymbiotic *P. bursaria*. (A) Schematic of the experimental design showing symbiotic (Sym), aposymbiotic (Apo), and paraquat-treated (PQ) conditions sampled across eight diel time points under 12-h:12-h LD entrainment. **(B)** Number of upregulated and downregulated differentially expressed genes (DEGs) between symbiotic and aposymbiotic cells at each time point (FDR ≤ 0.01, |log₂FC| ≥ 1). **(C)** Principal component analysis (PCA) of variance-stabilized gene expression (log₂(TPM+1)) from symbiotic and aposymbiotic cells. PC1 (50.7% variance) separates cell states, and PC2 (22.0% variance) reflects temporal structure in symbiotic samples.

### Endosymbiosis amplifies diel rhythmic transcription under LD entrainment

To identify which host genes exhibit statistically supported 24-h periodicity, we applied RAIN (Rhythmicity Analysis Incorporating Nonparametric methods), a nonparametric approach that detects oscillations in time-series data without assuming sinusoidal waveforms (*25*). RAIN identified 3,538 rhythmic genes in symbiotic cells and 705 in aposymbiotic cells (FDR < 0.05), partitioned into 3,300 symbiotic-specific rhythmic genes, 467 aposymbiotic-specific rhythmic genes, and 238 core rhythmic genes shared by both conditions (Fig. 2A; data file S3A-S3B). This seven-fold expansion of rhythmic genes in the symbiotic-specific condition (3,300 genes) compared to the aposymbiotic-specific condition (467 genes) represents the central quantitative signature of symbiont-associated temporal organization under LD entrainment and is evident in the comparative heatmaps of symbiotic-specific, aposymbiotic-specific, and core rhythmic categories (Fig. 2B-2D). Since our time series spans a single 24-h LD cycle, these rhythms reflect diel structure under LD entrainment rather than free-running circadian persistence. Notably, diel transcriptional programs can arise from light-linked metabolic and redox cues even when canonical TTFL circuitry is absent or highly divergent (*26*). Beyond increasing the number of rhythmic genes, symbiosis was also associated with stronger phase ordering across thousands of genes under LD entrainment (Fig. 2B).

**Fig. 2.**
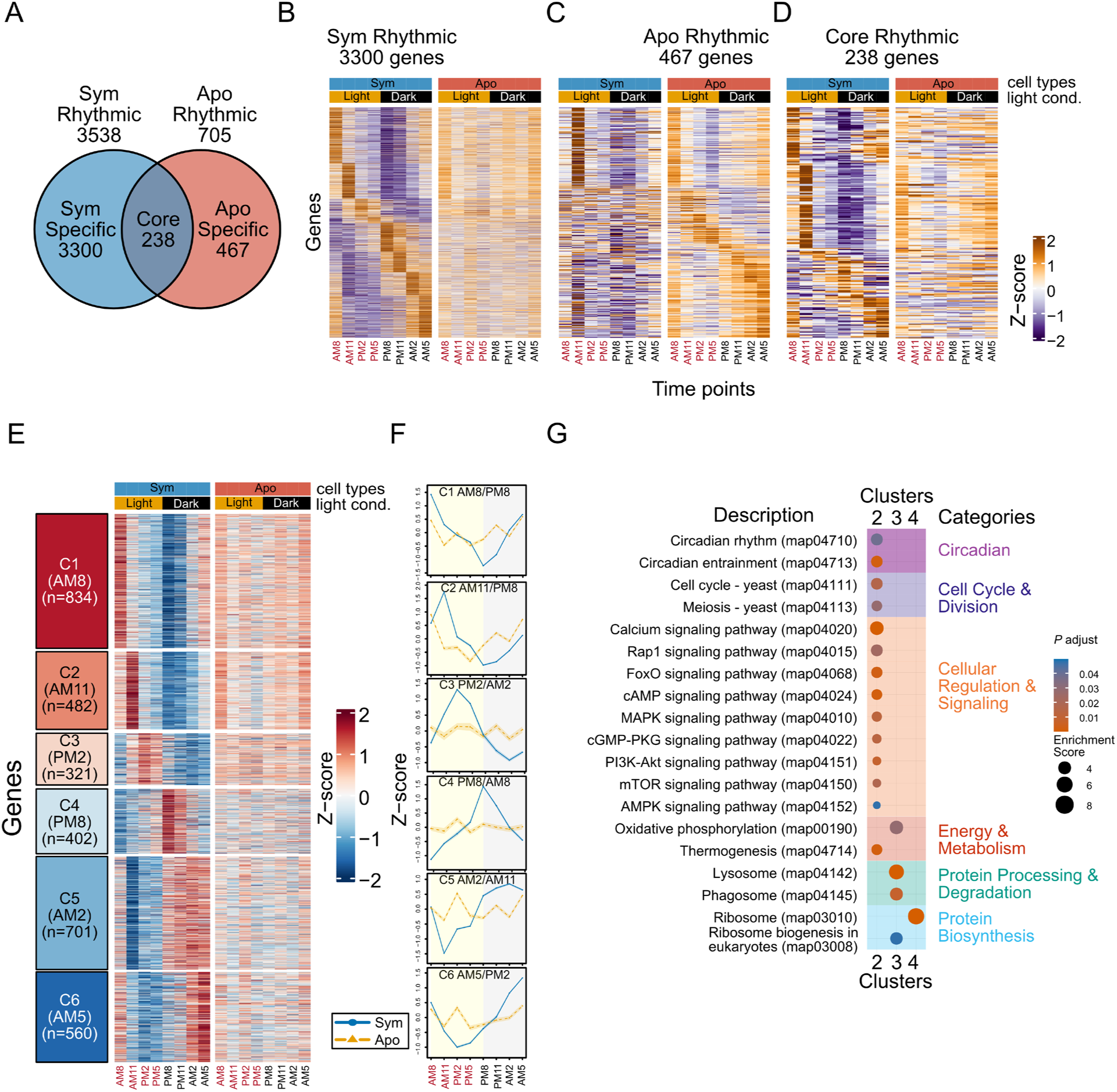
Temporal organization and functional annotation of symbiotic-specific rhythmic gene clusters. (**A**) Venn diagram showing the number of rhythmic genes identified by RAIN (FDR < 0.05) for symbiotic (3,538) and aposymbiotic (705) cells, partitioned into symbiotic-specific (3,300), aposymbiotic-specific (467), and core rhythmic (238) categories. (**B-D**) Heatmaps of Z-scaled log₂(TPM+1) expression across eight diel time points for symbiotic and aposymbiotic cells, shown separately for symbiotic-specific (B), aposymbiotic-specific (C), and core rhythmic (D) gene sets, annotated by cell type and LD condition. (**E**) GMM clustering of 3,300 symbiotic-specific rhythmic genes resolved six temporal clusters (C1-C6), shown as heatmaps with symbiotic and aposymbiotic profiles displayed side by side. (**F**) Mean temporal trajectories for each cluster, displaying Z-scored expression across time points with peak and trough phases indicated for C1-C6. (**G**) KEGG pathway enrichment for clusters C2-C4, the only clusters with statistically significant pathway enrichment, with pathways grouped into functional categories. Dot size denotes enrichment score, and color represents adjusted P value (FDR < 0.05). GO enrichment for all symbiotic-specific rhythmic clusters is provided in fig. S5.

### Symbiosis orchestrates diel activation and subsequent nocturnal remodeling

To determine how the expanded rhythmic program of symbiotic cells is distributed across the day, next we clustered the 3,300 symbiosis-specific rhythmic genes according to their temporal profiles. GMM clustering of these 3,300 genes resolved six temporal clusters spanning the LD cycle in symbiotic cells (Fig. 2E-F; data file S3C). Across these clusters, mean profiles are distributed across distinct phases of the LD schedule (Fig. 2F), supporting an ordered daily program rather than a single dominant peak time. Comparable diel transcriptome structuring linked to symbiosis has been described in cnidarian-algal systems (*22, 23*). In this framework, the cluster series provides an empirical basis for describing how major functional themes are deployed throughout the day in symbiotic cells, as detailed below.

At light onset, the AM8-peaking cluster is dominated by motile cilium and microtubule-based movement and transport, calcium handling, mechanosensory-linked signaling, and proteostasis categories (fig. S5; data file S4A). Light-responsive behaviors in *P. bursaria* have been linked to ciliary control and membrane potential responses (*15*). Notably, symbiotic *P. bursaria* exhibit photoaccumulation that requires symbiont photosynthetic electron transport (*27*) suggesting that algal activity directly shapes host motility. The enrichment of motility and sensory-associated genes specifically at light onset suggests that symbiosis introduces a time-of-day regulatory layer onto these pathways, positioning the host to respond to predictable light-phase changes in symbiont physiology.

Cluster 2 peaks sharply early in the light period, at AM11, and is enriched for a regulatory pulse that combines kinase- and second-messenger signaling with energy and stress sensing, alongside autophagy and proteolysis-related pathways (Fig. 2G; fig. S5; data files S4A-S4B). Rather than interpreting this enrichment in terms of isolated pathways, the co-occurrence of signaling, phosphorylation-centered regulation, and quality-control programs supports a coordinated early-light control window that could gate downstream bioenergetic, biosynthetic, and trafficking investments later in the light phase as the symbiotic program transitions into the light phase under LD entrainment (*28*). Consistent with DEG-based Cluster 5, RAIN-identified Cluster 2 confirms that nutrient-sensing and quality-control programs exhibit formal 24-h periodicity in symbiotic cells, peaking at AM11.

Mid-light, Cluster 3 peaks at PM2 and combines oxidative phosphorylation and ribosome biogenesis with vesicle-associated acidification and degradative trafficking categories, including lysosome and phagosome terms and intracellular pH regulation (Fig. 2G; fig. S5; data files S4A-S4B). This pattern supports a daytime investment phase that couples bioenergetic output and biogenesis with membrane trafficking and acidification systems. The co-enrichment of lysosomal, phagosomal, and pH-regulatory terms at mid-light is consistent with the behavioral observation that symbiotic *P. bursaria* cells increase their rate of bacterial ingestion under light conditions (*14*).

The remaining three clusters span the dark phase and collectively show enrichment for the translation machinery (Cluster 4, peaking at PM8; Fig. 2G; data file S4B), ubiquitin-linked protein turnover (Cluster 5, peaking at AM2; fig. S5; data file S4A), and checkpoint-linked phosphorylation signaling (Cluster 6, rising toward AM5; fig. S5; data file S4A), suggesting a night program oriented toward translational investment, proteome maintenance, and regulatory preparation for the next light transition (*29*).

In summary, this six-cluster organization describes an LD-aligned program in symbiotic cells resolved from a single 24-h cycle. Early-light modules emphasize motility, signaling, and resource allocation-associated regulation; mid-light modules emphasize energy, biogenesis, and vesicle and pH control; dark onset is enriched for translation-capacity annotations; and the night phases emphasize proteome maintenance and regulatory preparation. This progression supports the hypothesis that symbiosis is associated with strengthened phase ordering of host programs under LD entrainment, aligning with photosynthesis-linked daily inputs acting as a major organizing influence (*22, 23, 29*).

### Aposymbiotic cells show lower-contrast LD-entrained rhythmic structure

Clustering of the 467 aposymbiotic-specific rhythmic genes resolved four clusters with lower temporal contrast and fewer enriched functional themes than determined for the symbiotic-specific set under the same LD schedule (Fig. 2C, fig. S6A-B; data file S3D). In early light, the program emphasizes growth and cellular remodeling with accompanying membrane investment, including glycerophospholipid biosynthesis, consistent with a morning-biased focus on cell structure and expansion in algae-free cells (fig. S7A-S7B, data files S4C-S4D). As cells approach dusk and the subsequent dark phase, the aposymbiotic rhythmic program shifts toward oxidative and lipid-centered metabolism, with enrichment for peroxisome-linked functions, fatty-acid degradation and metabolism, and oxidative phosphorylation (fig. S7A-S7B, data files S4C-S4D), consistent with greater engagement of lipid turnover and respiratory catabolism when symbiont-derived photosynthate is absent (*11, 13*). This reduced LD-entrained scaffold is consistent with endogenous time-of-day regulation documented previously for naturally aposymbiotic ciliates. For example, *Paramecium tetraurelia* exhibits a ∼24-h rhythm in cadmium sensitivity that persists in constant darkness, is temperature-compensated, and can be phase-shifted by light pulses (*30*). Together, these patterns support a more constrained, lower-contrast rhythmic program in algae-free cells under LD entrainment, contrasting with the substantially expanded and more strongly phase-structured architecture of the symbiotic state.

### Core rhythmic genes define a shared condition-remodeled diel scaffold

Across the 238 genes identified as rhythmic in both symbiotic and aposymbiotic cells by RAIN (FDR < 0.05, Fig. 2A, 2D, data files S3A and S3B), we explored whether shared rhythmic membership also implies conserved timing. GMM clustering of z-scaled time courses within each condition revealed that the shared set follows broadly similar diel trajectories in both states, but several mean profiles display a shift in phase by approximately one sampling interval (figs. S6C-S6F). For example, C1 peaks near AM8 in both conditions, yet its trough occurs at PM8 in symbiotic cells versus PM5 in aposymbiotic cells (figs. S6C-S6F), revealing that the same scaffold is deployed to different schedules depending on symbiont status. This profile-level similarity does not translate into stable gene-level assignments. Sankey mapping uncovered extensive reshuffling of genes among trajectory classes between conditions (fig. S6G). Of the 238 core genes, 187 (approx. 79 percent) switch cluster membership, including a prominent cohort from a later-morning trajectory in symbiotic cells to a more dawn-weighted trajectory in aposymbiotic cells (fig. S6G, data file S3E). Our inspection of the annotations suggests that the core set spans broadly acting regulators and maintenance functions, including signaling factors, membrane transport and homeostasis components, and redox-buffering enzymes (data file S3E), reinforcing the view of a functionally diverse scaffold rather than a single pathway module.

Peroxiredoxin family members exemplify this remodeling within the shared scaffold. A peroxiredoxin 2C gene shifts from symbiotic C1 to aposymbiotic C2, and a 1-cys peroxiredoxin domain protein shifts from symbiotic C3 to aposymbiotic C4 (data file S3E). Since peroxiredoxin oxidation rhythms have been proposed as a conserved readout of cellular timekeeping (*4*), these reassignments place redox-linked maintenance on different temporal trajectories between symbiotic and aposymbiotic states without changing rhythmic membership. Given that photosynthetic activity by intracellular *Chlorella* is expected to increase host reactive oxygen species, the rhythmic deployment of peroxiredoxin on a symbiosis-specific schedule may reflect the temporal coupling of antioxidant defense with peak photosynthetic output.

Together, these results show that aposymbiotic cells retain an LD-entrained core rhythmic scaffold, whereas the symbiotic state remodels temporal deployment through modest phase shifts and widespread redistribution among trajectory classes. Extensive diel structure under LD cycles can arise either from canonical TTFL circadian systems or from light-driven entrainment of metabolic and redox timing. Consequently, we next tested whether there is evidence for recognizable orthologous TTFL clock components in the *P. bursaria* genome.

### Canonical circadian clock genes lack orthology support in *P. bursaria*

We screened the *P. bursaria* host genome for candidate orthologs of recognizable TTFL components using 42 reference clock proteins from representative model organisms spanning animals, plants, fungi, and cyanobacteria (data file S5A). Orthology searches did not provide convincing evidence for canonical TTFL gene sets in *P. bursaria*. The few significant hits showed domain-level similarity rather than full-length orthology, consistent with shared regulatory domains rather than conserved clock proteins (data file S5A). None of these candidates were rhythmic in either symbiotic or aposymbiotic cells under our LD-entrained time series (RAIN FDR < 0.05, data file S5A), and core animal and fungal TTFL components were not detected (data file S5A). Together, these results provide no support for recognizable orthologs of canonical TTFL clock components in *P. bursaria* under the conditions we examined. Loss of canonical TTFL components has been documented across alveolates, including in apicomplexans that have intrinsic oscillators lacking clock gene orthologs (*31*) and dinoflagellates in which circadian rhythmicity persists with no detectable rhythmic transcript abundance, consistent with predominantly post-translational timing (*32*). Whereas clock gene absence in *Hydra vulgaris* was inferred from genome annotation alone (*33*), our screening combined dual-method orthology detection across 42 reference clock proteins with domain-level evaluation and assessment of rhythmicity for all candidate hits.

### Timing-associated domains are common among symbiotic rhythmic genes

Given the lack of orthology support for canonical TTFL components, we wondered whether symbiosis-associated rhythmicity is associated with gene families involved in post-translational and second-messenger regulation. We curated 1,929 genes carrying timing- and signaling-associated domains, including casein kinases (*34*), EF-hand Ca²⁺-binding proteins (*35*), F-box (*36*) or Kelch (*37*) ubiquitin ligases, WD40 scaffolds (*38*), PAS-domain sensors (*39*), and selected transcription factor families (*40, 41*) (data file S5B). Of these, 239 were rhythmic in at least one condition, with most rhythmic domain genes detected in the symbiotic state (197 symbiotic-specific) compared with far fewer in the aposymbiotic state (29 aposymbiotic-specific), and only 13 shared across both (table S1, data file S5B). For example, 17 casein kinases and 5 F-box proteins were rhythmic almost exclusively under symbiosis, alongside 86 EF-hand Ca²⁺-binding proteins and 77 WD40 scaffold proteins. This distribution suggests that phosphorylation, Ca²⁺-linked control, targeted protein turnover, and scaffold-based complex organization among the prominent regulatory features of the symbiosis-expanded rhythmic program. Casein kinases and F-box ubiquitin ligases form a phosphorylation-degradation axis that controls oscillator protein turnover in fungi and animals (*42*), casein kinase 2 subunits encode classic period-length mutants in *Neurospora* (*43*), and post-translational oscillations involving kinases and redox cycles can sustain rhythmicity independently of transcriptional feedback (*44*). The prevalence of these same domain families among symbiotic rhythmic genes is consistent with diel timing in *P. bursaria* relying on analogous post-translational mechanisms rather than a canonical TTFL. Whether these domain-bearing genes function as active timing regulators remains to be tested.

### Paraquat shifts symbiotic temporal identity toward an aposymbiotic-like architecture

The preceding analyses show that symbiotic cells exhibit an expanded, temporally-structured diel transcriptome relative to aposymbiotic cells, and that shared core rhythmic genes are retimed between states. However, these comparisons do not distinguish whether this temporal organization is more tightly coupled to ongoing photosynthetic activity or to symbiont presence alone. To answer that question, we perturbed photosynthetic electron transport in symbiotic cells by administering paraquat (PQ) while retaining symbionts and examined how host expression trajectories were modified under the same LD schedule.

Photochemical efficiency was comparable across cultures at the onset of treatment (t = 0 h), with a mean Fv/Fm of 0.64 (Fig. 3A, fig. S8A, data file S6A). We selected paraquat for transcriptomic profiling because it produced rapid and sustained suppression of Fv/Fm while preserving host viability and motility, whereas diuron/DCMU caused loss of motility at effective concentrations. To maximize temporal coverage with fewer sampling events, we identified redundant time points using pairwise Pearson correlations of symbiotic versus aposymbiotic log₂FC profiles across all eight time points (fig. S8B). Three correlated pairs emerged, PM2-PM5 (r = 0.79), PM8-PM11 (r = 0.82), and AM2-AM5 (r = 0.79), with AM11 showing no strong correlation with any other time point. We selected one representative from each pair (PM5, PM11), included AM11 as a unique profile, and used AM8 as the light-onset anchor, excluding phases when photosynthetic electron transport is not active (i.e., AM2 and AM5), though retaining PM11 because that system would only just have ceased operating and represents the light-to-dark transition. Under these conditions, PQ treatment was associated with strong suppression of symbiont photochemical efficiency, with abnormal host phenotypes not being observed during sampling, suggesting that the impact of PQ-generated oxidative stress on host cells was minimal, suggesting that the impact of PQ-generated oxidative stress on host cells was limited, thereby supporting its use as a photosynthesis-linked perturbation in our experimental design (Fig. 3A; fig. S8A). However, because PQ was not applied to aposymbiotic cells, we cannot fully distinguish photosynthesis-dependent effects from direct host responses to PQ-generated oxidative stress. Accordingly, the PQ-associated shifts described below should be interpreted with this caveat in mind. PQ-treated symbiotic profiles at the four selected time points were compared with expression profiles from the corresponding time points in the untreated symbiotic and aposymbiotic eight-time-point dataset. After expression-level filtering, PQ comparisons included 3,260 symbiotic-specific rhythmic genes, 466 aposymbiotic-specific rhythmic genes, and all 238 core rhythmic genes.

**Fig. 3.**
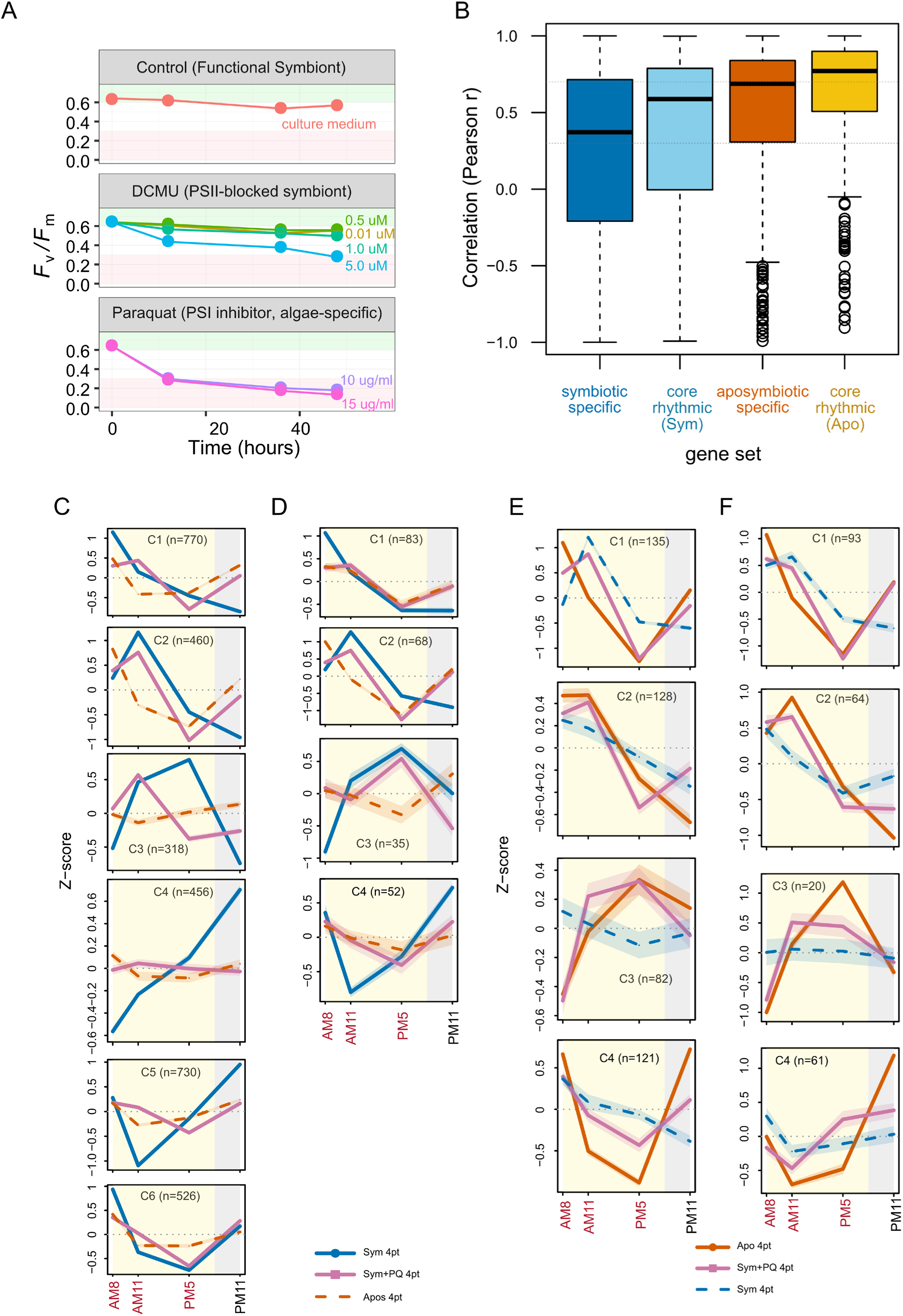
Photochemical verification of PQ treatment and comparison of PQ-treated temporal trajectories against rhythmic and PQ response clusters. (**A**) Maximum quantum yield of photosystem II (Fv/Fm) measured over 48 h in symbiotic *P. bursaria* exposed to control medium, DCMU (PSII inhibitor), or paraquat (PQ, which diverts electrons from PSI). Control cultures maintain stable photochemical efficiency; DCMU reduces Fv/Fm across concentrations of 0.01-5.0 µM; paraquat rapidly suppresses Fv/Fm at concentrations of 10-15 µg/mL. Points represent mean values for each condition. (**B**) Gene-wise Pearson correlation between PQ-treated symbiotic and untreated reference expression profiles across four matched time points (AM8, AM11, PM5, PM11). Boxplots show correlation distributions for Symbiotic-Specific (n = 3,260), Core Rhythmic Sym (n = 238), Aposymbiotic-Specific (n = 466), and Core Rhythmic Apo (n = 238) gene sets. Dashed lines indicate r = 0, 0.3, and 0.7. (**C**) Mean Z-scaled expression trajectories for the six symbiotic-specific rhythmic clusters (C1-C6) across four matched time points, with PQ-treated symbiotic and untreated aposymbiotic profiles overlaid on each cluster. (**D**) Mean trajectories for core rhythmic gene clusters defined from symbiotic temporal profiles, with PQ-treated symbiotic and untreated aposymbiotic profiles overlaid. (**E**) Mean trajectories for four aposymbiotic-specific rhythmic clusters (C1-C4), with PQ-treated symbiotic and untreated symbiotic profiles projected onto each cluster. (**F**) Mean trajectories for core rhythmic gene clusters defined from aposymbiotic temporal profiles, with PQ-treated symbiotic and untreated symbiotic profiles overlaid. Corresponding cluster heatmaps for panels C-F are shown in fig. S9.

To assess if PQ treatment shifted host temporal identity, we correlated the four-time-point expression profile of each gene in PQ-treated symbiotic cells against its matched untreated symbiotic and aposymbiotic reference profiles (see Methods). Among symbiotic-specific rhythmic genes, PQ-treated profiles showed reduced correspondence to the symbiotic reference (median r = 0.372) (Fig. 3B, Table S2, data file S6B). Among genes previously identified as rhythmic only in aposymbiotic cells, PQ-treated symbiotic profiles aligned more closely with the aposymbiotic reference (median r = 0.688) (Fig. 3B, Table S2, data file S6B). Similarly, core rhythmic genes aligned more strongly with the aposymbiotic than with the symbiotic reference (median r = 0.771 versus 0.588) (Fig. 3B, Table S2, data file S6B).

To visualize these shifts at the trajectory level, we projected PQ-treated symbiotic profiles onto the pre-defined rhythmic clusters from untreated cells. Among symbiotic-specific clusters, C1 and C2 retained their respective peak timing under PQ treatment but with reduced amplitude, and both converged toward aposymbiotic trajectories from PM5 onward. C4 and C5 showed near-complete flattening of PQ-treated trajectories toward aposymbiotic-like profiles. C3 and C6 exhibited more complex, cluster-specific trajectories that did not conform to a single directional pattern (Fig. 3C, fig. S9A). When viewed against symbiotic-defined clusters, core rhythmic genes showed clear convergence toward aposymbiotic timing in C1 and C4, with PQ-treated trajectories closely tracking the aposymbiotic reference, whereas C2 showed partial convergence at specific time points (Fig. 3D, fig. S9B). When we projected PQ-treated profiles onto aposymbiotic-specific clusters, all four clusters showed similar trajectories to the aposymbiotic reference, with C2 and C3 showing particularly close alignment (Fig. 3E, fig. S9C). Core rhythmic genes viewed against aposymbiotic-defined clusters presented consistent convergence in C1 through C3, with slightly weaker alignment in C4 (Fig. 3F, fig. S9D). Together, both the correlation analysis and the cluster-level projections suggest that perturbation of symbiont photochemical function biases host temporal organization toward an aposymbiotic-like state, even though symbionts are retained.

### Gene-level classification highlights paraquat-sensitive host systems

We classified gene-level responses using correlation-based thresholds against reference patterns (see Methods): genes with r < 0.3 to the symbiotic reference were classified as rhythm-abolished; those with 0.3 <= r <= 0.7 as partially disrupted; and those with r <= -0.5 as phase-inverted.

Gene-level classification supported the same directional shift (Fig. 4, data file S6C). Among symbiotic-specific rhythmic genes, disruption predominated under the PQ treatment (31.7% rhythm abolished, 27.2% partially disrupted, 14.7% phase inverted) (Fig. 4, data file S6C). In contrast, aposymbiotic-specific and core rhythmic genes showed substantial convergence toward aposymbiotic reference behavior, with core genes showing particularly strong convergence (21.4% converged to the aposymbiotic pattern, 38.7% converged with increased amplitude) (Fig. 4, data file S6C).

**Fig. 4.**
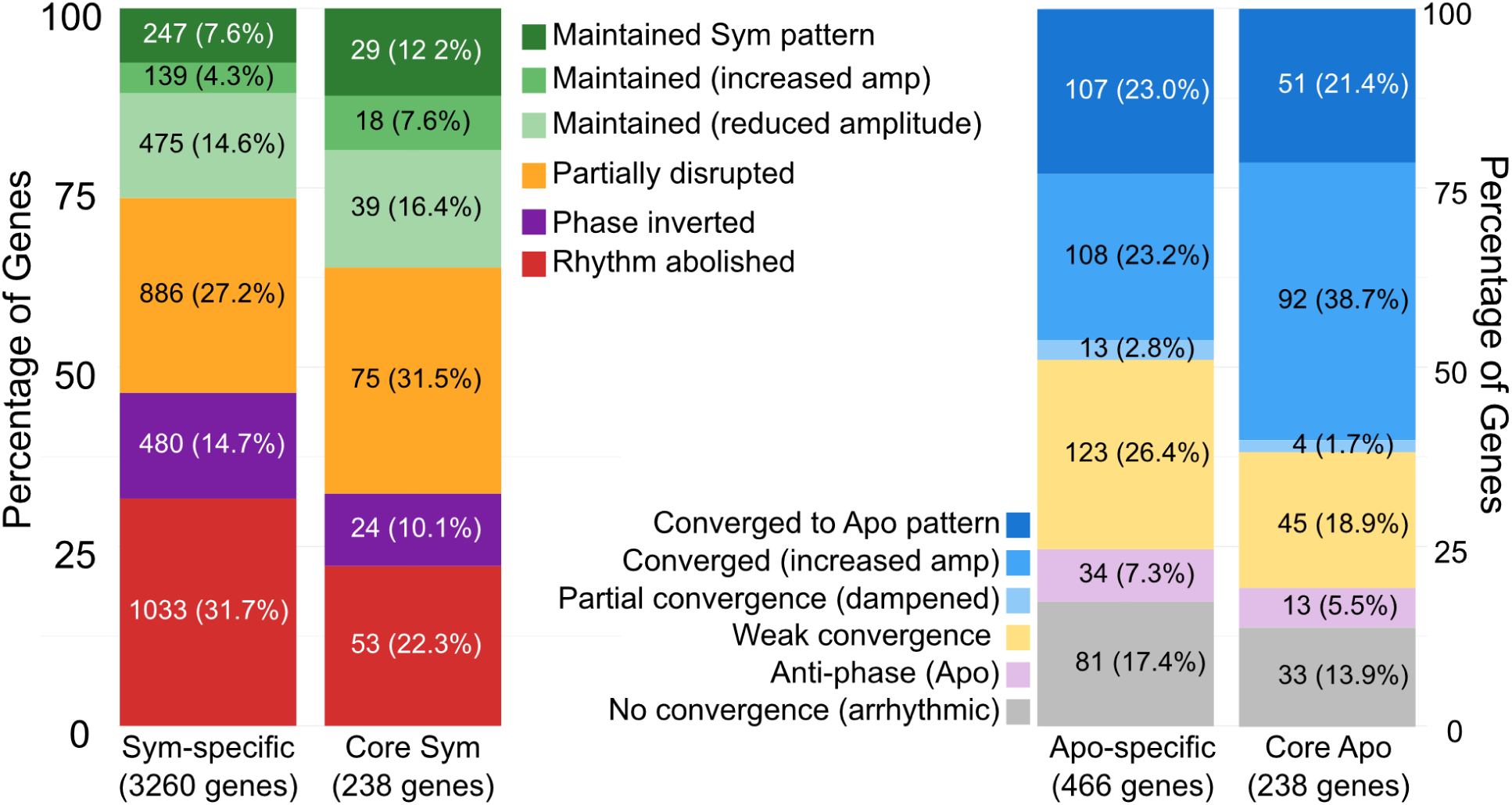
Classification of PQ-induced temporal expression outcomes across symbiotic-specific, aposymbiotic-specific, and core rhythmic gene sets. Stacked bar plots showing the distribution of PQ response categories for symbiotic-specific rhythmic genes (3,260 genes), aposymbiotic-specific rhythmic genes (466 genes), and core rhythmic genes (238 genes). Categories include “maintained symbiotic pattern”, “maintained with increased or reduced amplitude”, “partially disrupted”, “rhythm abolished”, “phase inverted”, “converged to aposymbiotic pattern”, “converged with increased amplitude”, “partial convergence (dampened)”, “weak convergence”, “anti-phase (apo)”, and “no convergence (arrhythmic)”. Percentages represent the proportion of genes assigned to each response type within each rhythmic gene class.

Timing-associated regulatory domains were preferentially disrupted under PQ treatment. Across rhythmic domain genes, 70.8% fell into the disrupted class, 6.8% were fully maintained, and 5.5% showed increased amplitude (tables S3-S4, data file S6D). Several domain families showed especially high disruption rates, including F-box proteins (100%), PAS domain proteins (87.5%), and WD40 repeat proteins (84.2%) (tables S3-S4, data file S6D). These are the same domain families that showed preferential representation among symbiosis-specific rhythmic genes, suggesting a link between the domain-heavy symbiotic timing program to photosynthesis sensitivity under PQ application.

Functional enrichment analysis clarified which host systems are most affected (fig. S10A-S10B, data files S7A-S7B). Categories that maintained symbiotic temporal behavior were enriched for cell-cycle and chromosome-segregation functions, consistent with a broadly deployed growth and resource allocation control program that remains temporally structured under LD even when symbiont photochemical performance is perturbed. Given that *P. bursaria* cells divide every 2.5-3 days rather than daily, these cell-cycle-associated genes likely reflect cumulative cue integration across the diel cycle rather than direct coupling to daily cell division. In contrast, photosynthesis-disrupted categories were enriched for cilium organization, microtubule-based movement, cell projection organization, and regulation of endocytosis, suggesting that the motility and trafficking programs that are prominent in the symbiotic diel state are among the categories most sensitive when symbiont photochemical competence is perturbed (fig. S10A-S10B, data files S7A-S7B) (*45, 46*). In *P. bursaria*, light history and endosymbiont photoadaptation alter ingestion rates and feeding behavior, whereas aposymbiotic cells lack this behavioral plasticity (*14*), providing functional context for the sensitivity of these systems under photosynthesis perturbation. Genes converging toward aposymbiotic patterns were enriched for lipid metabolism and fatty-acid catabolism pathways (fig. S10B, data file S7B), pointing to a shift toward aposymbiotic-like catabolic signatures when photosynthesis-linked symbiont input is compromised.

Taken together, our PQ treatment was associated with a directional shift toward aposymbiotic-like temporal organization across correlation structures, gene-level response classes, regulatory-domain sensitivity, and functional enrichment. Since paraquat perturbs photosynthetic electron transport (*47*), the most conservative interpretation is that intact symbiont photochemical activity is a major contributor to the symbiosis-expanded temporal architecture under LD entrainment, with post-translational regulatory mechanisms showing particular sensitivity to photosynthesis perturbation (*28, 29, 48*).

### Cross-conservation of symbiosis-induced diel expression programs

To test whether the symbiosis-associated diel expression programs identified in *P. bursaria* reflect species-specific regulatory rewiring or conserved responses to endosymbiosis, we compared temporal profiles with those of the distantly related ciliate *Tetrahymena utriculariae* (*49*). The common ancestor of *Paramecium* and *Tetrahymena* diverged approximately 580-640 million years ago (*50*), yet *T. utriculariae* has independently evolved an endosymbiotic relationship with green algae within traps of the carnivorous plant *Utricularia reflexa* (*49*). *T. utriculariae* temporal expression data were obtained from a parallel time-series transcriptomic study (*24*). Cross-species temporal profiles were compared for one-to-one ortholog pairs across six matched zeitgeber time points using Pearson correlation (see Methods; Data file 8, A-B).

First, we assessed if genes that are rhythmic specifically in symbiotic cells display conserved temporal organization in *T. utriculariae*. Symbiotic-specific rhythmic genes exhibited significantly positive cross-species correlations (n = 947, median r = 0.354; *P* adj = 2.34 x 10^-37^), suggesting that despite approximately 580-640 million years of divergence, endosymbiont-associated diel programs are conserved across two independently evolved ciliate-algal endosymbioses (Fig. 5; Data file 8, C-D). In contrast, aposymbiotic-specific rhythmic genes showed negative cross-species correlations (n = 123, median r = -0.305; *P* adj = 1.05 x 10^-3^), consistent with that intrinsic host rhythmic programs have diverged extensively between the two species (Fig. 5; Data file 8, C-D). The contrast between these two distributions was highly significant (*P* adj = 1.57 x 10^-13^), suggesting that it is specifically the endosymbiont-driven component of the host diel program that is shared, rather than the host’s intrinsic timing architecture.

**Fig. 5.**
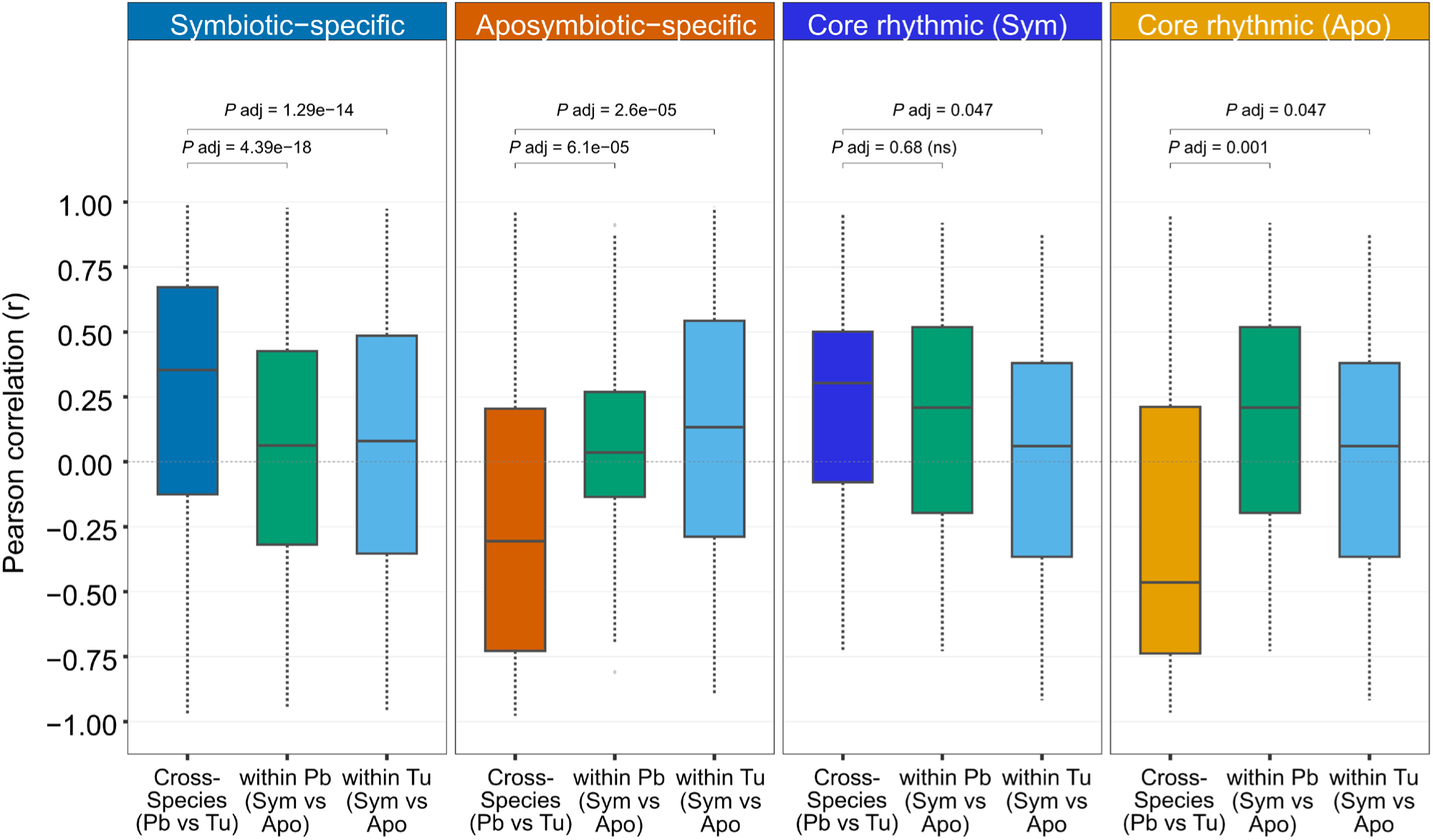
Cross-species conservation of diel expression programs between *P. bursaria* and *T. utriculariae*. Boxplots showing Pearson correlation distributions for 1:1 ortholog pairs across four gene sets. Orange: cross-species correlation between *P. bursaria* and *T. utriculariae* symbiotic temporal profiles. Green: within *P. bursaria* correlation between symbiotic and aposymbiotic profiles. Blue: within *T. utriculariae* correlation between symbiotic and aposymbiotic profiles. Symbiotic-specific genes show significantly positive cross-species correlation (median r = 0.354, *P* adj = 2.34 × 10⁻³⁷), indicating convergent endosymbiont-driven rhythmicity, whereas aposymbiotic-specific genes show negative cross-species correlation (median r = -0.305, *P* adj = 1.05 × 10⁻³). Cross-species correlations were significantly different from within-species correlations for condition-specific gene sets (*P* adj < 6.12 × 10⁻⁵). Dashed line: r = 0. *P* adj (Wilcoxon rank-sum test with Benjamini-Hochberg correction, 14 tests).

Next, we explored if the same pattern holds for genes rhythmic under both conditions. Core rhythmic genes in the symbiotic condition showed positive cross-species correlations (n = 40, median r = 0.304), whereas those in the aposymbiotic condition showed negative correlations (n = 40, median r = -0.465; *P* adj = 6.22 x 10^-4^) (Fig. 5; Data file 8, D). Thus, even among constitutively rhythmic genes, conservation follows the symbiotic rather than the aposymbiotic temporal signature.

To confirm that cross-species correlations reflect endosymbiont-driven convergence rather than general expression similarity, we compared them against within-species symbiotic versus aposymbiotic correlations computed separately for each species (Fig. 5; Data file 8, D). Within-species correlations were near zero for symbiotic-specific genes (within *P. bursaria* median r = 0.063; within *T. utriculariae* median r = 0.080), confirming condition-dependent rather than constitutive temporal profiles. Cross-species correlations were significantly higher than within-species baselines for symbiotic-specific genes (*P* adj = 4.39 x 10^-18^ and 1.29 x 10^-14^) and significantly lower for aposymbiotic-specific genes (*P* adj < 6.12 x 10^-5^), supporting the view that the conserved pattern is specific to the endosymbiont-associated component of the diel program.

Together, these results suggest that photosynthetic endosymbionts are associated with convergent temporal constraints on host gene expression across two independently evolved ciliate-algal partnerships, while intrinsic host diel programs have diverged extensively, pointing to endosymbiont activity acting as a shared organizing input rather than the diel patterns arising from a conserved host timing mechanism.

## Discussion

Photosymbioses couple light-driven symbiont physiology to host cellular state, creating a setting in which daily inputs can structure host time-of-day organization under LD entrainment. In the facultative ciliate-algal system we assessed, symbiotic *P. bursaria* exhibits an expanded and phase-ordered temporal transcriptome relative to algae-free cells, and perturbing photosynthetic electron transport in the endosymbiont is associated with a shift of host expression toward an aposymbiotic-like architecture. Together, these patterns support the view that endosymbiont photochemical function is closely associated with temporal organization under LD entrainment, consistent with photosynthesis-linked inputs providing daily organizing signals that structure host resource allocation across the LD cycle. Comparable day-night structuring linked to symbiont status has been described previously for cnidarian-algal systems (*22, 23*), underscoring how this coupling strategy is recurrent across photosymbiotic partnerships.

A central conceptual finding is that symbiosis amplifies rather than creates host diel temporal structure. The host maintains an LD-aligned diel scaffold even in the absence of symbionts. Symbiont function amplifies and reorganizes this scaffold, expanding it across additional cellular systems and strengthening phase ordering throughout the day. This distinction between baseline temporal organization and symbiosis-driven amplification clarifies how facultative photosymbioses can restructure host temporal organization while retaining the capacity to operate in an algae-free state. One possibility is that metabolic surplus from symbiont photosynthate permits temporal partitioning of costly cellular processes, whereas algae-free cells under resource limitations may maintain more continuous allocations. Testing this hypothesis would require direct metabolic measurements. Alternative mechanisms could also contribute to this temporal structure. Symbiont cell division, which is regulated by host nutritional state and light conditions (*51*), imposes periodic metabolic demands that are independent of photosynthate supply. However, given that *P. bursaria* cells divide every 2.5-3 days rather than daily, the division-associated genes in the diel program we considered likely represent cumulative integration of cues. Light-dependent changes in perialgal vacuole pH (*52*) or symbiont-derived metabolite release (*53*) could also provide daily cues that do not require photosynthetic electron transport per se. Our perturbation of photosynthetic electron transport using paraquat provides evidence against the simplest model of a symbiont presence-based effect, since under our experimental conditions the symbionts remain present, yet temporal structure is disrupted. Nevertheless, disentangling photosynthate supply from other light-dependent symbiont activities will require perturbations that more selectively target specific metabolic outputs.

The absence of recognizable canonical TTFL clock components in *P. bursaria*, combined with the prominence of the post-translational regulatory machinery among symbiosis-specific rhythmic genes, is consistent with LD-aligned temporal organization being shaped by entrainment-linked inputs and downstream regulatory layers, rather than by a recognizable transcript-level TTFL. In contrast to cnidarian-algal systems in which aposymbiotic morphs retain robust endogenous rhythmicity through canonical TTFL clock components (*23*), aposymbiotic *P. bursaria* maintains a reduced LD-entrained rhythmic scaffold despite lacking recognizable orthologs of canonical TTFL components. This pattern is consistent with the retained aposymbiotic program being structured primarily by light-driven entrainment rather than by a recognizable canonical TTFL mechanism. A parallel scenario has been reported for *Hydra vulgaris*, which exhibits LD-dependent diel behavioral and transcriptional rhythms in the absence of canonical clock genes, with those rhythms being lost under constant-light or constant-dark conditions (*33*). In *P. bursaria*, the additional daily metabolic input of endosymbiont photosynthesis may provide a more specific organizing cue than light entrainment alone, with the symbiosis-expanded program shifting toward an aposymbiotic-like organization when endosymbiont photochemical function is perturbed under LD entrainment. Post-translational mechanisms involving kinases, ubiquitin ligases, calcium-binding proteins, and scaffolding domains represent plausible pathways for translating daily photosynthesis-linked variation in physiological state into coordinated protein-level control (*28, 42–44*).

The regulatory breadth of the symbiotic state is further supported by parallel analysis of alternative splicing from the same dataset we used here, which identified 992 differentially spliced introns with splicing enhancers upregulated in symbiotic cells (*54*). Considered together with the transcriptional and post-translational layers we have documented here, that finding supports a multi-tier regulatory architecture in which host gene expression is coordinated across multiple levels in response to symbiont-associated inputs.

Our cross-species comparison indicates that two independently evolved ciliate-algal endosymbioses show partially convergent host diel organization. This pattern is consistent with photosynthetic endosymbionts providing a shared organizing input, potentially through daily photosynthesis-linked metabolic cues that are shared across distinct ciliate-algal associations. The partial nature of this conservation likely reflects differences in both symbiont identity and ecological context. The two systems differ in their algal partners, with *Chlorella* sp. acting as endosymbionts in *P. bursaria* and *Micractinium tetrahymenae* doing so in *T. utriculariae* (*49*). *T. utriculariae* also inhabits the enclosed environment of *Utricularia reflexa* bladder traps rather than open freshwater (*24, 49*). The anti-correlation of aposymbiotic-specific programs between the two species supports the opposite conclusion. In the absence of a shared symbiont-derived organizing input, host diel architectures diverge independently. Species-specific metabolic baselines and endogenous regulatory histories likely shape these distinct programs.

Several limitations define the scope of our interpretations. First, rhythmicity calls are based on a single 24-hour LD cycle, capturing diel structure under entrainment rather than free-running persistence. Although RAIN is designed for short time series, a single cycle cannot distinguish reproducible diel oscillation from transient temporal heterogeneity, so additional sampling cycles would strengthen confidence in the phase assignments we report here. Second, we used four time points for PQ perturbation profiling rather than a full eight-point series. Third, PQ displays pleiotropic chemistry (*47*). Treating aposymbiotic cells with PQ would help distinguish direct host responses to oxidative stress from photosynthesis-dependent effects. Shared responses across both cell types would indicate host-directed PQ sensitivity, while symbiotic-specific responses would reflect perturbation of photosynthetic function.

Complementary perturbations and direct metabolic measurements will be necessary to identify which physiological variables most strongly predict host phase structure. Finally, transcript abundance is an indirect proxy for metabolic flux and post-translational state. Accordingly, integrating proteomics and metabolomics would strengthen mechanistic assignments of the regulatory layers that translate symbiont-linked inputs into coordinated host programs at the protein and metabolic levels.

In summary, our results support a model in which photosynthetic inputs from endosymbionts reorganize and amplify the host diel gene-expression architecture in *P. bursaria* through integrated transcriptional, post-transcriptional, and post-translational regulatory layers. This multi-tier control strategy may represent a recurring feature of facultative photosymbioses, providing a framework for how host programs are coordinated with daily variation in symbiont productivity across changing environmental contexts.

## Materials and Methods

### Strains and culture conditions

*P. bursaria* cells were grown in lettuce medium and fed with *Klebsiella pneumoniae* (NBRC 100048 strain). Cell cultures were maintained at 23 °C with a 12 h:12 h light:dark cycle. Fresh bacteria-containing lettuce medium was added every 3 days, with cells entering early stationary phase 1 day after their last feeding (*55*).

### *P. bursaria* DK2 strain generation

The P. bursaria DK2 strain was generated from parent strains Dd1 and KM2, originally obtained from the Symbiosis Laboratory at Yamaguchi University, following a conjugation protocol described previously (*55*). Aposymbiotic *P. bursaria* Dd1 cells and symbiotic *P. bursaria* KM2 cells were harvested during early stationary phase and washed with modified Dryl’s solution (substituting KH₂PO₄ for NaH₂PO₄·2H₂O). Following a 2-day starvation period, hundreds of cells from both strains were combined to promote conjugation. Individual conjugating pairs were isolated after 5 hours and transferred to separate droplets of modified Dryl’s solution. Upon separation of exconjugants, each cell was isolated to establish clonal lines. The resulting clones were maintained on *K. pneumoniae*-supplemented lettuce medium (2.5% Boston lettuce extract in modified Dryl’s solution) with a minimum 2-day delay before feeding to prevent disruption of macronuclear development. F1 progeny validation was performed using PCR followed by Sanger sequencing of diagnostic genomic regions. The endogenous endosymbiont in the parent strains is *Chlorella variabilis*.

### Aposymbiotic cell generation

Aposymbiotic cells were generated by treating symbiotic cultures with cycloheximide (10 μg/ml) as described previously (*56*). Cells were maintained in lettuce medium until the complete loss of endosymbionts was confirmed by microscopic examination.

### Genome sequencing and annotation

The *P. bursaria* DK2 genome was generated using Illumina paired-end reads from the DK2 strain, with Dd1 haplotype 1 as the reference (*55*). Variant calling was performed using GATK v4.0.3 (*57*) with default parameters. The identified variants were then phased using HAPCUT2 v1.0 (*58*) to generate the two haplotypes. Gene annotation was conducted independently for each haplotype using RNA-seq data from the DK2 strain and protein sequences from *P. caudatum* and *P. tetraurelia*, obtained from ParameciumDB (http://paramecium.i2bc.paris-saclay.fr), following the pipeline described previously (*59*). we assembled the nuclear genome of *P. bursaria* (DK2) into 1,030 gap-free contigs totaling 53.64 Mb (GC 28.80%; N50 = 98,763 bp; BUSCO completeness 85.4%) and annotated 32,695 genes (31,093 transcripts).

### Non-redundant genome annotation

To capture the complete gene repertoire for downstream time-series and diel rhythm analyses, we constructed a non-redundant, haplotype-resolved pangenome for the *P. bursaria* DK2 strain. We first split diploid gene models and sequences into H1/H2 sets. Putative allelic pairs were detected by protein-level reciprocal best hits using MMseqs2 with balanced stringency (≥90% identity and ≥90% bidirectional coverage), then validated by splice-aware genomic mapping with minimap2 to require synteny and ≥90% locus overlap. Genes lacking validated partners underwent rescue mapping of the CDS to the opposite haplotype to distinguish bidirectional rescues (treated as allelic equivalents), unidirectional rescues (retained separately), counterparts at unannotated loci, and unique candidates, while filtering out isoform artifacts (qcov ≤200%). We constructed a bidirectional relationship graph and collapsed transitive chains using connected-component clustering to ensure each locus is represented exactly once. For each component, a single representative gene was chosen using a quality-based hierarchy that prioritized complete annotations/ORFs, consistent haplotype tie-breaking for identical alleles, greater protein length, and RNA-seq splice-junction support. Unpaired genes were retained as unique representatives. The workflow was implemented in Snakemake using standard versions of MMseqs2 (*60*), minimap2 (*61*), and InterProScan v5.40-77 (*62*), producing a non-redundant genome comprising 18,325 CDS, proteins, and gene ID references. For downstream transcriptome analyses, we used this haplotype-resolved non-redundant pangenome. Unless otherwise noted, all RNA-seq quantification, differential expression, rhythmicity, and enrichment analyses reported herein use this non-redundant pangenome (18,325 loci) as the reference gene set.

### Protein function prediction

The protein functions of *P. bursaria* DK2 genes were predicted using multiple tools following a comprehensive approach described previously (*24*). InterProScan v5.40-77 with the --goterms option was used to assign Gene Ontology (GO) terms to proteins. Functional annotation was expanded by inferring protein-protein interactions using DScript v0.2.2 (*63*) with the trained model topsy_turvy_v1.sav. An interaction-probability threshold of 0.92 was chosen to match the density of predicted positives observed for *Saccharomyces cerevisiae* in BioGRID (*64*). For genes lacking functional annotations, GO-term enrichment among predicted interaction partners was tested gene-by-gene using Fisher’s exact test and Bonferroni correction, with statistical significance set at adjusted *P <* 1 × 10⁻¹⁰. Additional annotations were generated using MapMan Mercator4 v7.0 (*65*). For genes still lacking functional assignments, additional predictions were obtained with PANNZER2 (*66*), and STRING v12.0 (*67*). This comprehensive approach substantially increased functional coverage, assigning GO terms to 23,955 genes (Biological Process: 12,565, Molecular Function: 12,256, and Cellular Component: 22,283) among the 31,093 *P. bursaria* DK2 genes.

KEGG pathway annotations were obtained using BlastKOALA and GhostKOALA (*68*) and eggNOG-mapper (*69*). KEGG Orthology (KO) assignments and KEGG pathway mappings were consolidated across all three methods, prioritizing consistent annotations and retaining method-specific assignments when supported by high-confidence matches. To enable organism-specific enrichment analyses, all functional annotations were integrated into a custom annotation database (org.Pbursaria.eg.db) constructed using the AnnotationForge R/Bioconductor package (*70*). This database consolidated GO terms (BP, MF, CC) from InterProScan and KO identifiers, with associated functional descriptions (KONAME), as well as KEGG pathway identifiers (PATH) and pathway names (PATHNAME) derived from the consolidated KEGG annotations.

Gene identifiers (GID) served as primary keys, enabling direct mapping between genes and their associated functional annotations for downstream enrichment analyses.

### Time-series experimental design and sample collection

#### Time-series RNA collection

For diel time-series analysis, both symbiotic and aposymbiotic *P. bursaria* DK2 cells were maintained under a 12 h light/12 h dark cycle with lights turning on at 8:00 AM and off at 8:00 PM. Samples were collected every 3 hours for a total of eight time points: AM8 (8:00 AM), AM11 (11:00 AM), PM2 (2:00 PM), PM5 (5:00 PM), PM8 (8:00 PM), PM11 (11:00 PM), AM2 (2:00 AM), and AM5 (5:00 AM). Three biological replicates were collected for each condition and time point.

#### Paraquat treatment experiments

To suppress symbiont photochemical efficiency in vivo, symbiotic *P. bursaria* cells were treated with one of two photosynthesis inhibitors: paraquat (PQ, 1,1′-dimethyl-4,4′-bipyridinium dichloride), a redox-active compound that diverts electrons from PSI and promotes the generation of reactive oxygen species (ROS) in chloroplasts; or diuron (DCMU, 3-(3,4-dichlorophenyl)-1,1-dimethylurea), a PSII inhibitor that blocks photosynthetic electron transport at the plastoquinone binding site. DCMU was tested at 0.01, 0.5, 1.0, and 5.0 µM, and PQ at 10 and 15 µg/mL. Working concentrations for both compounds were determined empirically using pulse-amplitude-modulated (PAM) fluorometry by measuring the maximum quantum yield of PSII (Fv/Fm) across a dose-response series, while simultaneously monitoring host motility to identify concentrations that suppress photosynthesis without introducing observable host impairment. Based on these criteria, PQ at a concentration of 10 µg/mL was selected for RNA-seq experiments.

Symbiotic *P. bursaria* cells were treated with PQ 24 hours prior to sample collection to ensure strong suppression of photosynthetic function. The selection of time points was guided by unsupervised hierarchical clustering of pairwise Pearson correlations among symbiotic versus aposymbiotic log₂FC profiles at all eight time points, with one representative selected from each major correlation block spanning the light phase and the light-to-dark transition. Three biological replicates were collected for each selected time point.

#### RNA extraction and sequencing analysis

Total RNA (n = 3 biological replicates per condition per time point) was extracted using TRI reagent followed by column cleanup. Cells were collected and lysed in TRI reagent, and RNA was purified following standard phenol-chloroform extraction protocols with subsequent column-based cleanup to ensure high-quality RNA for library preparation (*24*).

Strand-specific RNA libraries were prepared with the SureSelect Strand-Specific RNA Library Preparation Kit for Illumina (Agilent Technologies) following the manufacturer’s instructions. Libraries were sequenced on a NovaSeq 6000 System (Illumina), generating 150 base pair paired-end reads. Reads were quality-checked and trimmed using fastp (*71*), then quantified using Salmon (*72*) with numBootstraps = 200 and gcBias correction against the constructed non-redundant pangenome.

Differential expression analyses were performed using the limma (*73*) framework in R. Transcript abundance estimates (TPM) were filtered to retain genes with TPM > 0.005 in at least 70% of samples, and then log2-transformed with pseudocount addition. Linear models were fitted to the log2-TPM matrix using lmFit, and empirical Bayes moderation (eBayes) (*73*)was applied to stabilize per-gene variance estimates. Genes exhibiting |log₂FC| ≥ 1 and adjusted P ≤ 0.01 (Benjamini-Hochberg correction) were classified as significantly differentially expressed (DEGs).

#### Diel rhythm analysis

Rhythmic gene expression patterns were identified using the RAIN (Rhythmic Analysis Incorporating Nonparametric methods) algorithm (*25*), a nonparametric approach for detecting rhythmic patterns in time-series data. RAIN was selected over alternative rhythm detection methods for three reasons specific to our experimental design. First, unlike parametric methods such as cosinor regression that assume sinusoidal waveforms, and unlike JTK_CYCLE (*74*) that tests against symmetric reference waveforms, RAIN detects rhythms with arbitrary waveform shapes by independently assessing the rising and falling segments of each oscillation using umbrella alternatives (*75*). This property is critical for analyzing *P. bursaria*, in which metabolically-driven diel rhythms are unlikely to conform to the idealized sinusoidal profiles typical of canonical circadian clock outputs. Second, RAIN employs a rank-based nonparametric framework, making it robust to the outliers and non-normal noise distributions commonly encountered in RNA-seq count data. Third, the method is designed to operate on time series comprising as few as 8-10 measurements per period, accommodating our single 24-h cycle sampled at eight time points. RAIN analysis was performed independently on symbiotic and aposymbiotic expression datasets using a 24-hour period corresponding to the LD cycle, with a sampling interval (deltat) of 3 hours matching our experimental design. For each gene, RAIN calculated rhythm parameters including phase and amplitude. Genes with adjusted *P-*values (Benjamini-Hochberg correction; FDR) < 0.05 were classified as rhythmic. This analysis identified rhythmic genes that were partitioned into three categories: symbiotic-specific rhythmic genes (oscillating exclusively in symbiotic cells); aposymbiotic-specific rhythmic genes (oscillating exclusively in aposymbiotic cells); and core rhythmic genes (oscillating in both conditions).

#### Temporal clustering of differentially expressed and rhythmic genes

To identify coherent temporal expression programs, we applied GMM clustering because it assigns genes to clusters with probabilistic confidence and does not assume spherical cluster geometries, making it well-suited for resolving overlapping temporal patterns in time-series data. GMM clustering was implemented through the mclust5 package (*76*). Prior to clustering, biological replicates were averaged for each condition and time point to obtain representative expression profiles, and expression data were z-score normalized (mean-centered and variance-scaled) to enable cross-gene comparison. Three complementary clustering strategies were employed. First, joint clustering was performed on the complete set of DEGs from both symbiotic and aposymbiotic conditions combined, allowing identification of shared and divergent temporal patterns across physiological states. Second, condition-specific clustering was conducted by applying GMM separately to symbiotic and aposymbiotic expression matrices, with profiles from the opposite condition overlaid for comparison to reveal temporal features masked in joint analysis. Third, rhythmic gene clustering was applied independently to each RAIN-derived rhythmic category (symbiotic-specific, aposymbiotic-specific, and core rhythmic gene sets) to resolve their distinct temporal architectures. The optimal number of clusters (k) was determined using Bayesian Information Criterion (BIC), with k values ranging from 2 to 15 tested systematically across all analyses.

#### Functional enrichment analysis of temporal clusters

Functional enrichment analysis was performed for all temporal clusters identified by GMM clustering, including for DEG clusters and rhythmic gene clusters, to characterize the biological processes and pathways associated with specific phases of the diel cycle. All enrichment analyses were conducted using the ClusterProfiler (*77*) package in R with the custom org.Pbursaria.eg.db annotation database.

For GO enrichment, the compareCluster function was applied with enrichGO to test for over-representation of Biological Process (BP), Molecular Function (MF), and Cellular Component (CC) terms independently for each temporal cluster. Statistical significance was assessed using Fisher’s exact test with Benjamini-Hochberg correction for multiple testing, with *P-*value and q-value cutoffs set at 0.05. The complete *P. bursaria* DK2 non-redundant pangenome gene set served as the background universe for enrichment calculations. Revigo (*78*) was used to reduce redundancy among significantly enriched GO terms.

For KEGG pathway enrichment, the compareCluster function was applied with the generic enricher function using custom TERM2GENE and TERM2NAME mappings extracted from org.Pbursaria.eg.db. TERM2GENE tables linked KEGG pathway identifiers (PATH) to gene identifiers (GID), whereas TERM2NAME tables provided pathway descriptions (PATHNAME) for result interpretation. Over-representation testing was performed using Fisher’s exact test with Benjamini-Hochberg correction (*P-*value and *q*-value cutoffs of 0.05), with the non-redundant pangenome serving as the background.

Enrichment scores for each term were calculated as the ratio of gene proportions: (genes in cluster with term / total genes in cluster) divided by (genes in background with term / total genes in background). For visualization, enrichment results were filtered to retain terms with enrichment scores ≥ 2.0, ensuring that only substantially enriched terms were displayed.

Enrichment results were visualized as dot plots with point size representing enrichment score and point color indicating adjusted *P-*value using a colorblind-friendly gradient (orange for low/significant *P-*values, blue for high/less significant *P-*values). Results were exported to multi-Excel workbooks containing all enrichment results, significantly enriched terms (adjusted *P* ≤ 0.05), and simplified GO results where applicable.

#### Rhythm visualization

Temporal expression patterns of rhythmic gene clusters were visualized using heatmaps displaying z-score normalized expression values across all eight time points (AM8, AM11, PM2, PM5, PM8, PM11, AM2, AM5). Genes were ordered by cluster assignment, and time points were ordered chronologically to reveal phase relationships. Cross-condition heatmaps were generated to compare temporal patterns between symbiotic and aposymbiotic states, with expression profiles from both conditions displayed to assess rhythm amplitude, phase, and coherence differences.

#### Transcription-translation feedback loop (TTFL) clock presence-absence screen

Orthology searches for canonical TTFL circadian clock genes were performed against reference sequences from animals (CLOCK, BMAL1, PER, CRY, NR1D, RORA, RORB), plants (CCA1, LHY, RVE, APRR), fungi (FRQ, WC-1, WC-2), and cyanobacteria (KaiA/B/C). Independent BLASTp (*79*) and DIAMOND (*80*) searches of the *P. bursaria* DK2 proteome were performed using an e-value threshold of 1e-05. BLASTp searches used max_hsps 1, comp_based_stats 2, and max_target_seqs 500. DIAMOND searches used --ultra-sensitive mode with equivalent parameters. Best hits were determined by lowest e-value per query and reciprocal top hits. For each hit, query coverage (percentage of TTFL reference protein aligned) and subject coverage (percentage of *P. bursaria* protein aligned) were calculated to distinguish full-length orthologs from domain-level matches. Matches were retained only if both BLASTp and DIAMOND returned results. Single-method hits were excluded to prioritize specificity. Supporting domain evidence from InterProScan (*62*) annotations was used to classify matches (bHLH+PAS for CLOCK/BMAL, cryptochrome-specific signatures for CRY, response regulator domains for PRR/NR1D, Zn2Cys6 domains for fungal WC-like factors). Alignment quality was evaluated by requiring e-value ≤ 1e-10 for orthologous relationships. Matches meeting the initial search threshold but exhibiting <30% query coverage were classified as domain-level homologs rather than true orthologs.

#### Diel timing-associated domain analysis

To characterize the post-translational regulatory machinery supporting diel temporal gene expression, we identified genes from the *P. bursaria* DK2 proteome bearing domains associated with diel timing mechanisms across eukaryotes, including casein kinases (*34*), EF-hand calcium-binding proteins (*35*), F-box proteins (*36*), Kelch-repeat proteins (*37*), WD40-repeat proteins (*38*), PAS domains (*39*), MYB transcription factors (*40*), and BTB/POZ domain proteins (*41*). Domain annotations were obtained from InterProScan analysis of the *P. bursaria* DK2 proteome. We then cross-referenced this domain-bearing gene set with the RAIN-identified rhythmic genes (FDR < 0.05) to quantify the extent to which diel timing-associated protein families exhibited oscillations under the symbiotic versus aposymbiotic conditions. Proteins bearing multiple timing-associated domain families were assigned combined annotations rather than being reduced to a single category, and all downstream counts were based on those combined annotations. Rhythmic status per condition (symbiotic, aposymbiotic, both), phase assignments, and *P*-values from RAIN analysis were used to derive category-level counts.

#### Paraquat-induced rhythm interference analysis

Since PQ-treated samples were collected at only four time points, which is insufficient for RAIN-based rhythm detection (requiring ≥6-8 time points), we developed a pattern-correlation approach to assess PQ-associated changes in temporal patterns relative to the reference states. For each rhythmic gene identified by RAIN under the symbiotic or aposymbiotic condition, we extracted expression values at the four matching time points from the eight time point reference dataset (AM8, AM11, PM5, PM11). Then, we calculated the Pearson correlation coefficient between the four time point reference pattern and the four time point PQ-treatment pattern for each gene, along with the associated *P-*value and 95% confidence interval from cor.test. Per-gene *P-*values were corrected for multiple testing using the Benjamini-Hochberg procedure across all tested genes. Amplitude was calculated as the standard deviation of expression values across the four time points for both the reference pattern (amplitude_ref) and the PQ-treatment pattern (amplitude_pq). The amplitude ratio was computed as amplitude_pq divided by amplitude_ref, providing a quantitative measure of how PQ treatment affected oscillation strength relative to the reference pattern. To compare correlation distributions across gene sets, one-sample Wilcoxon signed-rank tests were used to assess if the median correlation of each gene set differed from zero, and two-sample Wilcoxon rank-sum tests were deployed to compare correlation distributions between the symbiotic-specific and aposymbiotic-specific gene sets, and between the core rhythmic under symbiosis and core rhythmic under aposymbiosis gene sets. All six *P-*values were corrected using the Benjamini-Hochberg procedure. We note that Pearson correlations computed from four data points have limited statistical power at the individual gene level (df = 2).

Accordingly, we have interpreted PQ-associated shifts primarily at the aggregate distribution level using the Wilcoxon tests, rather than assigning high confidence to individual gene classifications.

#### Paraquat-induced interference frameworks (Disruption and Convergence)

To clarify the biological meaning of rhythm changes under PQ treatment, genes were assigned interpretation categories using two complementary frameworks based on Pearson correlation thresholds (r > 0.7, high; 0.30-0.70, moderate; r < 0.30, negligible) following established benchmarks for biological data (*81*). The Disruption Framework was applied to symbiotic-specific rhythmic genes and core rhythmic genes, and they were analyzed against symbiotic references to characterize how symbiotic rhythms change under PQ treatment. Within this framework, genes were classified as: maintained symbiotic pattern (PQ-insensitive patterns with r > 0.7 and 0.75 ≤ amplitude ratio ≤ 1.25); maintained with increased amplitude (r > 0.7 and ratio > 1.25); maintained with reduced amplitude (r > 0.7 and ratio < 0.75); partially disrupted (0.3 ≤ r ≤ 0.7); rhythm abolished (r < 0.3); or phase inverted (r ≤ -0.5).

The Convergence Framework was applied to aposymbiotic-specific rhythmic genes and core rhythmic genes, and the results were analyzed against aposymbiotic references to determine whether genes adopt aposymbiotic-like expression patterns under PQ treatment. Within this framework, genes were classified as: converged to aposymbiotic pattern (strong convergence to the aposymbiotic pattern with r > 0.7 and 0.75 ≤ amplitude ratio ≤ 1.25); converged with increased amplitude (r > 0.7 and ratio > 1.25); partial convergence with dampening (r > 0.7 and ratio < 0.75); weak convergence (0.3 ≤ r ≤ 0.7); no convergence; arrhythmic (r < 0.3); or anti-phase to aposymbiotic (r ≤ -0.5). This dual framework enabled differentiation between PQ-insensitive pattern maintenance and PQ-associated pattern disruption or convergence to aposymbiotic states. Core rhythmic genes (n=238) were analyzed twice: once against symbiotic four-time-point subsets (Disruption Framework) and once against aposymbiotic four-time-point subsets (Convergence Framework), to comprehensively characterize their responses to PQ treatment.

#### Cross-species conservation of diel expression patterns

Orthologous gene pairs between *P. bursaria* and *T. utriculariae* were identified using a combination of three complementary orthology inference methods: Reciprocal Best BLAST Hits (RBH) with BLASTp (*79, 82*) (e-value threshold of 1e-5, ≥30% identity); OrthoFinder v2.5.4 (*83*); and OrthoMCL single-copy clusters (*84*). Non-redundant collation of all pairs across methods was retained to maximize coverage. For the cross-species correlation analysis, only strict one-to-one ortholog pairs were used, yielding 4,437 unique *P. bursaria*-*T. utriculariae* pairs supported by one or more methods.

To assess conservation of diel expression programs, Z-score normalized temporal expression profiles of *P. bursaria* genes and their *T. utriculariae* orthologs were compared using Pearson correlation across six matched zeitgeber time points (AM8, AM11, PM2, PM5, PM8, PM11). Only *P. bursaria* genes classified as being diel-rhythmic by RAIN (Benjamini-Hochberg-adjusted *P* < 0.05) with a one-to-one *T. utriculariae* ortholog were included. Ortholog pairs were assigned to the Symbiotic-Specific, Aposymbiotic-Specific, Core Rhythmic (Sym), or Core Rhythmic (Apo) gene sets based on *P. bursaria* rhythmicity classification.

To contextualize cross-species correlations, within-species correlations between symbiotic and aposymbiotic temporal profiles were computed separately for *P. bursaria* and *T. utriculariae*. For each gene, log_2_(TPM + 1)-transformed expression values were averaged across biological replicates at each of the six shared time points to generate per-condition temporal profiles, and the Pearson correlation between symbiotic and aposymbiotic profiles was computed.

Statistical significance of correlation distributions was assessed using Wilcoxon tests.

One-sample Wilcoxon signed-rank tests were used to evaluate if the median cross-species correlation of each gene set differed from zero. Two-sample Wilcoxon rank-sum tests were deployed to compare cross-species correlation distributions between Symbiotic-Specific and Aposymbiotic-Specific gene sets, and between Core Rhythmic (Sym) and Core Rhythmic (Apo) gene sets. Cross-species correlations were also compared against within *P. bursaria* and within *T. utriculariae* correlations for each gene set using Wilcoxon rank-sum tests. All *P*-values were corrected together using the Benjamini-Hochberg procedure.

#### Statistical Analysis

Statistical tests used in this study are specified in the corresponding Methods subsections and figure legends. Unless otherwise noted, all reported *P* values were corrected for multiple comparisons using the Benjamini-Hochberg procedure. Analyses were performed in R (v4.3.1) using base statistical functions and the RAIN package for rhythmicity detection.

## Acknowledgments

We thank the members of the Leu lab for their helpful discussions and comments on the manuscript. We also thank John O’Brien for manuscript editing and the IMB Genomics Core for experimental assistance.

## Funding

This work was supported by Academia Sinica of Taiwan (grant no. AS-IA-110-L01 and AS-GCS-113-L03).

This work was supported by the National Science and Technology Council of Taiwan (NSTC 113-2326-B-001-002).

MMK was supported by an NSTC postdoctoral fellowship (NSTC 113-2811-B-001-065).

## Author contributions

Conceptualization: MMK, JYL

Methodology: MMK, JYL

Investigation: MMK, YHC

Formal analysis: MMK

Data curation: MMK, YHC, CFJL, CLY

Software: MMK

Visualization: MMK Resources: CK, JYL

Supervision: JYL

Writing-original draft: MMK, JYL

Writing-review & editing: MMK, YHC, CFJL, CLY, CK, JYL

## Competing interests

The authors declare that they have no competing interests.

## Data and materials availability

The transcriptomic data, *P. bursaria* DK2 Oxford Nanopore genomic reads, and diploid gene annotation generated in this study are available from NCBI under BioProject PRJNA1279681. The *P. bursaria* DK2 haplotype-resolved genome assemblies (GenBank: GCA_043974915.1, GCA_043974975.1) are available under BioProjects PRJNA556774. All data needed to evaluate the conclusions in the paper are present in the paper and/or the Supplementary Materials.

## Supplementary Materials

**Fig. S1.**
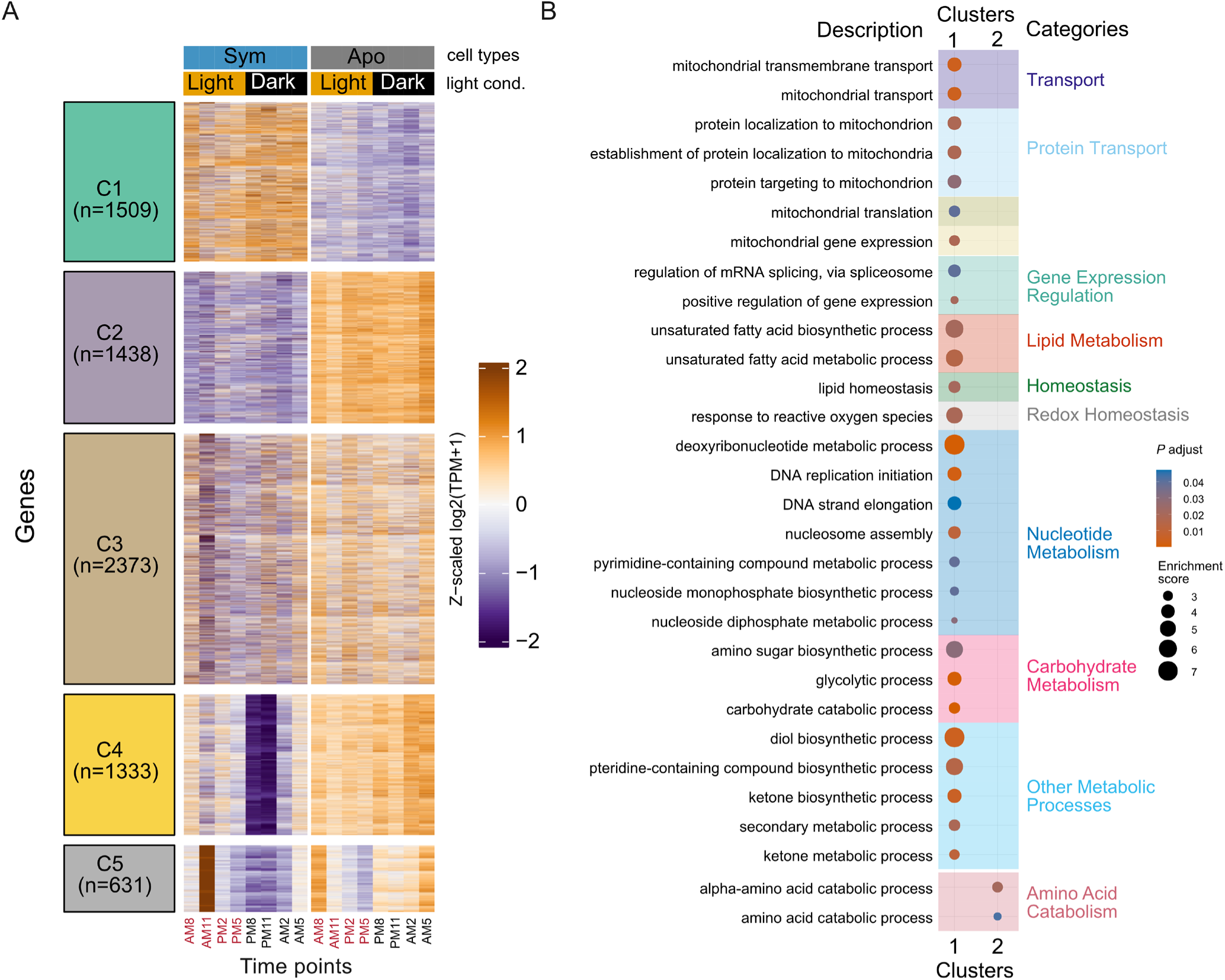
Temporal GMM clustering of differentially expressed genes and GO biological process enrichment of Clusters 1 and 2 in *P. bursaria*. **(A)** Gaussian mixture model (GMM) clustering of 7,284 differentially expressed genes (FDR < 0.01; |log₂FC| > 1) across the combined dataset. Five clusters (C1-C5) are shown with Z-scaled log₂(TPM+1) expression profiles across all timepoints for both conditions, annotated by condition and light-dark status; corresponding mean temporal trajectories for each cluster are provided in fig. S2. (**B**) Gene Ontology (GO) biological process enrichment for C1 and C2, with enriched terms grouped into functional categories. Dot size indicates enrichment score, and color represents adjusted *P* value; GO enrichment for Clusters 4 and 5 is in fig. S3; no significant enrichment was detected for Cluster 3.

**Fig. S2.**
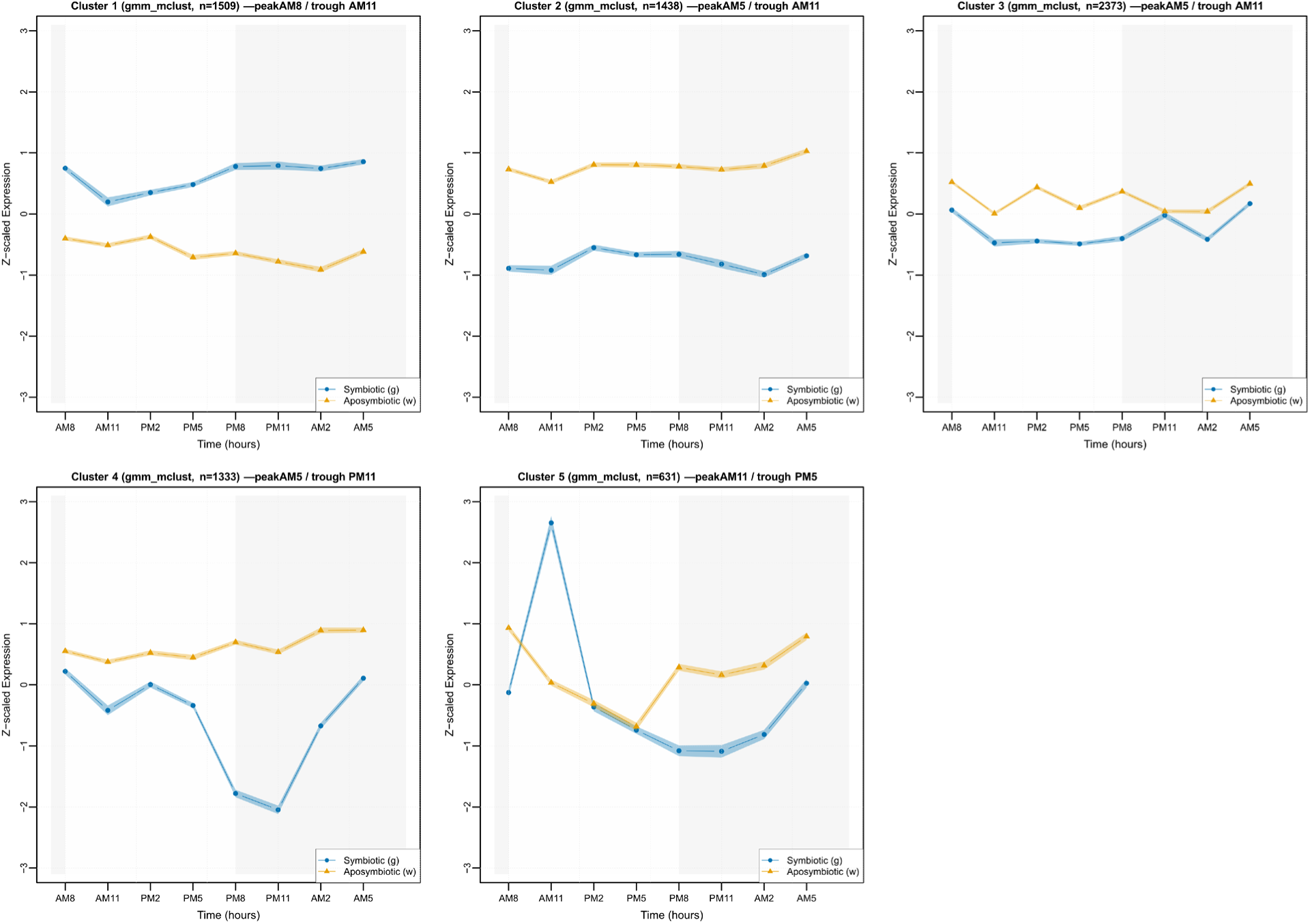
Mean temporal expression profiles for the five joint GMM clusters corresponding to heatmaps in Fig. S1A. Mean Z-scaled log₂(TPM+1) expression trajectories for genes assigned to Cluster 1 (n = 1,509), Cluster 2 (n = 1,438), Cluster 3 (n = 2,373), Cluster 4 (n = 1,333), and Cluster 5 (n = 631) derived from Gaussian mixture model clustering of the joint symbiotic and aposymbiotic dataset. These trajectories correspond to the cluster heatmaps shown in **Fig. S1A**. Each panel shows the average expression pattern across eight diel time points (AM8, AM11, PM2, PM5, PM8, PM11, AM2, AM5) for symbiotic (g) and aposymbiotic (w) cells, plotted in chronological order.

**Fig. S3.**
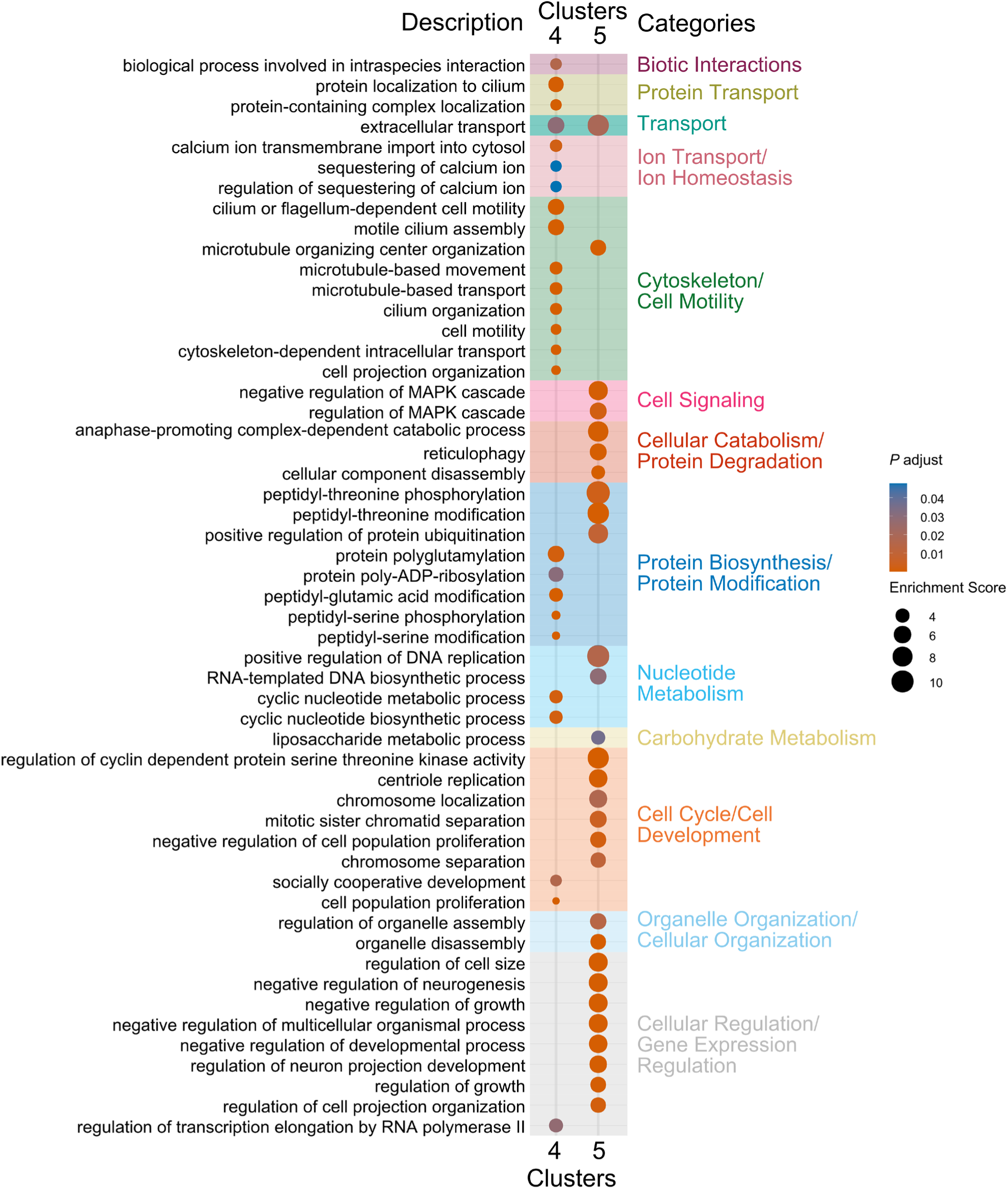
GO biological process enrichment for Clusters 4 and 5 corresponding to Fig. S1B. Gene Ontology biological process enrichment for genes assigned to Cluster 4 and Cluster 5 from the joint GMM clustering shown in fig. S1A, complementing the enrichment analysis for Clusters 1 and 2 in fig. S1B. Enriched terms (adjusted *P* < 0.05) are displayed and grouped into manually curated functional categories, including biotic interactions, ion transport and homeostasis, protein transport, cytoskeleton and cell motility, cellular organization, cell signaling, cellular catabolism and protein degradation, nucleotide metabolism, carbohydrate metabolism, cell cycle and development, and protein biosynthesis and modification. Dot size indicates enrichment score, and color denotes adjusted *P* value.

**Fig. S4.**
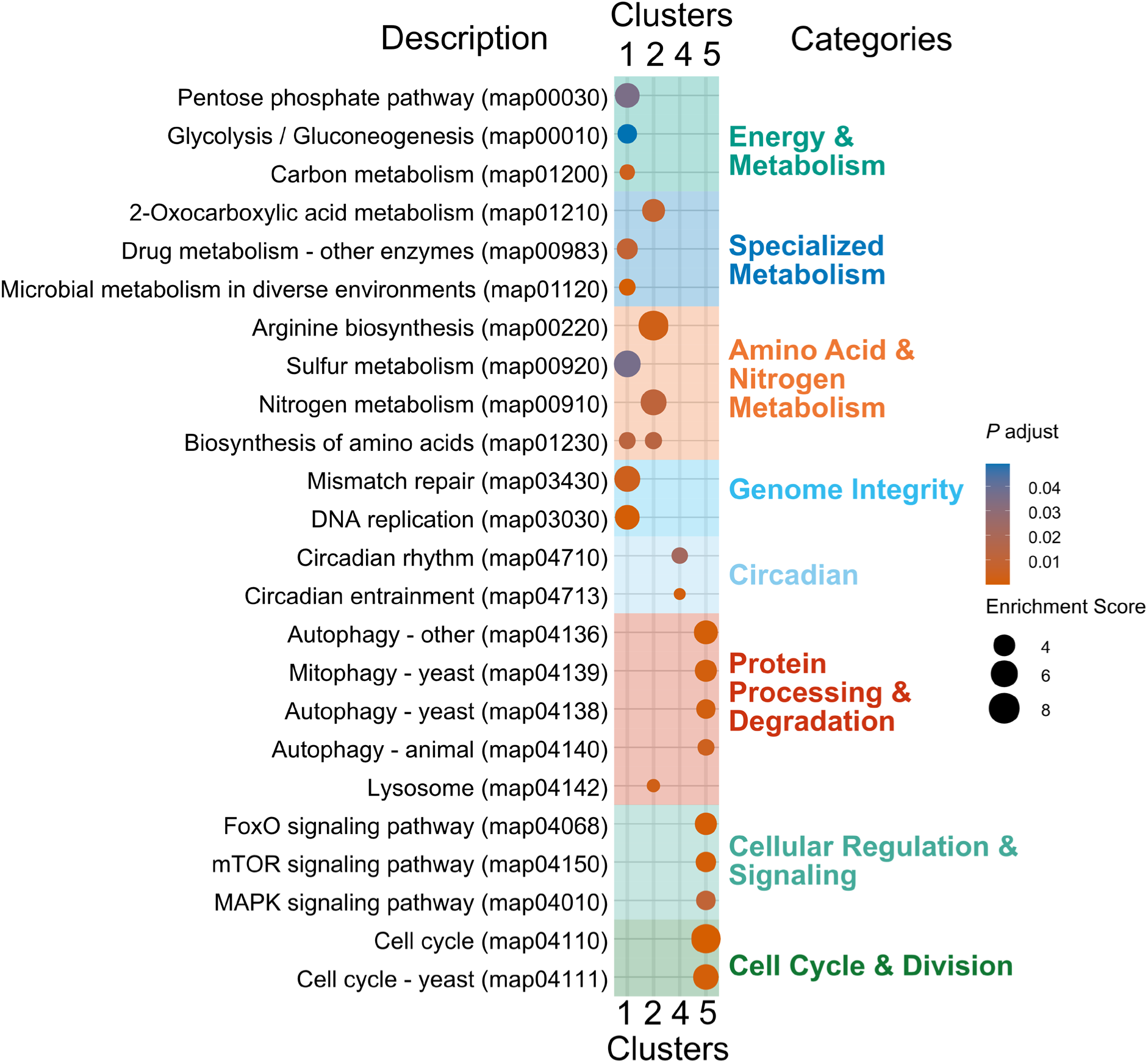
KEGG pathway enrichment for GMM clusters corresponding to fig. S1A. KEGG pathway enrichment for genes assigned to Clusters 1, 2, 4, and 5 from the joint GMM clustering shown in fig. S1A, complementing the GO biological process enrichments presented in fig. S1B and figs. S3. Enriched KEGG pathways (adjusted *P* < 0.05) are displayed with functional groupings, including energy and carbon metabolism, specialized metabolism, amino acid and nitrogen metabolism, genome integrity, circadian pathways, protein processing and degradation, cofactor metabolism, cellular organization, cellular regulation and signaling, cell cycle and division, and immune and defense processes. Dot size indicates enrichment score, and color represents adjusted *P* value.

**Fig. S5.**
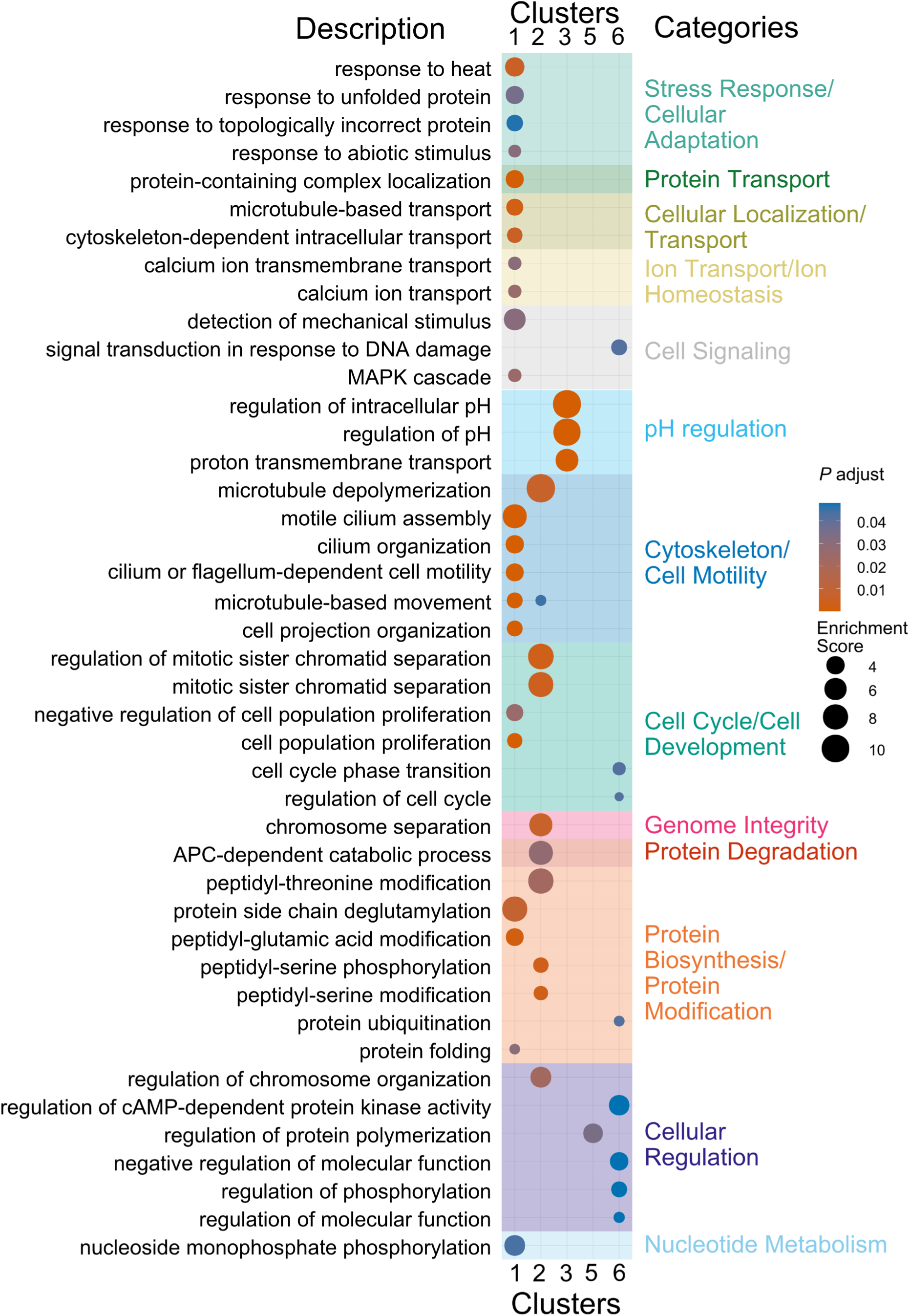
GO biological process enrichment for the symbiotic-specific rhythmic clusters shown in Fig. 2E-F. Gene Ontology biological process enrichment (adjusted *P* < 0.05) for the six symbiotic-specific rhythmic clusters (C1-C6) derived from Gaussian mixture model clustering of 3,300 rhythmic genes. Enriched terms are grouped into functional categories, including stress response and cellular adaptation, protein transport, cellular localization and intracellular transport, ion transport and homeostasis, cell signaling, pH regulation, cytoskeleton and cell motility, cell cycle and cell development, genome integrity and protein degradation, protein biosynthesis and post-translational modification, and nucleotide metabolism. Dot size represents enrichment score, and color denotes adjusted *P* value. These enrichments provide functional characterization of the temporal gene modules shown in Fig. 2E-F.

**Fig. S6.**
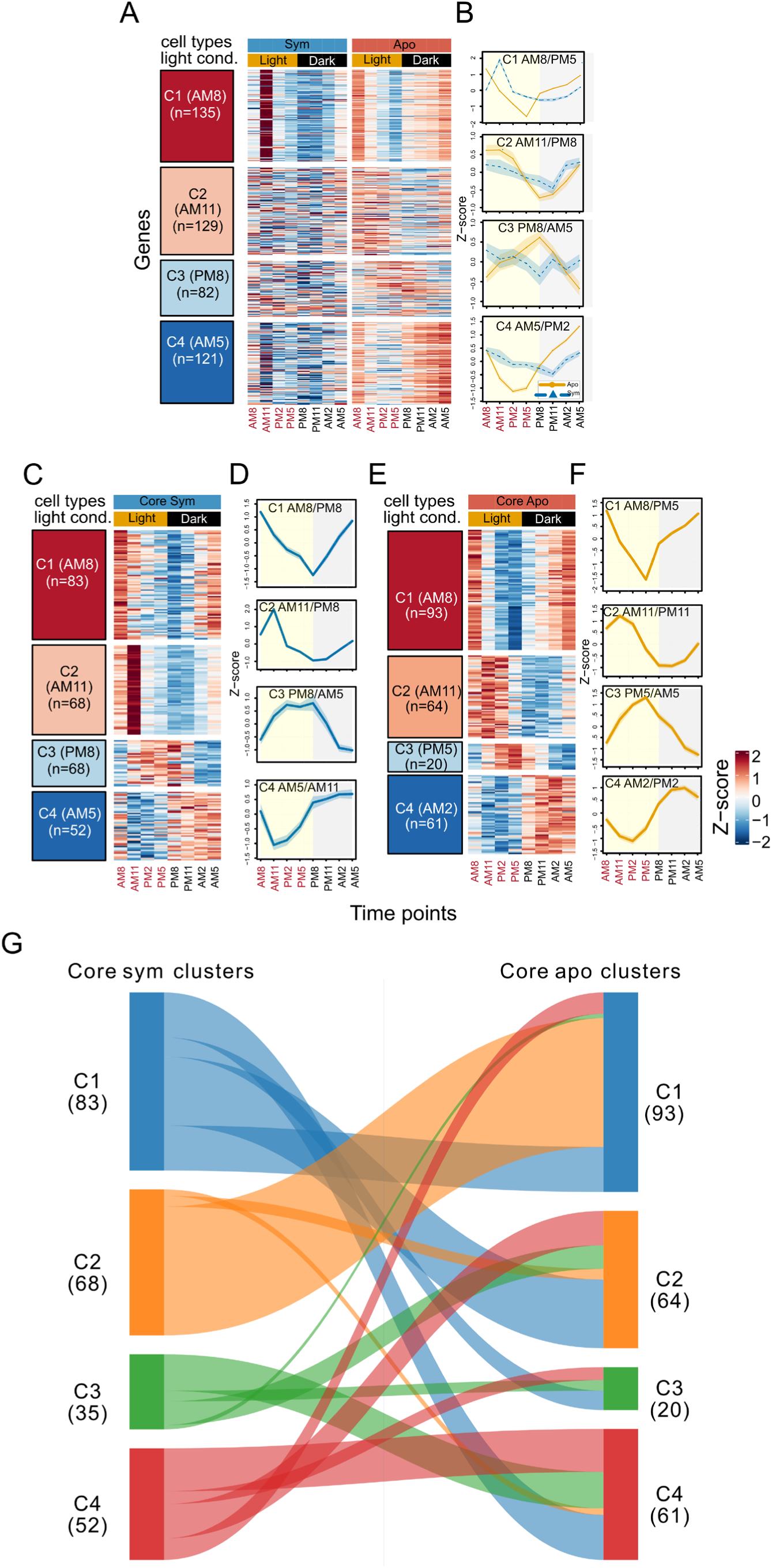
Clustering of aposymbiotic-specific rhythmic genes and condition-specific clustering of the core rhythmic scaffold supporting Fig. 2A, C, and D. (**A**) Heatmap of Z-scored expression for the 467 aposymbiotic-specific rhythmic genes clustered into four temporal groups (C1-C4; gene counts shown) across eight diel timepoints (AM8, AM11, PM2, PM5, PM8, PM11, AM2, AM5). Expression is shown for both symbiotic and aposymbiotic samples, with column annotations indicating cell type and light-dark condition. (**B**) Mean Z-scored temporal trajectories for the four aposymbiotic-specific clusters in panel (A), with cluster labels indicating the approximate peak/trough timing (C1 AM8/PM5, C2 AM11/PM8, C3 PM8/AM5, C4 AM5/PM2). (**C-D**) Gaussian mixture model clustering of the 238 core rhythmic genes performed within symbiotic cells (“Core Sym”), shown as a cluster heatmap (**C**) and corresponding mean trajectories (**D**) for four clusters (C1-C4; gene counts shown; labels indicate approximate peak/trough timing: C1 AM8/PM8, C2 AM11/PM8, C3 PM8/AM5, C4 AM5/AM11). (**E-F**) Gaussian mixture model clustering of the same 238 core rhythmic genes performed within aposymbiotic cells (“Core Apo”), shown as a cluster heatmap (**E**) and mean trajectories (**F**) for four clusters (C1-C4; gene counts shown; labels indicate approximate peak/trough timing: C1 AM8/PM5, C2 AM11/PM11, C3 PM5/AM5, C4 AM2/PM2). (**G**) Sankey/alluvial mapping linking core-cluster assignments between the symbiotic and aposymbiotic clustering results, illustrating redistribution of the same core genes among condition-specific temporal trajectory classes.

**Fig S7.**
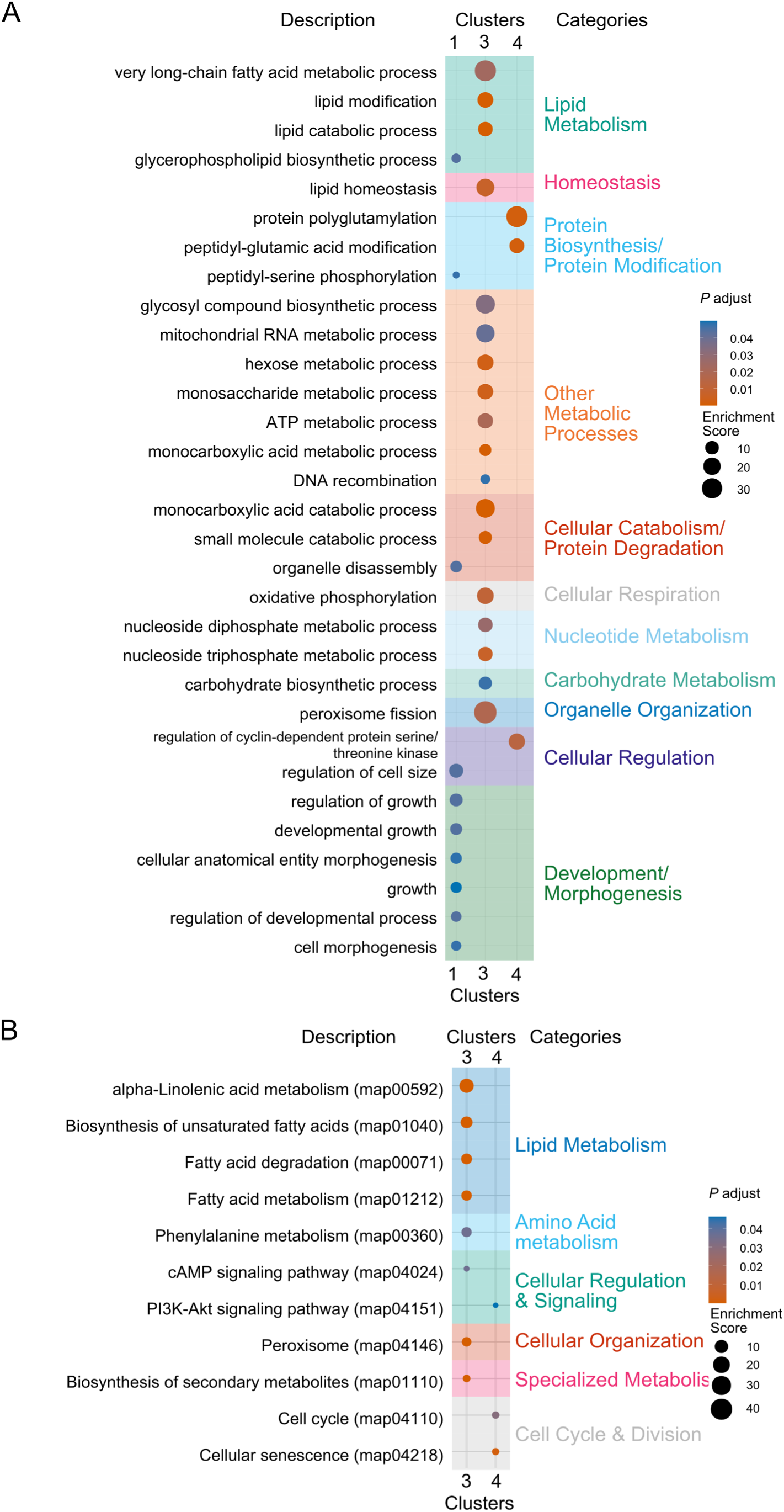
Functional enrichment of aposymbiotic-specific rhythmic clusters corresponding to fig. S6A-B. (**A**) Gene Ontology biological process enrichment (adjusted P < 0.05) for the four aposymbiotic-specific rhythmic clusters (C1-C4) derived from Gaussian mixture model clustering of 467 rhythmic genes identified by RAIN. Enriched terms are grouped into functional categories, including lipid metabolism, homeostasis, protein biosynthesis and modification, other metabolic processes, cellular catabolism and protein degradation, cellular respiration, nucleotide metabolism, carbohydrate metabolism, organelle organization, cellular regulation, and development and morphogenesis. Dot size indicates enrichment score, and color represents adjusted *P* value. (**B**) KEGG pathway enrichment for the same clusters, with significant pathways (adjusted *P* < 0.05) categorized into lipid metabolism, amino acid metabolism, cellular regulation and signaling, cellular organization, specialized metabolism, and cell cycle and division. These enrichments provide functional characterization of the temporal modules formed by aposymbiotic-specific rhythmic genes.

**Fig. S8.**
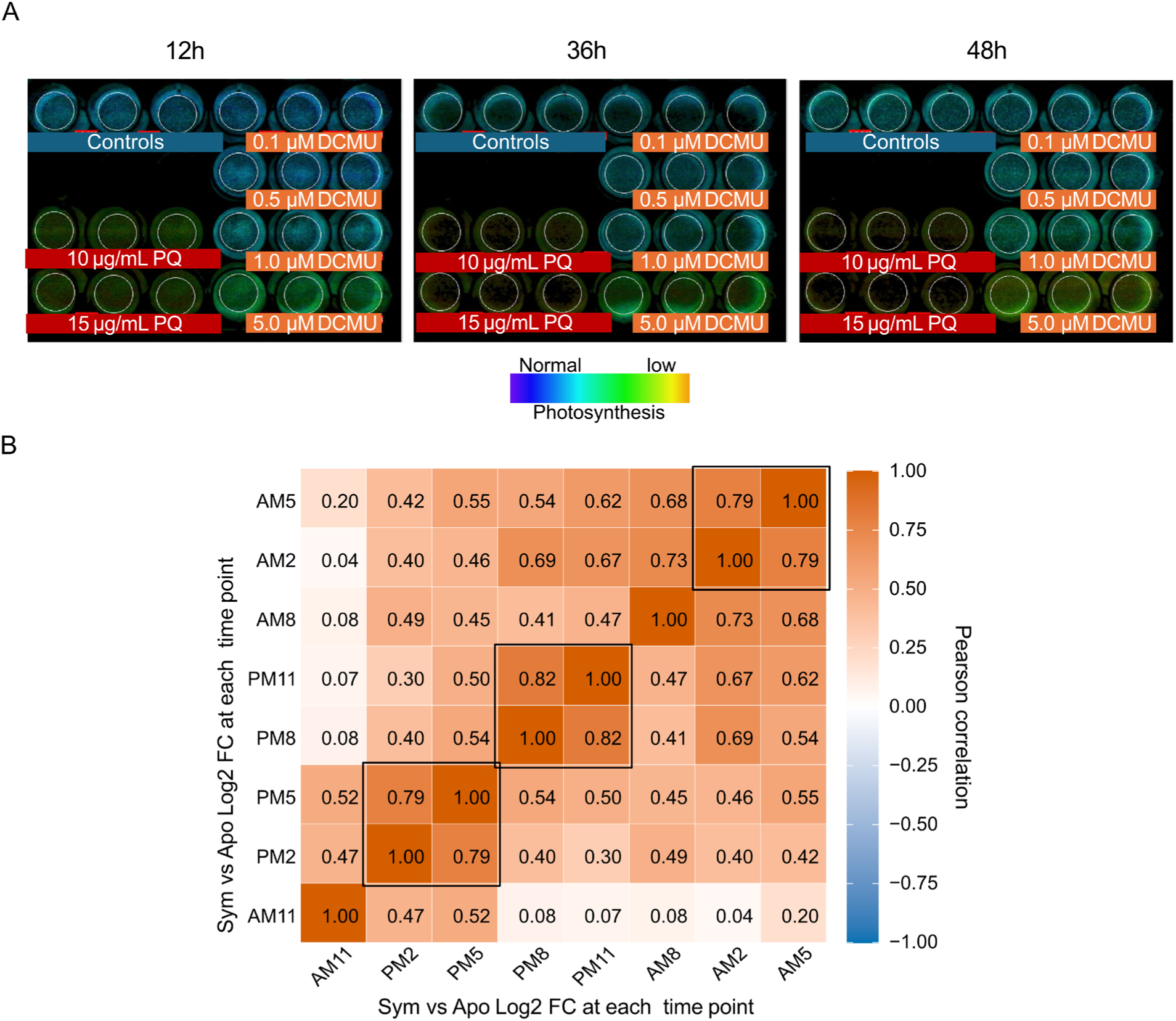
Photochemical response to paraquat and timepoint correlation analysis used for PQ experimental design. (**A**) Chlorophyll fluorescence images of symbiotic *P. bursaria* cultures exposed to control medium, DCMU (0.1-5.0 µM), or paraquat (10-15 µg/ml) at 12 h, 36 h, and 48 h. Fluorescence intensity is displayed on a pseudocolor scale from normal (blue) to low (yellow), illustrating the differential effects of PSII inhibition by DCMU and PSI inhibition by paraquat. (**B**) Pearson correlation matrix of Sym versus Apo Log₂ fold-change values across the eight diel timepoints (AM8, AM11, PM2, PM5, PM8, PM11, AM2, AM5). Blocks of high correlation (r ≥ 0.79-1.00) identify sets of redundant timepoints, supporting selection of AM8, AM11, PM5, and PM11 as representative sampling points for the paraquat RNA-seq experiment.

**Fig. S9.**
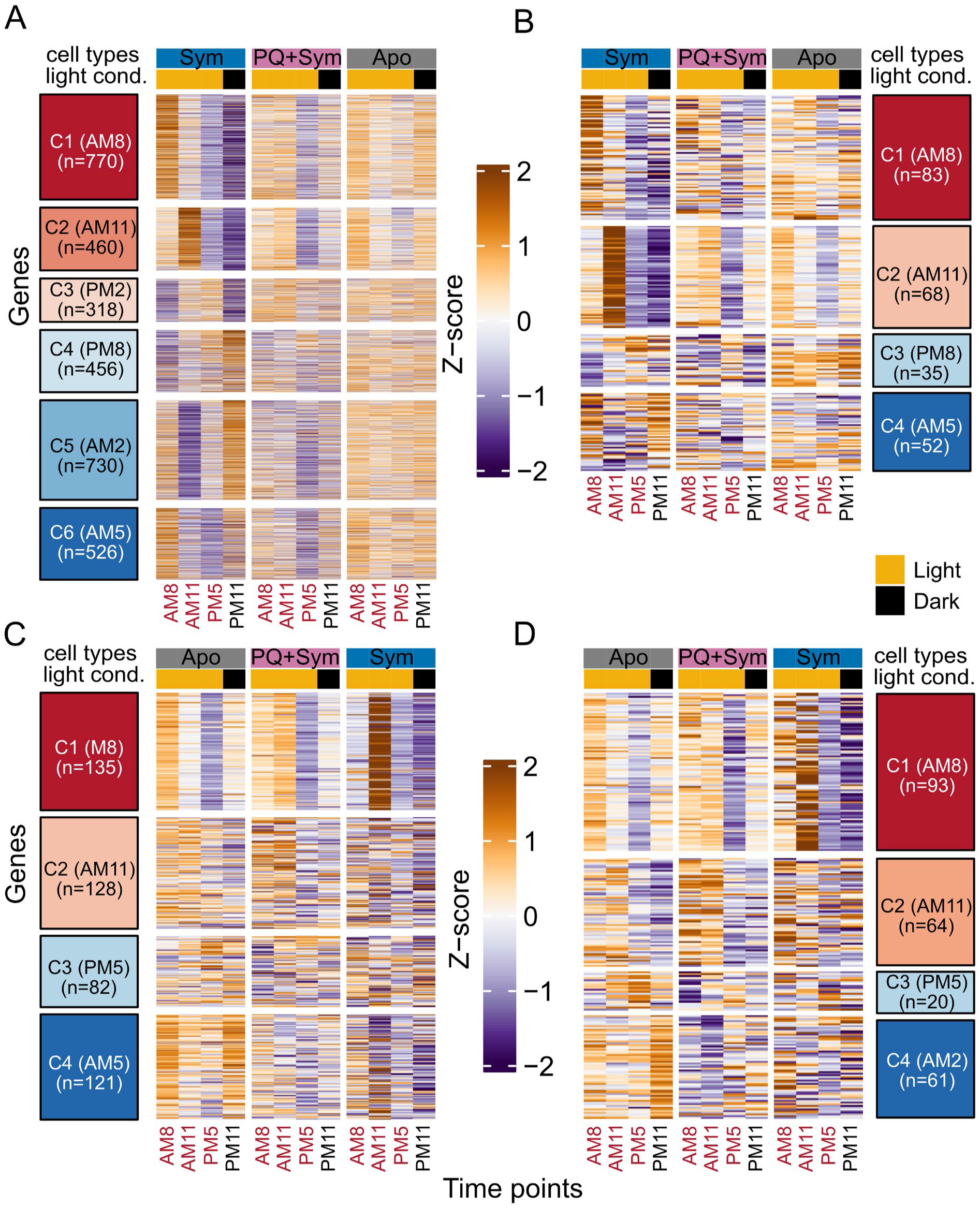
Gene-level expression heatmaps for temporal clusters shown as mean profiles in Fig. 3C-F. (**A**) Heatmaps of Z-scaled log_2_(TPM+1) expression for the six symbiotic-specific rhythmic clusters (C1-C6) across symbiotic (Sym), paraquat-treated symbiotic (PQ+Sym), and aposymbiotic (Apo) samples at the four matched zeitgeber timepoints (AM8, AM11, PM5, PM11). (**B**) Heatmaps of the four core rhythmic clusters (C1-C4) displayed across the same three conditions and four matched timepoints. (**C**) Heatmaps of the four aposymbiotic-specific rhythmic clusters (C1-C4) shown across Apo, PQ+Sym, and Sym samples at the same matched timepoints. (**D**) Heatmaps of the four paraquat-response clusters (C1-C4), summarizing distinct PQ-associated temporal response patterns across Apo, PQ+Sym, and Sym samples. For all panels, columns are ordered by time-of-day and annotated by cell type and light-dark condition as indicated; cluster labels denote the phase grouping used for the corresponding mean trajectory plot

**Fig. S10.**
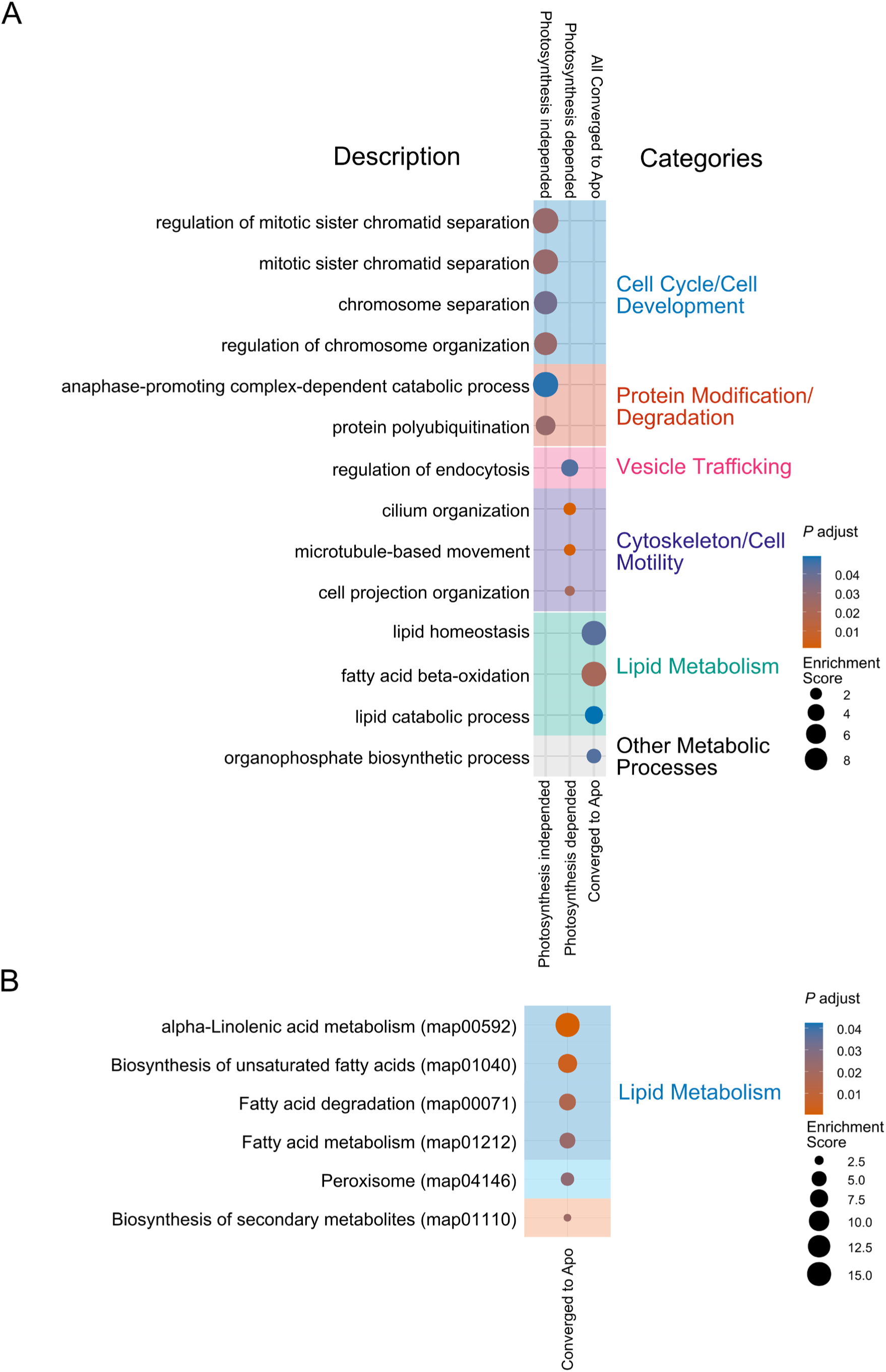
Functional enrichment of paraquat-response categories corresponding to Fig. 4. (**A**) Gene Ontology biological process enrichment (adjusted P < 0.05) for genes assigned to the PQ-insensitive, PQ-sensitive, and converged-to-aposymbiotic paraquat-response categories defined in Figure 4. Enriched terms are grouped into categories such as lipid metabolism, protein modification and degradation, vesicle trafficking, cytoskeleton and cell motility, other metabolic processes, and cell cycle and cell development. Dot size represents enrichment score, and color denotes adjusted P value. (**B**) KEGG pathway enrichment (adjusted *P* < 0.05) for the converged-to-aposymbiotic category, with significant pathways including fatty acid metabolism, fatty acid degradation, peroxisome-related processes, unsaturated fatty acid biosynthesis, α-linolenic acid metabolism, and secondary metabolite biosynthesis. Dot size indicates enrichment score, and color represents adjusted *P* value.

**Table S1.**
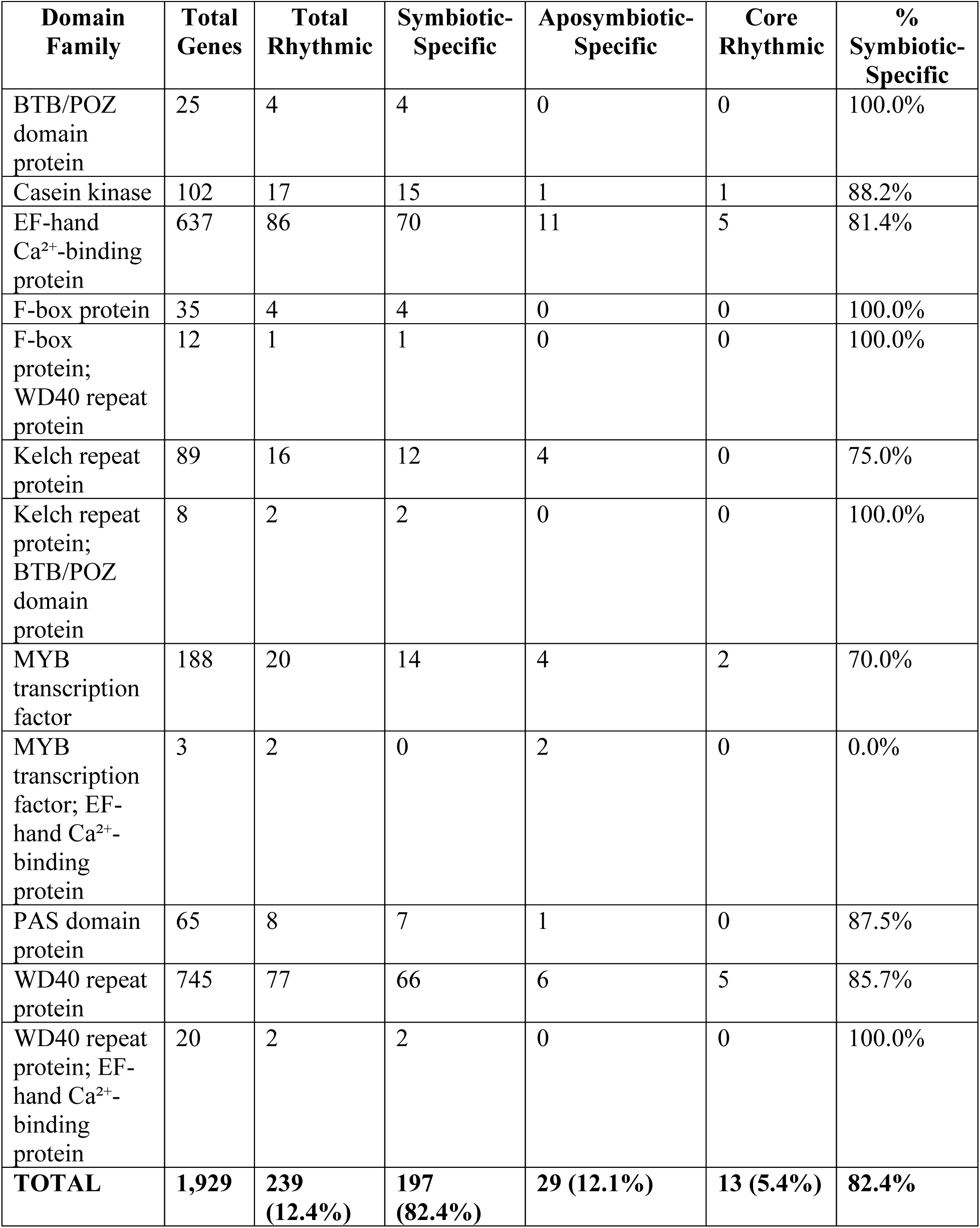
Rhythmicity of circadian-associated domain families across symbiotic-specific, aposymbiotic-specific, and core gene categories. Of 239 rhythmic domain genes identified (12.4% of 1,929 curated circadian-associated domains), 197 (82.4%) are symbiotic-specific, 29 (12.1%) are aposymbiotic-specific, and 13 (5.4%) are core rhythmic. The table includes single-domain families and multi-domain gene products. See Supporting Data 6 for complete gene-level annotations.

**Table S2.**
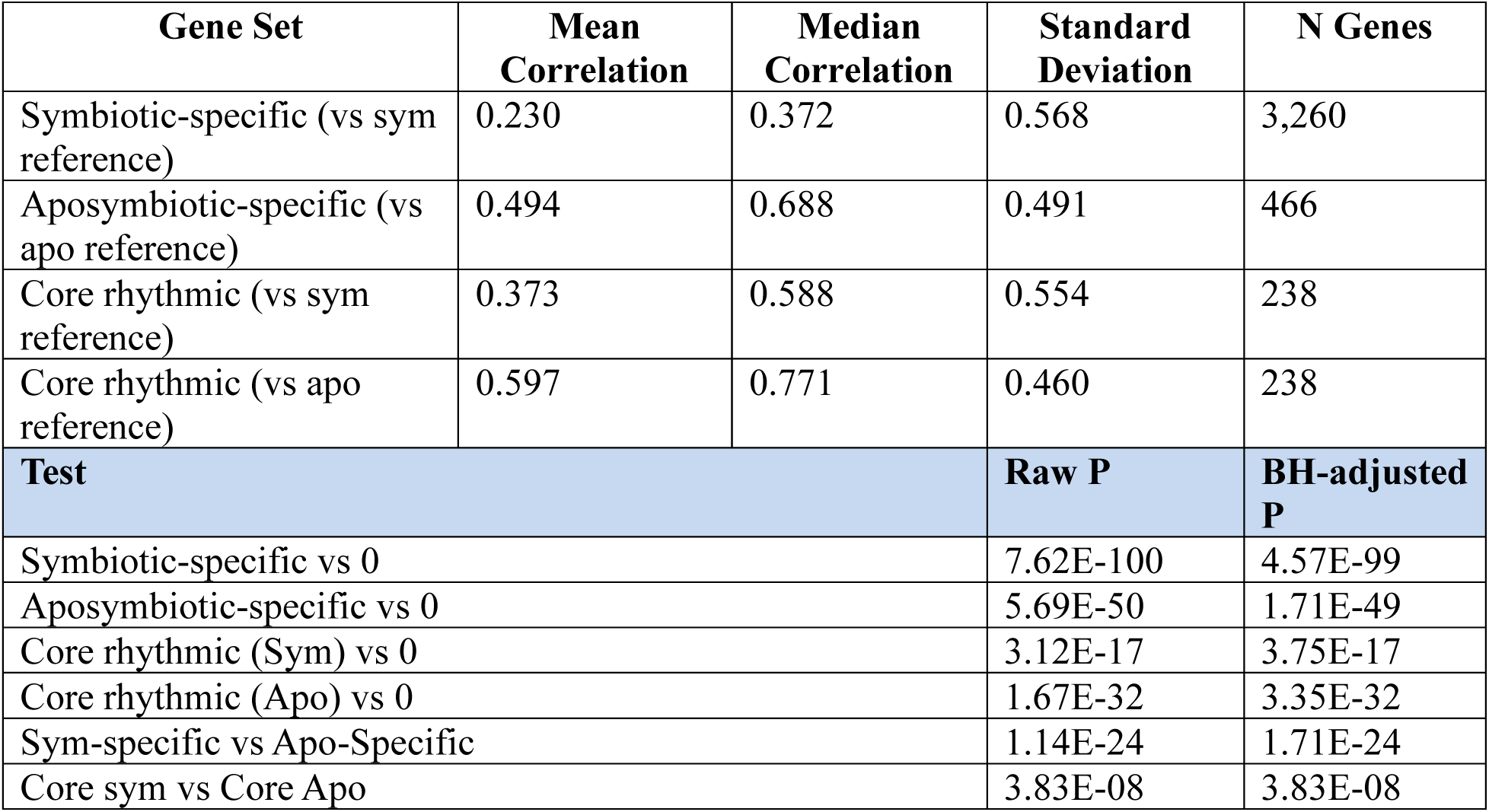
Pearson correlation of paraquat-treated four-timepoint expression profiles to symbiotic-specific, aposymbiotic-specific, and core rhythmic reference temporal profiles, with Wilcoxon tests. PQ-treated symbiotic cells (sampled at AM8, AM11, PM5, PM11) were evaluated across 3,260 symbiotic-specific, 466 aposymbiotic-specific, and 238 core rhythmic host genes. For each gene set, Pearson correlations (r) were computed between PQ-treated profiles and the corresponding 8-timepoint symbiotic or aposymbiotic reference temporal profiles and summarized by mean/median r and standard deviation. Statistical support is provided by distribution-level Wilcoxon tests (BH-corrected): one-sample signed-rank tests assess whether median r differs from 0 (tests vs 0), and rank-sum tests compare correlation distributions between symbiotic-specific vs aposymbiotic-specific sets and between core correlations computed against symbiotic vs aposymbiotic references. Because correlations are estimated from only four timepoints, inference is based on gene-set–level tests rather than per-gene correlation *P* values.

**Table S3.**
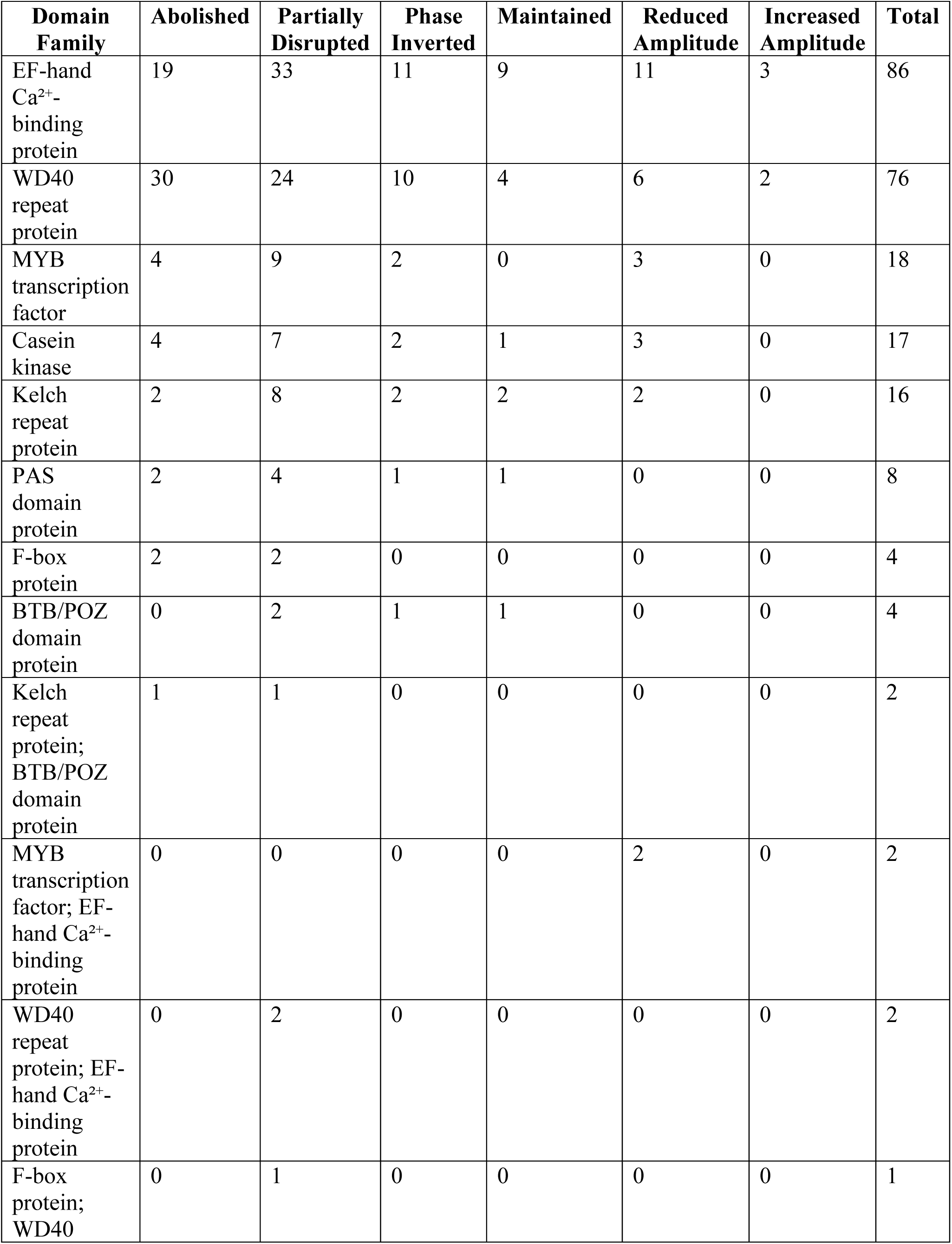
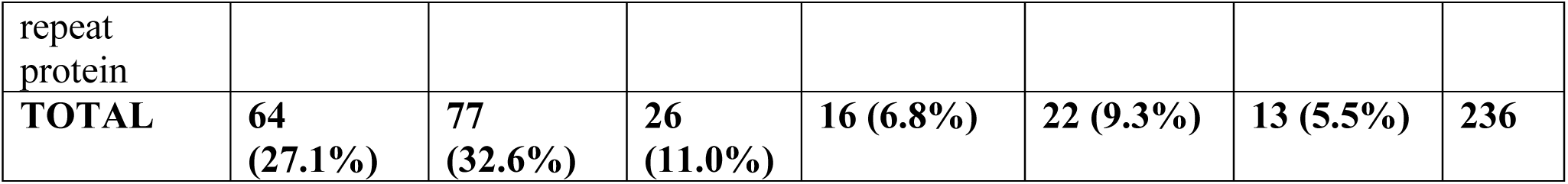
Distribution of circadian-associated domain genes across PQ disruption categories. Of 236 circadian-associated domain genes identified in symbiotic-specific, aposymbiotic-specific, and core rhythmic gene sets (derived from the 239 rhythmic domain genes in Supporting Table S1, excluding 3 bZIP genes not included in the original curation), 167 (70.8%) were preferentially disrupted under photosynthesis inhibition (Abolished + Partially Disrupted + Phase Inverted categories), while only 69 (29.2%) were maintained, dampened, or amplified. Domain families traditionally implicated in eukaryotic circadian control—EF-hand Ca²⁺-binding proteins (22.1% abolished, 38.4% partially disrupted), WD40 repeat proteins (39.5% abolished, 31.6% partially disrupted), and casein kinases (23.5% abolished, 41.2% partially disrupted)—showed the highest representation in disrupted categories, indicating that photosynthetic input is required to maintain the stability of core circadian-associated protein machinery.

**Table S4.**
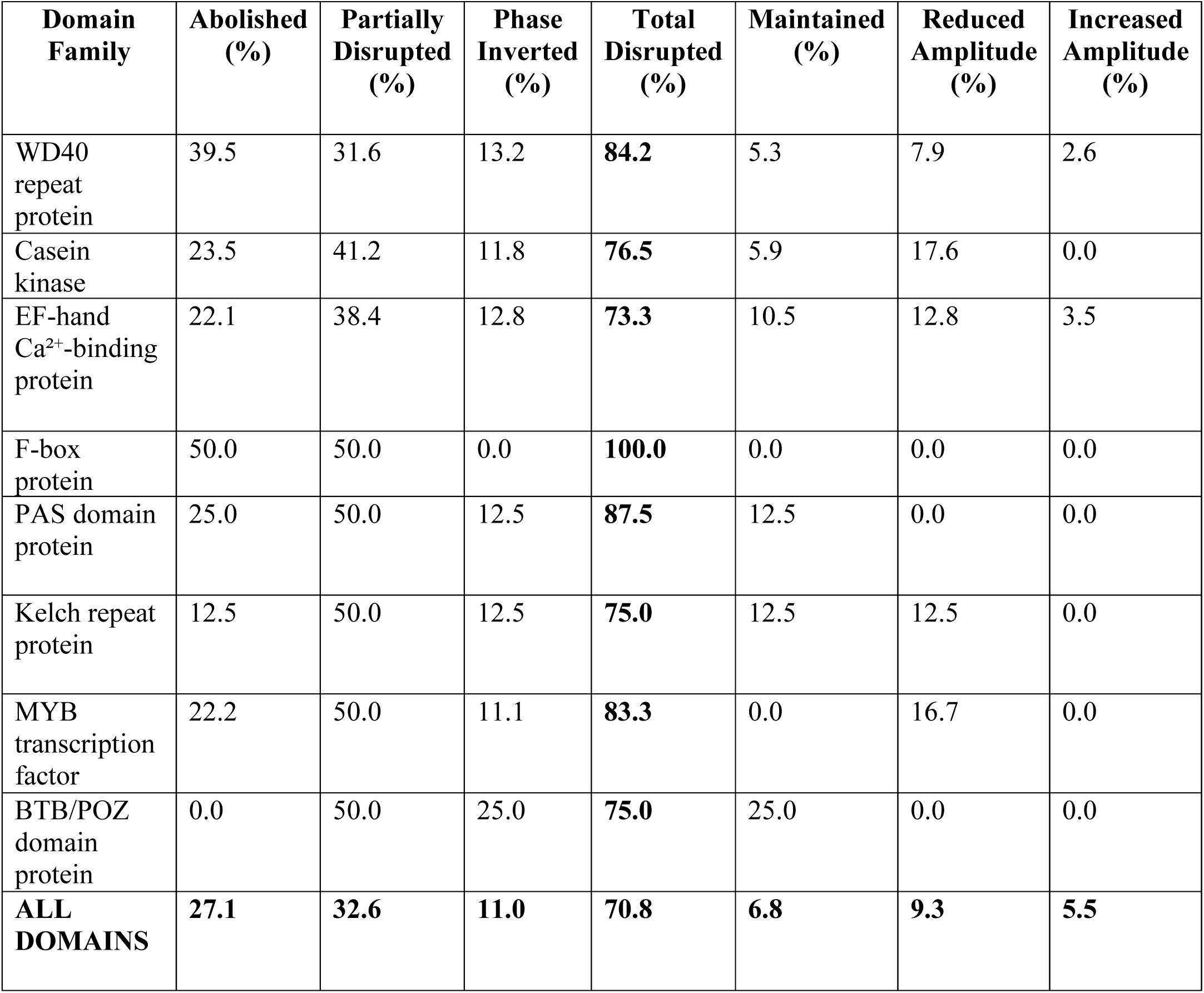
Percentage breakdown of circadian-associated domain gene responses to photosynthesis inhibition. For each domain family, the table shows the percentage of genes falling into each disruption category. Across all domain families, 70.8% of genes were disrupted (abolished, partially disrupted, or phase-inverted), with WD40 repeat proteins (84.2%), PAS domain proteins (87.5%), F-box proteins (100.0%), and MYB transcription factors (83.3%) showing particularly high disruption rates. Conversely, only 6.8% of domain genes were fully maintained, and 5.5% showed increased amplitude under PQ treatment. This pattern demonstrates that circadian-associated protein machinery is systematically destabilized when photosynthetic energy input is removed, indicating that the post-translational regulatory infrastructure underpinning temporal organization in *P. bursaria* depends fundamentally on endosymbiotic photosynthetic activity.

**Data file S1. Gene expression matrix and differential expression results for all conditions**

**(A)** TPM (transcripts per million) expression values for all 18,325 non-redundant *P. bursaria* DK2 pangenome genes across all three conditions: symbiotic (g_, 8 timepoints x 3 replicates), aposymbiotic (w_, 8 timepoints x 3 replicates), and paraquat-treated symbiotic (pq_, 4 timepoints x 3 replicates). Timepoints are AM8, AM11, PM2, PM5, PM8, PM11, AM2, and AM5 under 12:12 LD entrainment.

**(B)** TPM expression values for the 7,284 differentially expressed genes (DEGs; FDR <= 0.01, |log2FC| >= 1) identified in at least one pairwise symbiotic-versus-aposymbiotic timepoint comparison, across all timepoints and replicates for symbiotic and aposymbiotic conditions.

**(C)** Differential expression statistics from limma/eBayes analysis for all DEGs at each of the eight pairwise symbiotic-versus-aposymbiotic timepoint contrasts. Columns include gene ID, gene description, log2 fold change (logFC), average expression (AveExpr), t-statistic, raw *P* value, BH-adjusted *P* value (adj.P.Val), B-statistic, contrast label, and significance classification.

**(D)** Per-contrast summary of differential expression results across all eight timepoint comparisons, reporting total genes tested, number of significant DEGs, counts of up- and down-regulated genes, and their proportions of the total tested.

**Data file S2. Gaussian mixture model clustering and functional enrichment of differentially expressed genes**

**(A)** GMM cluster assignments for the 7,284 DEGs from joint symbiotic-aposymbiotic clustering into five temporal clusters (C1-C5), including maximum log2FC and minimum BH-adjusted *P* value per gene across all timepoint contrasts.

**(B)** GO biological process enrichment results for each of the five DEG temporal clusters. Columns include cluster ID, GO term ID and description, manually curated functional category, gene ratio, background ratio, rich factor, fold enrichment, z-score, raw *P* value, BH-adjusted *P* value, q value, contributing gene IDs, and gene count.

**(C)** KEGG pathway enrichment results for each of the five DEG temporal clusters. Columns include cluster ID, pathway ID and description, major and detailed functional groupings, gene ratio, background ratio, rich factor, fold enrichment, z-score, raw *P* value, BH-adjusted *P* value, q value, contributing gene IDs, and gene count.

**Data file S3. RAIN rhythmicity analysis and temporal cluster assignments for rhythmic gene sets**

**(A)** RAIN rhythmicity results for all 18,325 genes under symbiotic conditions, including raw *P* value, detected period (h), phase (h), BH-adjusted *P* value, rhythmicity classification (True/False at FDR < 0.05), and gene description.

**(B)** RAIN rhythmicity results for all 18,325 genes under aposymbiotic conditions, with the same fields as sheet A.

**(C)** GMM cluster assignments for the 3,300 symbiotic-specific rhythmic genes into six temporal clusters (C1-C6), with Z-score-normalized expression profiles across eight diel timepoints for both symbiotic and aposymbiotic conditions.

**(D)** GMM cluster assignments for the 467 aposymbiotic-specific rhythmic genes into four temporal clusters (C1-C4), with Z-score-normalized expression profiles across eight diel timepoints for both symbiotic and aposymbiotic conditions.

**(E)** GMM cluster assignments for the 238 core rhythmic genes (rhythmic in both conditions), including independent cluster assignments within symbiotic (Core_Sym_Cluster) and aposymbiotic (Core_Apo_Cluster) conditions, and Z-score-normalized expression profiles across eight diel timepoints.

**Data file S4. Functional enrichment of rhythmic gene clusters**

**(A)** GO biological process enrichment results (adjusted P < 0.05) for the six symbiotic-specific rhythmic gene clusters (C1-C6) identified by GMM clustering of 3,300 genes. Columns include cluster ID, GO term ID and description, major functional category, gene ratio, background ratio, rich factor, fold enrichment, z-score, raw *P* value, BH-adjusted *P* value, q value, contributing gene IDs, and gene count.

**(B)** KEGG pathway enrichment results (adjusted P < 0.05) for the six symbiotic-specific rhythmic gene clusters (C1-C6). Columns follow the same structure as sheet A with KEGG pathway IDs and functional category groupings.

**(C)** GO biological process enrichment results (adjusted P < 0.05) for the four aposymbiotic-specific rhythmic gene clusters (C1-C4) identified by GMM clustering of 467 genes.

**(D)** KEGG pathway enrichment results (adjusted P < 0.05) for the four aposymbiotic-specific rhythmic gene clusters (C1-C4).

**Data file S5. TTFL clock gene orthology screen and timing-associated domain gene annotations**

**(A)** Results of the dual-method (BLASTp + DIAMOND) orthology screen for 42 TTFL reference clock proteins from animals (CLOCK, BMAL1, PER, CRY, NR1D, RORA, RORB), plants (CCA1, LHY, RVE, APRR), fungi (FRQ, WC-1, WC-2), and cyanobacteria (KaiA/B/C) against the *P. bursaria* DK2 proteome. Columns include reference gene name, source organism, best *P. bursaria* hit, BLASTp e-value, sequence identity (%), alignment length (aa), query and subject protein lengths (aa), query and subject coverage (%), DIAMOND hit consistency, domain-level classification from InterProScan, and rhythmicity status under LD entrainment.

**(B)** Annotations for 1,929 genes bearing timing-associated protein domains curated from the *P. bursaria* DK2 proteome, including casein kinases, EF-hand Ca2+-binding proteins, F-box proteins, Kelch-repeat proteins, WD40 proteins, PAS domain proteins, MYB transcription factors, and BTB/POZ domain proteins. Columns include gene ID, symbiotic and aposymbiotic GMM cluster assignments, domain category, rhythmic status in symbiotic and aposymbiotic conditions, phase assignments (h), and RAIN *P* values for both conditions.

**Data file S6. Paraquat photochemical measurement and gene-level temporal response data**

**(A)** Photosystem II maximum quantum yield (Fv/Fm) measurements over 48 h for symbiotic *P. bursaria* cultures exposed to control medium, DCMU (0.01-5.0 uM), or paraquat (10-15 ug/mL). Columns include treatment, time point (h), inhibitor type, mean Fv/Fm, and standard error.

**(B)** Gene-level paraquat response results for all rhythmic gene sets (symbiotic-specific, n = 3,260; aposymbiotic-specific, n = 466; core rhythmic, n = 238). Columns include interpretation category, gene ID, gene set, reference condition, Pearson correlation to reference, reference and PQ-treated amplitudes, amplitude ratio, mean expression values, mean fold change, and disruption category classification.

**(C)** Summary of PQ response category distributions by gene set, reporting counts and percentages for each classification (maintained, maintained with increased/reduced amplitude, partially disrupted, rhythm abolished, phase inverted, converged to aposymbiotic pattern, converged with increased amplitude, partial convergence with dampening, weak convergence, no convergence arrhythmic, anti-phase to aposymbiotic).

**(D)** PQ disruption results for the 236 timing-associated circadian domain genes identified within rhythmic gene sets, with gene-level Pearson correlation, amplitude, amplitude ratio, interpretation category, gene set, and domain family annotation.

**Data file S7. Functional enrichment of paraquat response categories**

**(A)** GO biological process enrichment results (adjusted P < 0.05) for three PQ response categories: photosynthesis-independent (PQ-insensitive, maintained symbiotic pattern), photosynthesis-dependent (PQ-sensitive, disrupted), and converged-to-aposymbiotic. Columns include response category, GO term ID and description, functional category grouping, gene ratio, background ratio, rich factor, fold enrichment, z-score, raw *P* value, BH-adjusted *P* value, q value, contributing gene IDs, and gene count.

**(B)** KEGG pathway enrichment results (adjusted P < 0.05) for PQ response categories. Columns include response category, pathway ID and description, KEGG functional category, gene ratio, background ratio, rich factor, fold enrichment, z-score, raw *P* value, BH-adjusted *P* value, q value, contributing gene IDs, and gene count.

**Data file S8. Cross-species conservation of diel expression programs between *P. bursaria* and T. utriculariae.**

**(A)** One-to-one ortholog pairs between *P. bursaria* and T. utriculariae derived from the union of three complementary orthology inference methods: Reciprocal Best BLAST Hits (RBH), OrthoFinder v2.5.4, and OrthoMCL. Columns include *P. bursaria* gene ID, T. utriculariae gene ID, source methods supporting the pair, and number of supporting methods.

**(B)** Ortholog coverage statistics by rhythmic gene set after 1:1 filtering, reporting total rhythmic *P. bursaria* genes per category (symbiotic-specific, aposymbiotic-specific, Core Rhythmic Sym, Core Rhythmic Apo), number with a T. utriculariae ortholog, and number of retained 1:1 pairs.

**(C)** Gene-level Pearson correlation between *P. bursaria* and T. utriculariae Z-score-normalized temporal expression profiles across six matched zeitgeber timepoints (AM8, AM11, PM2, PM5, PM8, PM11). Columns include *P. bursaria* gene ID, gene set, number of T. utriculariae orthologs, T. utriculariae gene ID, Pearson correlation, raw *P* value, BH-adjusted *P* value, 95% confidence interval bounds, and maximum circular correlation.

**(D)** Statistical test results for cross-species and within-species correlation distributions across all gene sets, including per-gene-set summary statistics (n, mean and median cross-species correlation, mean and median within-*P. bursaria* and within-*T. utriculariae* correlations) and Wilcoxon signed-rank and rank-sum test results with BH-adjusted *P* values.

## Notes

### Competing Interest Statement

The authors have declared no competing interest.

