## Supporting figures S1 to S10, and Table S1 to S4 for "Endosymbiotic algal photosynthesis shapes diel transcriptome architecture in its ciliate host *Paramecium bursaria*"

Supplementary Materials for

### **Photosynthetic endosymbionts influence the temporal transcriptome in the *Paramecium bursaria* host**

Md Mostafa Kamal *et al.*

**This PDF file includes:**

Figs. S1 to S10
Tables S1 to S4
Data file S1 to S8 captions

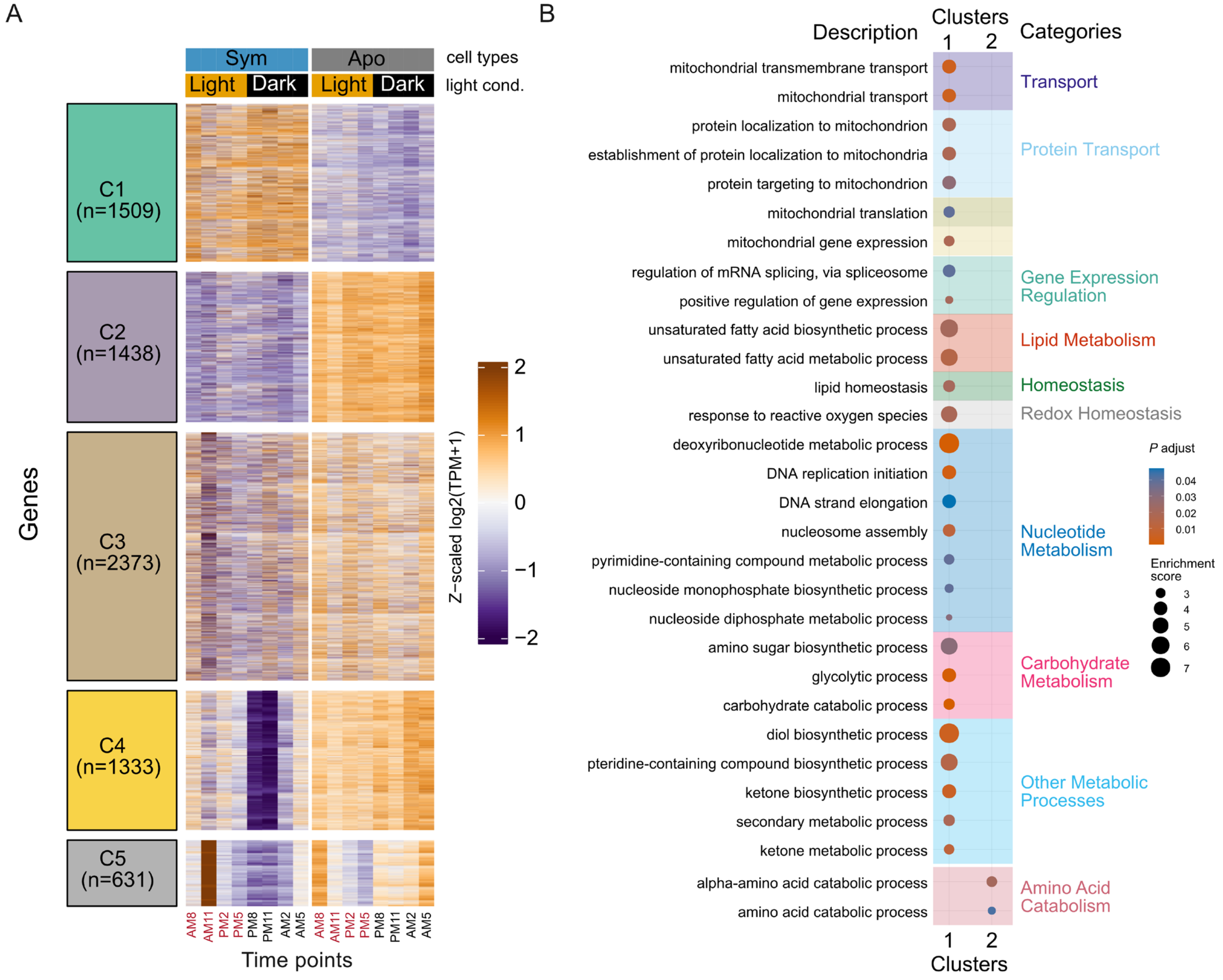


**Fig. S1. Temporal GMM clustering of differentially expressed genes and GO biological process enrichment of Clusters 1 and 2 in *P. bursaria*.**

**(A)** Gaussian mixture model (GMM) clustering of 7,284 differentially expressed genes (FDR < 0.01; $|\log_2FC| > 1$) across the combined dataset. Five clusters (C1-C5) are shown with Z-scaled $\log_2(TPM+1)$ expression profiles across all timepoints for both conditions, annotated by condition and light-dark status; corresponding mean temporal trajectories for each cluster are provided in fig. S2. (**B**) Gene Ontology (GO) biological process enrichment for C1 and C2, with enriched terms grouped into functional categories. Dot size indicates enrichment score, and color represents adjusted *P* value; GO enrichment for Clusters 4 and 5 is in fig. S3; no significant enrichment was detected for Cluster 3.

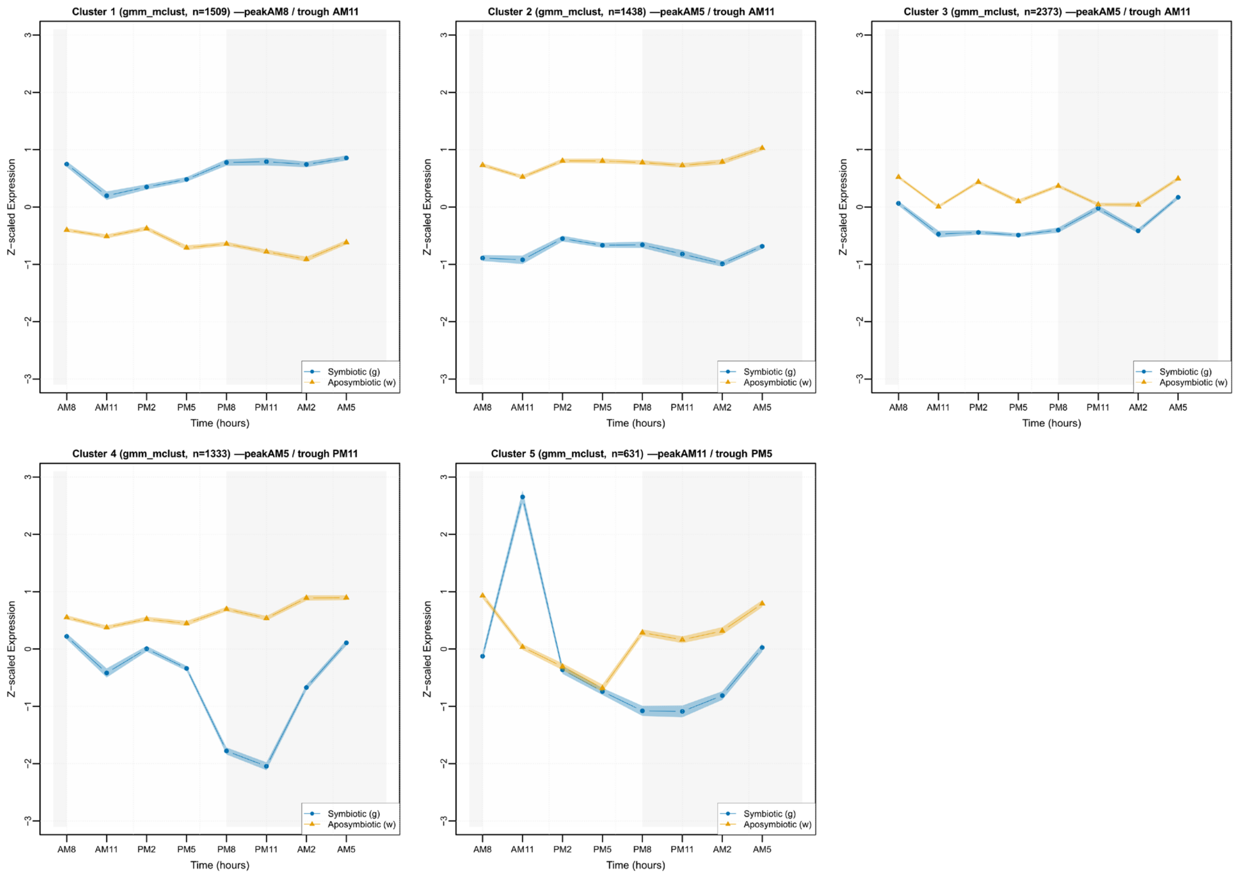


**Fig. S2. Mean temporal expression profiles for the five joint GMM clusters corresponding to heatmaps in Fig. S1A.**

Mean Z-scaled $\log_2$(TPM+1) expression trajectories for genes assigned to Cluster 1 (n = 1,509), Cluster 2 (n = 1,438), Cluster 3 (n = 2,373), Cluster 4 (n = 1,333), and Cluster 5 (n = 631) derived from Gaussian mixture model clustering of the joint symbiotic and aposymbiotic dataset. These trajectories correspond to the cluster heatmaps shown in **Fig. S1A**. Each panel shows the average expression pattern across eight diel time points (AM8, AM11, PM2, PM5, PM8, PM11, AM2, AM5) for symbiotic (g) and aposymbiotic (w) cells, plotted in chronological order.

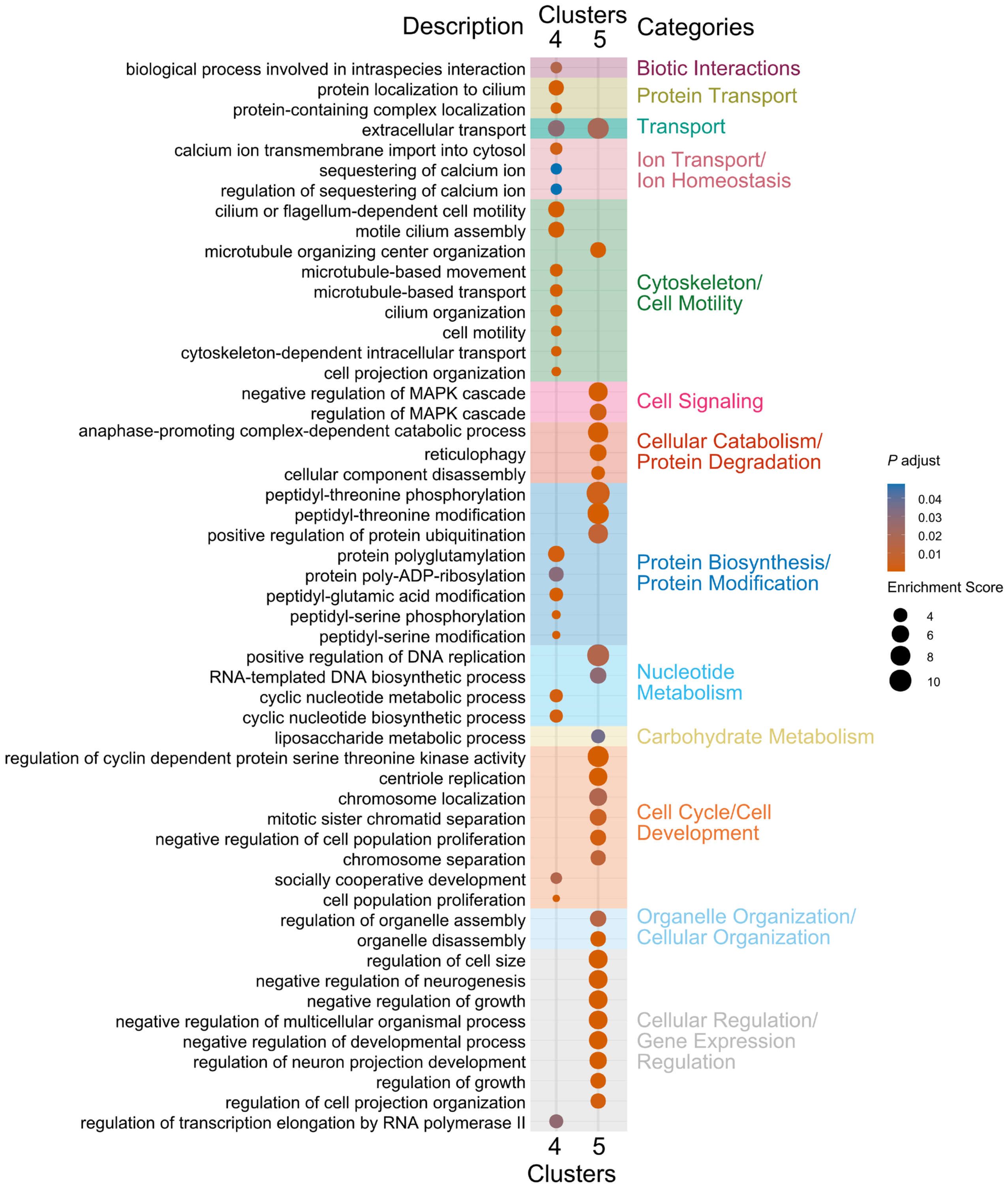
Description
Clusters
4
5
Categories
biological process involved in intraspecies interaction
protein localization to cilium
protein-containing complex localization
extracellular transport
calcium ion transmembrane import into cytosol
sequestering of calcium ion
regulation of sequestering of calcium ion
cilium or flagellum-dependent cell motility
motile cilium assembly
microtubule organizing center organization
microtubule-based movement
microtubule-based transport
cilium organization
cell motility
cytoskeleton-dependent intracellular transport
cell projection organization
negative regulation of MAPK cascade
regulation of MAPK cascade
anaphase-promoting complex-dependent catabolic process
reticulophagy
cellular component disassembly
peptidyl-threonine phosphorylation
peptidyl-threonine modification
positive regulation of protein ubiquitination
protein polyglutamylation
protein poly-ADP-ribosylation
peptidyl-glutamic acid modification
peptidyl-serine phosphorylation
peptidyl-serine modification
positive regulation of DNA replication
RNA-templated DNA biosynthetic process
cyclic nucleotide metabolic process
cyclic nucleotide biosynthetic process
liposaccharide metabolic process
regulation of cyclin dependent protein serine threonine kinase activity
centriole replication
chromosome localization
mitotic sister chromatid separation
negative regulation of cell population proliferation
chromosome separation
socially cooperative development
cell population proliferation
regulation of organelle assembly
organelle disassembly
regulation of cell size
negative regulation of neurogenesis
negative regulation of growth
negative regulation of multicellular organismal process
negative regulation of developmental process
regulation of neuron projection development
regulation of growth
regulation of cell projection organization
regulation of transcription elongation by RNA polymerase II
Biotic Interactions
Protein Transport
Transport
Ion Transport/
Ion Homeostasis
Cytoskeleton/
Cell Motility
Cell Signaling
Cellular Catabolism/
Protein Degradation
Protein Biosynthesis/
Protein Modification
Nucleotide
Metabolism
Carbohydrate Metabolism
Cell Cycle/Cell
Development
Organelle Organization/
Cellular Organization
Cellular Regulation/
Gene Expression
Regulation
P adjust
0.04
0.03
0.02
0.01
Enrichment Score
4
6
8
10
4
5
Clusters

**Fig. S3. GO biological process enrichment for Clusters 4 and 5 corresponding to Fig. S1B.**

Gene Ontology biological process enrichment for genes assigned to Cluster 4 and Cluster 5 from the joint GMM clustering shown in fig. S1A, complementing the enrichment analysis for Clusters 1 and 2 in fig. S1B. Enriched terms (adjusted $P < 0.05$) are displayed and grouped into manually curated functional categories, including biotic interactions, ion transport and homeostasis, protein transport, cytoskeleton and cell motility, cellular organization, cell signaling, cellular catabolism and protein degradation, nucleotide metabolism, carbohydrate metabolism, cell cycle and development, and protein biosynthesis and modification. Dot size indicates enrichment score, and color denotes adjusted $P$ value.

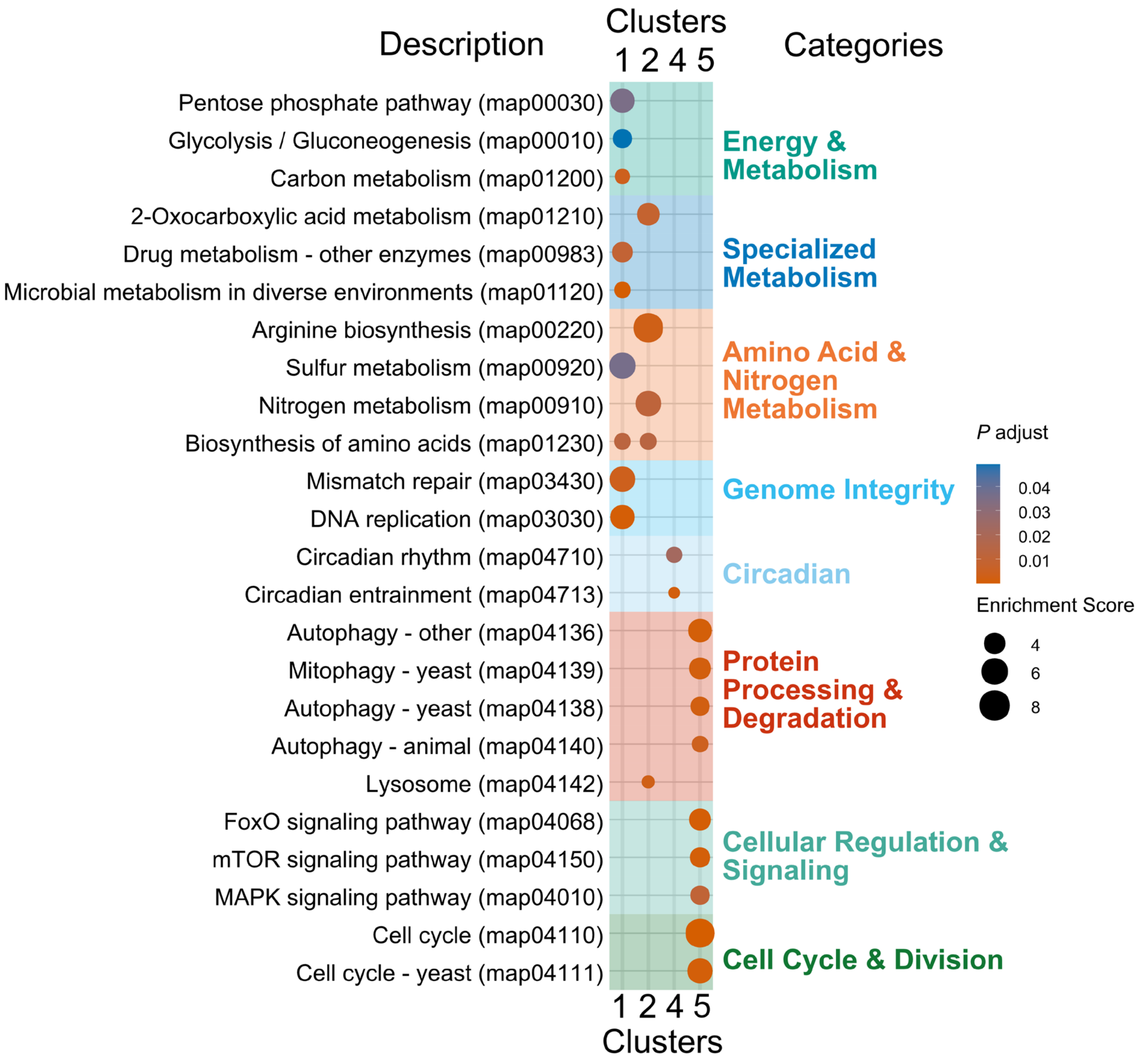


**Fig. S4. KEGG pathway enrichment for GMM clusters corresponding to fig. S1A.** KEGG pathway enrichment for genes assigned to Clusters 1, 2, 4, and 5 from the joint GMM clustering shown in fig. S1A, complementing the GO biological process enrichments presented in fig. S1B and figs. S3. Enriched KEGG pathways (adjusted $P < 0.05$) are displayed with functional groupings, including energy and carbon metabolism, specialized metabolism, amino acid and nitrogen metabolism, genome integrity, circadian pathways, protein processing and degradation, cofactor metabolism, cellular organization, cellular regulation and signaling, cell cycle and division, and immune and defense processes. Dot size indicates enrichment score, and color represents adjusted *P* value.

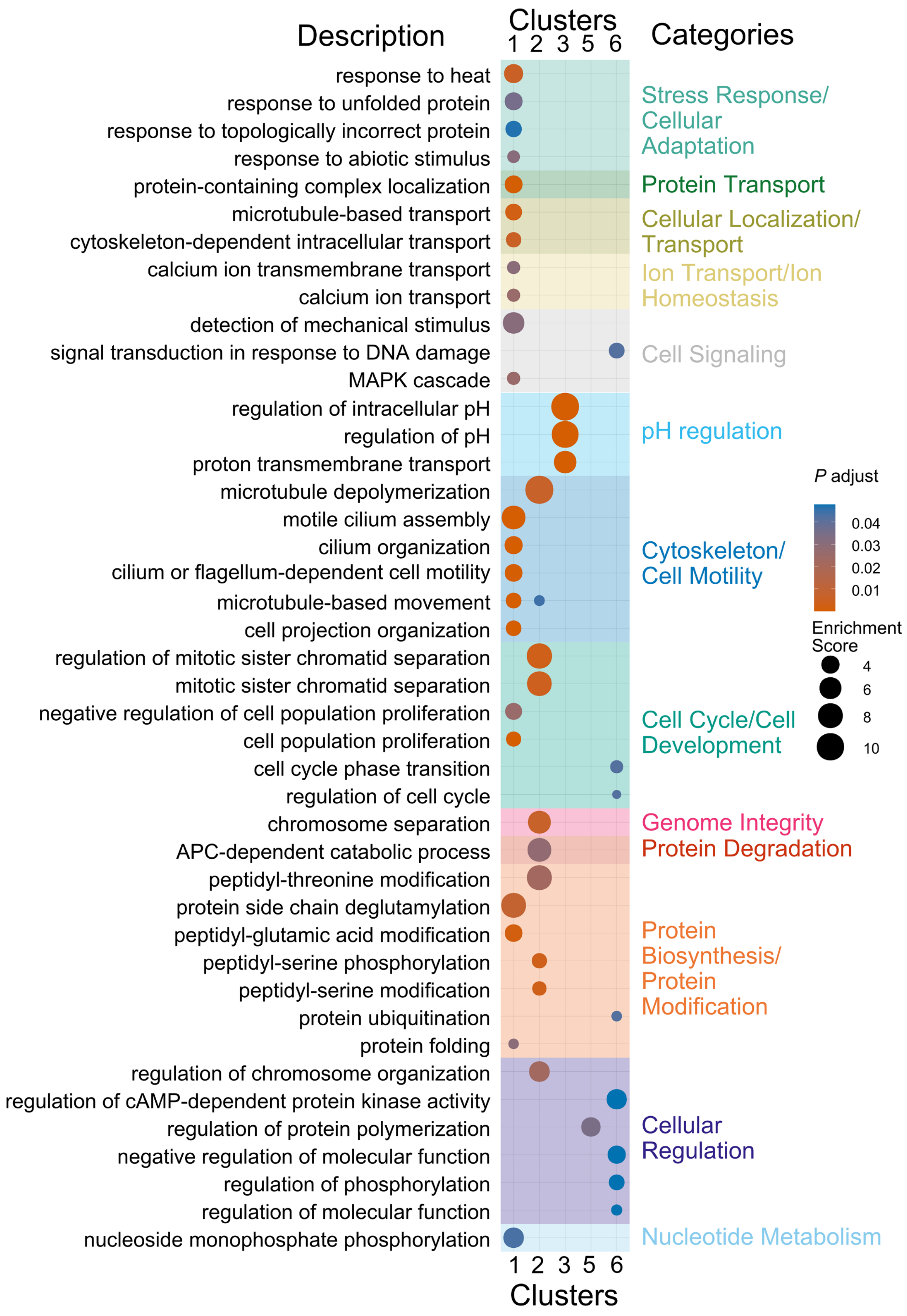
Description
Clusters
1 2 3 5 6
Categories
response to heat
response to unfolded protein
response to topologically incorrect protein
response to abiotic stimulus
protein-containing complex localization
microtubule-based transport
cytoskeleton-dependent intracellular transport
calcium ion transmembrane transport
calcium ion transport
detection of mechanical stimulus
signal transduction in response to DNA damage
MAPK cascade
regulation of intracellular pH
regulation of pH
proton transmembrane transport
microtubule depolymerization
motile cilium assembly
cilium organization
cilium or flagellum-dependent cell motility
microtubule-based movement
cell projection organization
regulation of mitotic sister chromatid separation
mitotic sister chromatid separation
negative regulation of cell population proliferation
cell population proliferation
cell cycle phase transition
regulation of cell cycle
chromosome separation
APC-dependent catabolic process
peptidyl-threonine modification
protein side chain deglutamylation
peptidyl-glutamic acid modification
peptidyl-serine phosphorylation
peptidyl-serine modification
protein ubiquitination
protein folding
regulation of chromosome organization
regulation of cAMP-dependent protein kinase activity
regulation of protein polymerization
negative regulation of molecular function
regulation of phosphorylation
regulation of molecular function
nucleoside monophosphate phosphorylation
Stress Response/ Cellular Adaptation
Protein Transport
Cellular Localization/ Transport
Ion Transport/Ion Homeostasis
Cell Signaling
pH regulation
Cytoskeleton/ Cell Motility
Cell Cycle/Cell Development
Genome Integrity
Protein Degradation
Protein Biosynthesis/ Protein Modification
Cellular Regulation
Nucleotide Metabolism
P adjust
0.04
0.03
0.02
0.01
Enrichment Score
4
6
8
10
1 2 3 5 6
Clusters

**Fig. S5. GO biological process enrichment for the symbiotic-specific rhythmic clusters shown in Fig. 2E-F.**

Gene Ontology biological process enrichment (adjusted $P < 0.05$) for the six symbiotic-specific rhythmic clusters (C1-C6) derived from Gaussian mixture model clustering of 3,300 rhythmic genes. Enriched terms are grouped into functional categories, including stress response and cellular adaptation, protein transport, cellular localization and intracellular transport, ion transport and homeostasis, cell signaling, pH regulation, cytoskeleton and cell motility, cell cycle and cell development, genome integrity and protein degradation, protein biosynthesis and post-translational modification, and nucleotide metabolism. Dot size represents enrichment score, and color denotes adjusted *P* value. These enrichments provide functional characterization of the temporal gene modules shown in Fig. 2E-F.

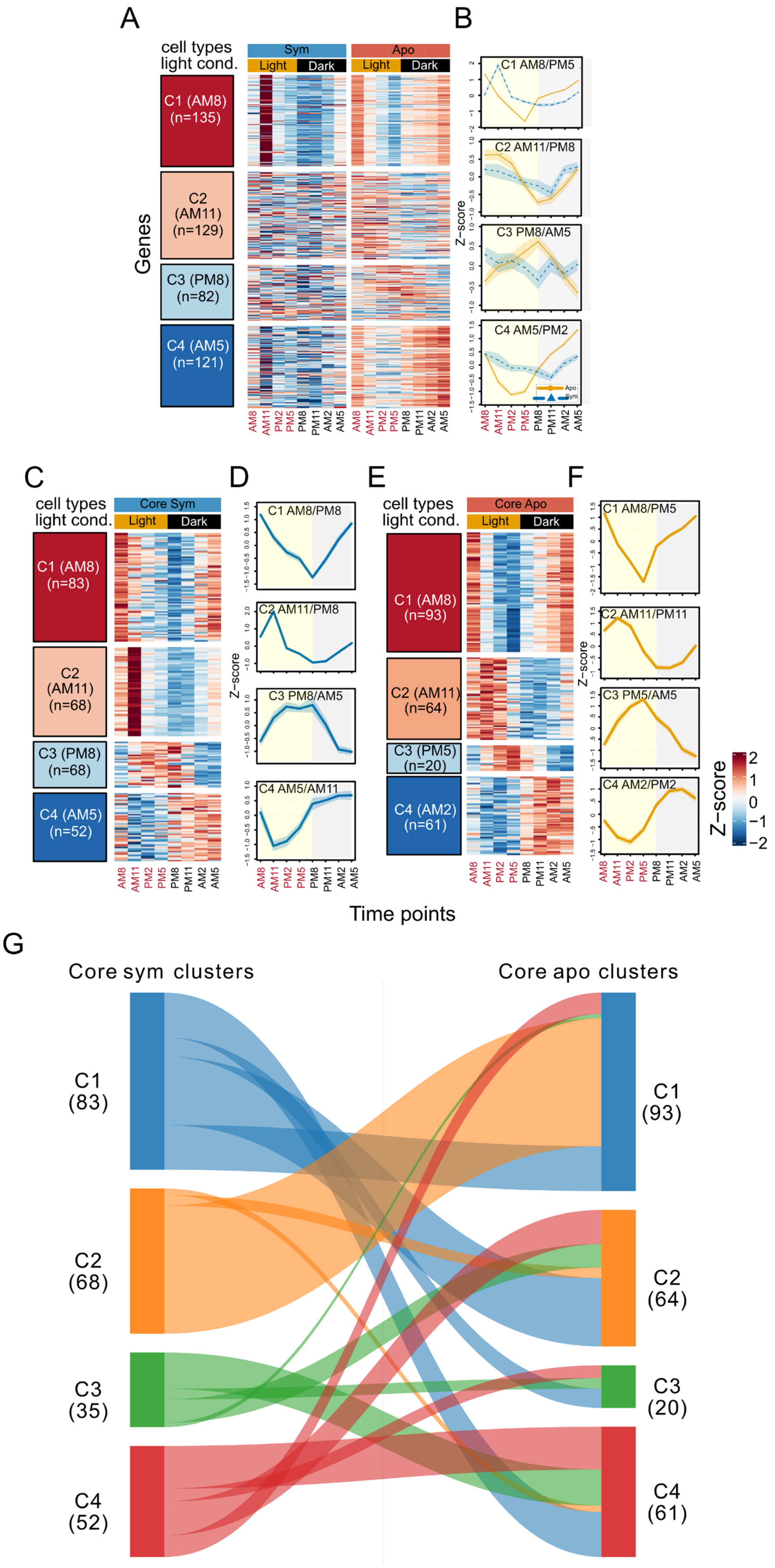
A
B
cell types
light cond.
Sym
Apo
Light
Dark
C1 (AM8) (n=135)
C2 (AM11) (n=129)
C3 (PM8) (n=82)
C4 (AM5) (n=121)
Genes
C1 AM8/PM5
C2 AM11/PM8
C3 PM8/AM5
C4 AM5/PM2
Z−score
Apo
Sym
AM8
AM11
PM2
PM5
PM8
PM11
AM2
AM5
C
D
E
F
Core Sym
C1 (AM8) (n=83)
C2 (AM11) (n=68)
C3 (PM8) (n=68)
C4 (AM5) (n=52)
C1 AM8/PM8
C2 AM11/PM8
C3 PM8/AM5
C4 AM5/AM11
Core Apo
C1 (AM8) (n=93)
C2 (AM11) (n=64)
C3 (PM5) (n=20)
C4 (AM2) (n=61)
C1 AM8/PM5
C2 AM11/PM11
C3 PM5/AM5
C4 AM2/PM2
Time points
G
Core sym clusters
Core apo clusters
C1 (83)
C2 (68)
C3 (35)
C4 (52)
C1 (93)
C2 (64)
C3 (20)
C4 (61)

**Fig. S6. Clustering of aposymbiotic-specific rhythmic genes and condition-specific clustering of the core rhythmic scaffold supporting Fig. 2A, C, and D.**
(**A**) Heatmap of Z-scored expression for the 467 aposymbiotic-specific rhythmic genes clustered into four temporal groups (C1-C4; gene counts shown) across eight diel timepoints (AM8, AM11, PM2, PM5, PM8, PM11, AM2, AM5). Expression is shown for both symbiotic and aposymbiotic samples, with column annotations indicating cell type and light-dark condition. (**B**) Mean Z-scored temporal trajectories for the four aposymbiotic-specific clusters in panel (A), with cluster labels indicating the approximate peak/trough timing (C1 AM8/PM5, C2 AM11/PM8, C3 PM8/AM5, C4 AM5/PM2). (**C-D**) Gaussian mixture model clustering of the 238 core rhythmic genes performed within symbiotic cells ("Core Sym"), shown as a cluster heatmap (**C**) and corresponding mean trajectories (**D**) for four clusters (C1-C4; gene counts shown; labels indicate approximate peak/trough timing: C1 AM8/PM8, C2 AM11/PM8, C3 PM8/AM5, C4 AM5/AM11). (**E-F**) Gaussian mixture model clustering of the same 238 core rhythmic genes performed within aposymbiotic cells ("Core Apo"), shown as a cluster heatmap (**E**) and mean trajectories (**F**) for four clusters (C1-C4; gene counts shown; labels indicate approximate peak/trough timing: C1 AM8/PM5, C2 AM11/PM11, C3 PM5/AM5, C4 AM2/PM2). (**G**) Sankey/alluvial mapping linking core-cluster assignments between the symbiotic and aposymbiotic clustering results, illustrating redistribution of the same core genes among condition-specific temporal trajectory classes.

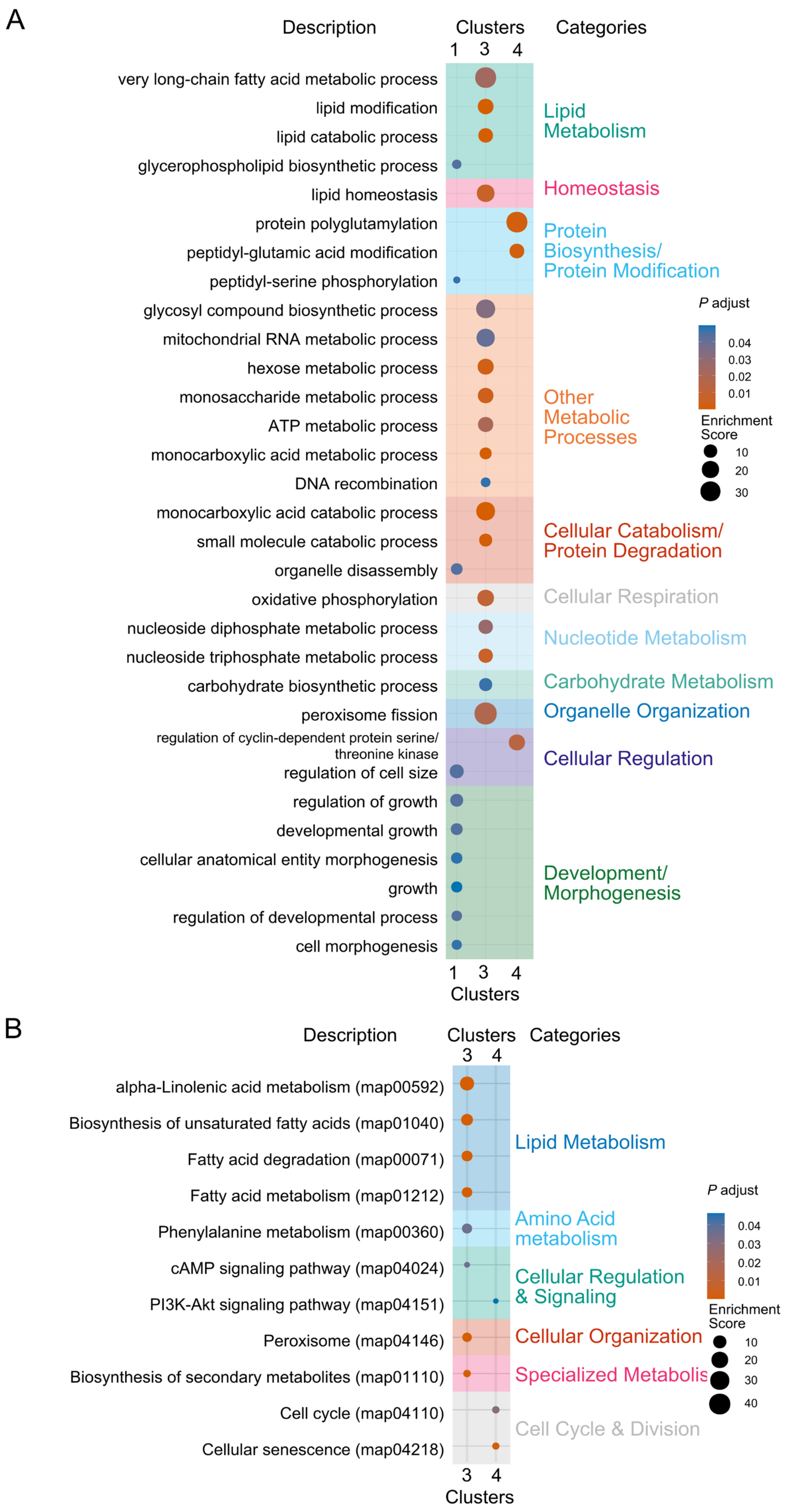
A
Description
Clusters
Categories
1 3 4
very long-chain fatty acid metabolic process
lipid modification
lipid catabolic process
glycerophospholipid biosynthetic process
lipid homeostasis
protein polyglutamylation
peptidyl-glutamic acid modification
peptidyl-serine phosphorylation
glycosyl compound biosynthetic process
mitochondrial RNA metabolic process
hexose metabolic process
monosaccharide metabolic process
ATP metabolic process
monocarboxylic acid metabolic process
DNA recombination
monocarboxylic acid catabolic process
small molecule catabolic process
organelle disassembly
oxidative phosphorylation
nucleoside diphosphate metabolic process
nucleoside triphosphate metabolic process
carbohydrate biosynthetic process
peroxisome fission
regulation of cyclin-dependent protein serine/threonine kinase
regulation of cell size
regulation of growth
developmental growth
cellular anatomical entity morphogenesis
growth
regulation of developmental process
cell morphogenesis
Lipid Metabolism
Homeostasis
Protein Biosynthesis/ Protein Modification
Other Metabolic Processes
Cellular Catabolism/ Protein Degradation
Cellular Respiration
Nucleotide Metabolism
Carbohydrate Metabolism
Organelle Organization
Cellular Regulation
Development/ Morphogenesis
P adjust
0.04
0.03
0.02
0.01
Enrichment Score
10
20
30
B
Description
Clusters
Categories
3 4
alpha-Linolenic acid metabolism (map00592)
Biosynthesis of unsaturated fatty acids (map01040)
Fatty acid degradation (map00071)
Fatty acid metabolism (map01212)
Phenylalanine metabolism (map00360)
cAMP signaling pathway (map04024)
PI3K-Akt signaling pathway (map04151)
Peroxisome (map04146)
Biosynthesis of secondary metabolites (map01110)
Cell cycle (map04110)
Cellular senescence (map04218)
Lipid Metabolism
Amino Acid metabolism
Cellular Regulation & Signaling
Cellular Organization
Specialized Metabolis
Cell Cycle & Division
P adjust
0.04
0.03
0.02
0.01
Enrichment Score
10
20
30
40

**Fig S7. Functional enrichment of aposymbiotic-specific rhythmic clusters corresponding to fig. S6A-B.**

(**A**) Gene Ontology biological process enrichment (adjusted $P < 0.05$) for the four aposymbiotic-specific rhythmic clusters (C1-C4) derived from Gaussian mixture model clustering of 467 rhythmic genes identified by RAIN. Enriched terms are grouped into functional categories, including lipid metabolism, homeostasis, protein biosynthesis and modification, other metabolic processes, cellular catabolism and protein degradation, cellular respiration, nucleotide metabolism, carbohydrate metabolism, organelle organization, cellular regulation, and development and morphogenesis. Dot size indicates enrichment score, and color represents adjusted *P* value. (**B**) KEGG pathway enrichment for the same clusters, with significant pathways (adjusted $P < 0.05$) categorized into lipid metabolism, amino acid metabolism, cellular regulation and signaling, cellular organization, specialized metabolism, and cell cycle and division. These enrichments provide functional characterization of the temporal modules formed by aposymbiotic-specific rhythmic genes.

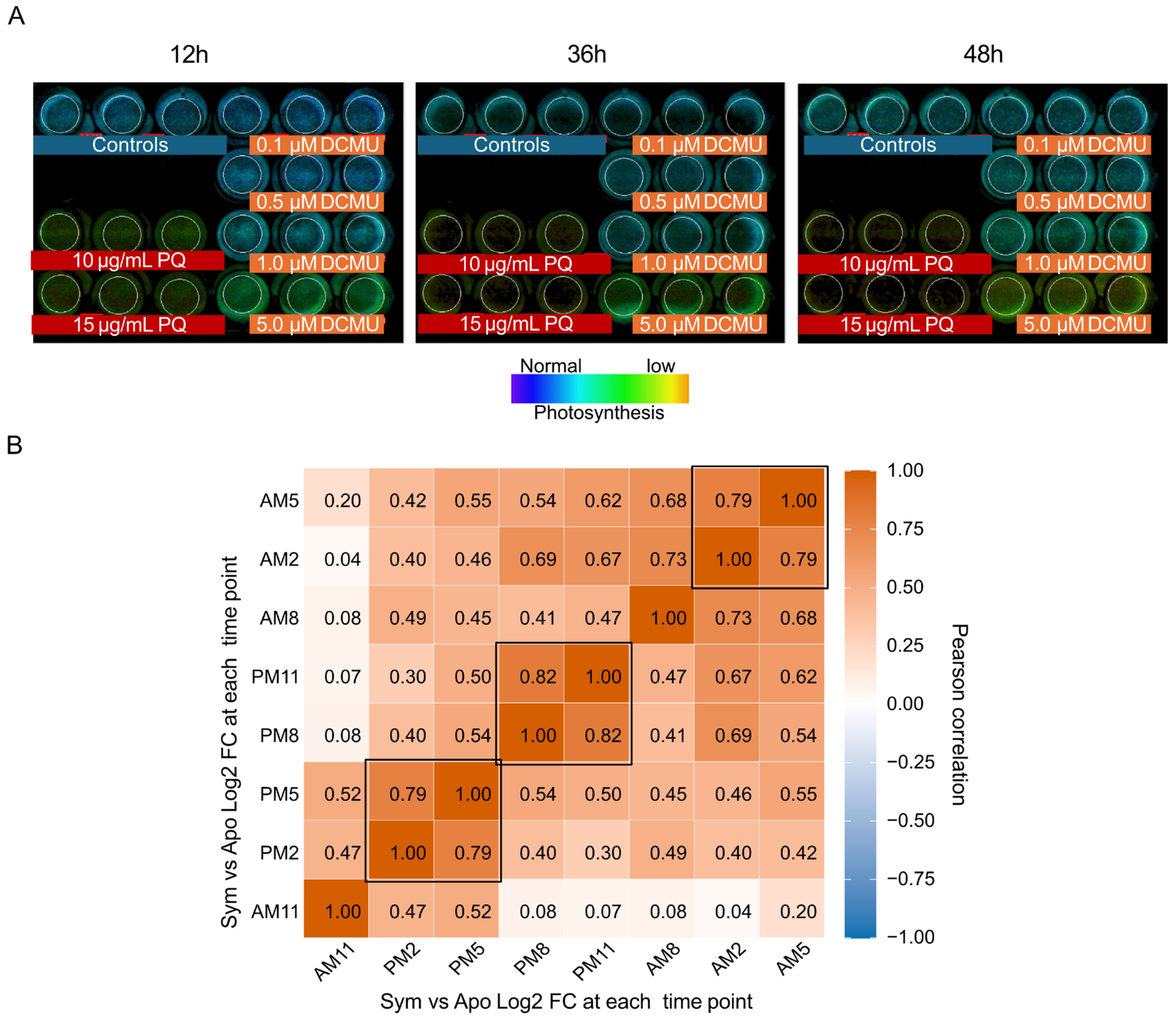


**Fig. S8. Photochemical response to paraquat and timepoint correlation analysis used for PQ experimental design.**

(**A**) Chlorophyll fluorescence images of symbiotic *P. bursaria* cultures exposed to control medium, DCMU (0.1-5.0 µM), or paraquat (10-15 µg/ml) at 12 h, 36 h, and 48 h. Fluorescence intensity is displayed on a pseudocolor scale from normal (blue) to low (yellow), illustrating the differential effects of PSII inhibition by DCMU and PSI inhibition by paraquat. (**B**) Pearson correlation matrix of Sym versus Apo $Log_2$ fold-change values across the eight diel timepoints (AM8, AM11, PM2, PM5, PM8, PM11, AM2, AM5). Blocks of high correlation ($r \geq 0.79$-1.00) identify sets of redundant timepoints, supporting selection of AM8, AM11, PM5, and PM11 as representative sampling points for the paraquat RNA-seq experiment.

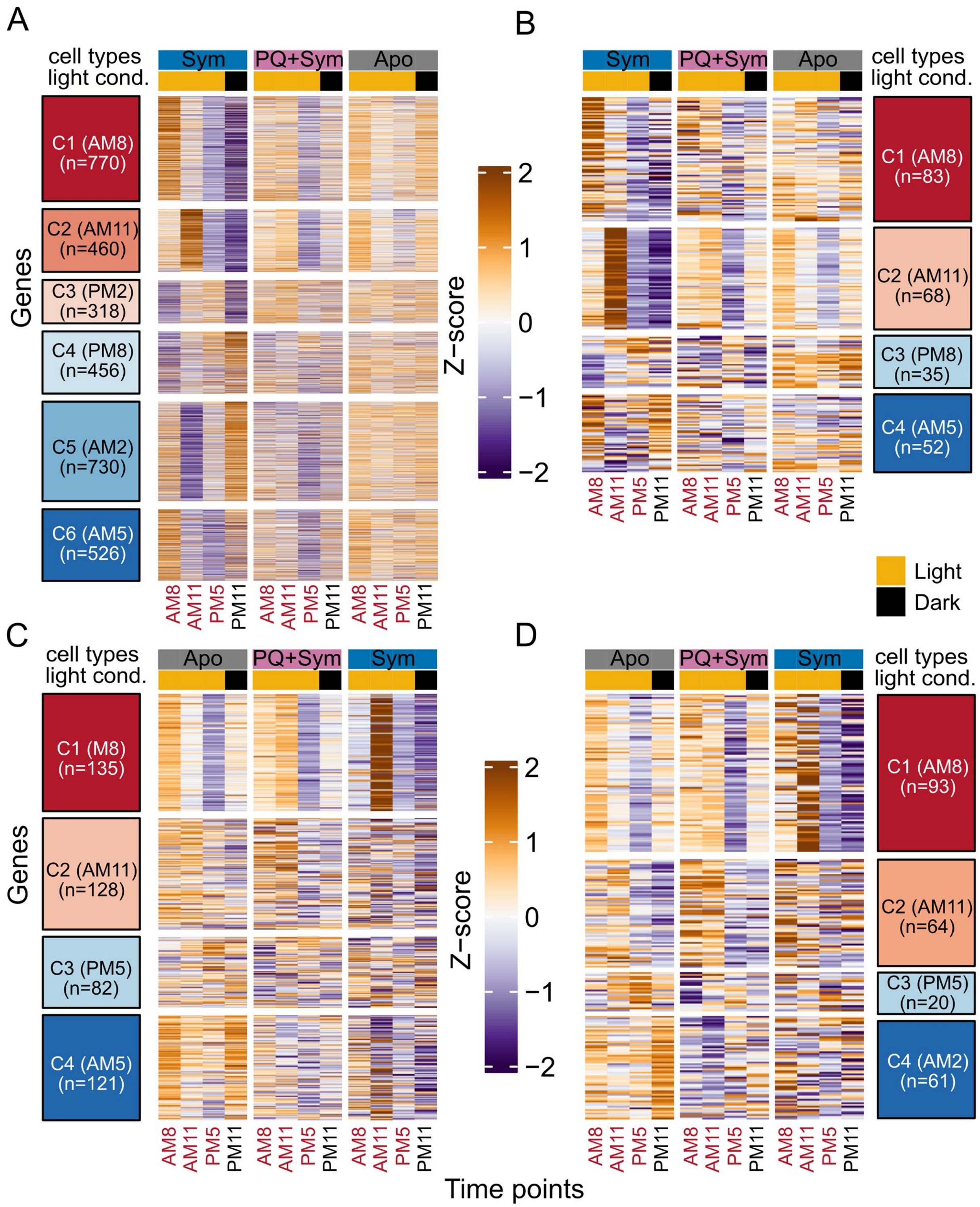
A
B
C
D
cell types
light cond.
Sym
PQ+Sym
Apo
Genes
C1 (AM8) (n=770)
C2 (AM11) (n=460)
C3 (PM2) (n=318)
C4 (PM8) (n=456)
C5 (AM2) (n=730)
C6 (AM5) (n=526)
C1 (AM8) (n=83)
C2 (AM11) (n=68)
C3 (PM8) (n=35)
C4 (AM5) (n=52)
C1 (M8) (n=135)
C2 (AM11) (n=128)
C3 (PM5) (n=82)
C4 (AM5) (n=121)
C1 (AM8) (n=93)
C2 (AM11) (n=64)
C3 (PM5) (n=20)
C4 (AM2) (n=61)
Z−score
2
1
0
−1
−2
AM8
AM11
PM5
PM11
Light
Dark
Time points

**Fig. S9. Gene-level expression heatmaps for temporal clusters shown as mean profiles in Fig. 3C-F.**

(**A**) Heatmaps of Z-scaled $\log_2$(TPM+1) expression for the six symbiotic-specific rhythmic clusters (C1-C6) across symbiotic (Sym), paraquat-treated symbiotic (PQ+Sym), and aposymbiotic (Apo) samples at the four matched zeitgeber timepoints (AM8, AM11, PM5, PM11). (**B**) Heatmaps of the four core rhythmic clusters (C1-C4) displayed across the same three conditions and four matched timepoints. (**C**) Heatmaps of the four aposymbiotic-specific rhythmic clusters (C1-C4) shown across Apo, PQ+Sym, and Sym samples at the same matched timepoints. (**D**) Heatmaps of the four paraquat-response clusters (C1-C4), summarizing distinct PQ-associated temporal response patterns across Apo, PQ+Sym, and Sym samples. For all panels, columns are ordered by time-of-day and annotated by cell type and light-dark condition as indicated; cluster labels denote the phase grouping used for the corresponding mean trajectory plot

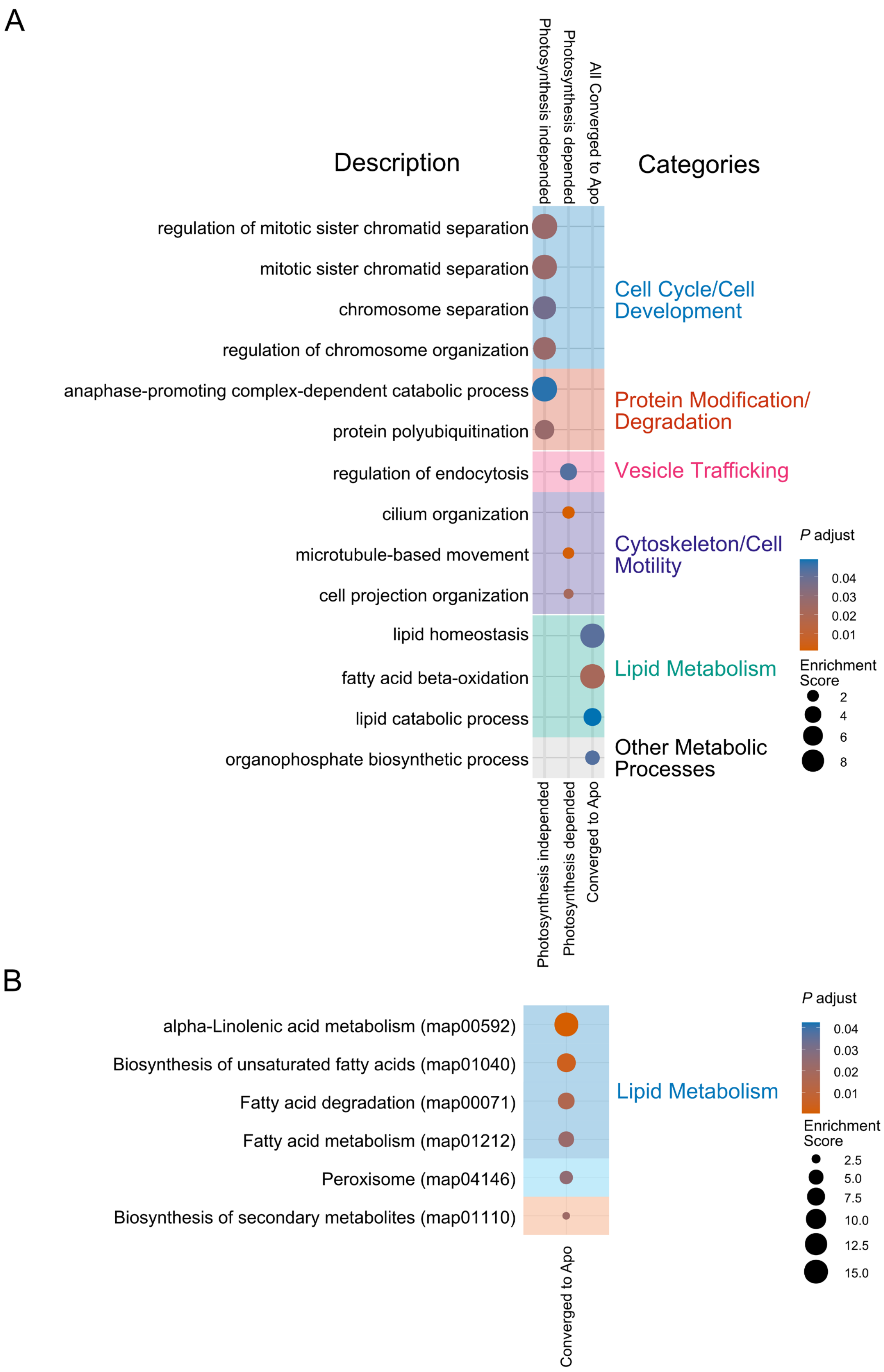
A
Description
Categories
Photosynthesis independed
Photosynthesis depended
All Converged to Apo
regulation of mitotic sister chromatid separation
mitotic sister chromatid separation
chromosome separation
regulation of chromosome organization
anaphase-promoting complex-dependent catabolic process
protein polyubiquitination
regulation of endocytosis
cilium organization
microtubule-based movement
cell projection organization
lipid homeostasis
fatty acid beta-oxidation
lipid catabolic process
organophosphate biosynthetic process
Cell Cycle/Cell Development
Protein Modification/ Degradation
Vesicle Trafficking
Cytoskeleton/Cell Motility
Lipid Metabolism
Other Metabolic Processes
P adjust
0.04
0.03
0.02
0.01
Enrichment Score
2
4
6
8
Photosynthesis independed
Photosynthesis depended
Converged to Apo
B
alpha-Linolenic acid metabolism (map00592)
Biosynthesis of unsaturated fatty acids (map01040)
Fatty acid degradation (map00071)
Fatty acid metabolism (map01212)
Peroxisome (map04146)
Biosynthesis of secondary metabolites (map01110)
Lipid Metabolism
P adjust
0.04
0.03
0.02
0.01
Enrichment Score
2.5
5.0
7.5
10.0
12.5
15.0
Converged to Apo

**Fig. S10. Functional enrichment of paraquat-response categories corresponding to Fig. 4.** (**A**) Gene Ontology biological process enrichment (adjusted P < 0.05) for genes assigned to the PQ-insensitive, PQ-sensitive, and converged-to-aposymbiotic paraquat-response categories defined in Figure 4. Enriched terms are grouped into categories such as lipid metabolism, protein modification and degradation, vesicle trafficking, cytoskeleton and cell motility, other metabolic processes, and cell cycle and cell development. Dot size represents enrichment score, and color denotes adjusted P value. (**B**) KEGG pathway enrichment (adjusted $P < 0.05$) for the converged-to-aposymbiotic category, with significant pathways including fatty acid metabolism, fatty acid degradation, peroxisome-related processes, unsaturated fatty acid biosynthesis, α-linolenic acid metabolism, and secondary metabolite biosynthesis. Dot size indicates enrichment score, and color represents adjusted $P$ value.

| Domain Family | Total Genes | Total Rhythmic | Symbiotic-Specific | Aposymbiotic-Specific | Core Rhythmic | % Symbiotic-Specific |
|---|---|---|---|---|---|---|
| BTB/POZ domain protein | 25 | 4 | 4 | 0 | 0 | 100.0% |
| Casein kinase | 102 | 17 | 15 | 1 | 1 | 88.2% |
| EF-hand $Ca^{2+}$-binding protein | 637 | 86 | 70 | 11 | 5 | 81.4% |
| F-box protein | 35 | 4 | 4 | 0 | 0 | 100.0% |
| F-box protein; WD40 repeat protein | 12 | 1 | 1 | 0 | 0 | 100.0% |
| Kelch repeat protein | 89 | 16 | 12 | 4 | 0 | 75.0% |
| Kelch repeat protein; BTB/POZ domain protein | 8 | 2 | 2 | 0 | 0 | 100.0% |
| MYB transcription factor | 188 | 20 | 14 | 4 | 2 | 70.0% |
| MYB transcription factor; EF-hand $Ca^{2+}$-binding protein | 3 | 2 | 0 | 2 | 0 | 0.0% |
| PAS domain protein | 65 | 8 | 7 | 1 | 0 | 87.5% |
| WD40 repeat protein | 745 | 77 | 66 | 6 | 5 | 85.7% |
| WD40 repeat protein; EF-hand $Ca^{2+}$-binding protein | 20 | 2 | 2 | 0 | 0 | 100.0% |
| **TOTAL** | **1,929** | **239 (12.4%)** | **197 (82.4%)** | **29 (12.1%)** | **13 (5.4%)** | **82.4%** |

**Table S1. Rhythmicity of circadian-associated domain families across symbiotic-specific, aposymbiotic-specific, and core gene categories.**

Of 239 rhythmic domain genes identified (12.4% of 1,929 curated circadian-associated domains), 197 (82.4%) are symbiotic-specific, 29 (12.1%) are aposymbiotic-specific, and 13 (5.4%) are core rhythmic. The table includes single-domain families and multi-domain gene products. See Supporting Data 6 for complete gene-level annotations.

| Gene Set | Mean Correlation | Median Correlation | Standard Deviation | N Genes |
|---|---|---|---|---|
| Symbiotic-specific (vs sym reference) | 0.230 | 0.372 | 0.568 | 3,260 |
| Aposymbiotic-specific (vs apo reference) | 0.494 | 0.688 | 0.491 | 466 |
| Core rhythmic (vs sym reference) | 0.373 | 0.588 | 0.554 | 238 |
| Core rhythmic (vs apo reference) | 0.597 | 0.771 | 0.460 | 238 |
| **Test** | | | **Raw P** | **BH-adjusted P** |
| Symbiotic-specific vs 0 | | | 7.62E-100 | 4.57E-99 |
| Aposymbiotic-specific vs 0 | | | 5.69E-50 | 1.71E-49 |
| Core rhythmic (Sym) vs 0 | | | 3.12E-17 | 3.75E-17 |
| Core rhythmic (Apo) vs 0 | | | 1.67E-32 | 3.35E-32 |
| Sym-specific vs Apo-Specific | | | 1.14E-24 | 1.71E-24 |
| Core sym vs Core Apo | | | 3.83E-08 | 3.83E-08 |

**Table S2. Pearson correlation of paraquat-treated four-timepoint expression profiles to symbiotic-specific, aposymbiotic-specific, and core rhythmic reference temporal profiles, with Wilcoxon tests.**

PQ-treated symbiotic cells (sampled at AM8, AM11, PM5, PM11) were evaluated across 3,260 symbiotic-specific, 466 aposymbiotic-specific, and 238 core rhythmic host genes. For each gene set, Pearson correlations (r) were computed between PQ-treated profiles and the corresponding 8-timepoint symbiotic or aposymbiotic reference temporal profiles and summarized by mean/median r and standard deviation. Statistical support is provided by distribution-level Wilcoxon tests (BH-corrected): one-sample signed-rank tests assess whether median r differs from 0 (tests vs 0), and rank-sum tests compare correlation distributions between symbiotic-specific vs aposymbiotic-specific sets and between core correlations computed against symbiotic vs aposymbiotic references. Because correlations are estimated from only four timepoints, inference is based on gene-set–level tests rather than per-gene correlation *P* values.

| Domain Family | Abolished | Partially Disrupted | Phase Inverted | Maintained | Reduced Amplitude | Increased Amplitude | Total |
|---|---|---|---|---|---|---|---|
| EF-hand $Ca^{2+}$-binding protein | 19 | 33 | 11 | 9 | 11 | 3 | 86 |
| WD40 repeat protein | 30 | 24 | 10 | 4 | 6 | 2 | 76 |
| MYB transcription factor | 4 | 9 | 2 | 0 | 3 | 0 | 18 |
| Casein kinase | 4 | 7 | 2 | 1 | 3 | 0 | 17 |
| Kelch repeat protein | 2 | 8 | 2 | 2 | 2 | 0 | 16 |
| PAS domain protein | 2 | 4 | 1 | 1 | 0 | 0 | 8 |
| F-box protein | 2 | 2 | 0 | 0 | 0 | 0 | 4 |
| BTB/POZ domain protein | 0 | 2 | 1 | 1 | 0 | 0 | 4 |
| Kelch repeat protein; BTB/POZ domain protein | 1 | 1 | 0 | 0 | 0 | 0 | 2 |
| MYB transcription factor; EF-hand $Ca^{2+}$-binding protein | 0 | 0 | 0 | 0 | 2 | 0 | 2 |
| WD40 repeat protein; EF-hand $Ca^{2+}$-binding protein | 0 | 2 | 0 | 0 | 0 | 0 | 2 |
| F-box protein; WD40 | 0 | 1 | 0 | 0 | 0 | 0 | 1 |

| | | | | | | | |
|---|---|---|---|---|---|---|---|
| repeat protein | | | | | | | |
| **TOTAL** | **64 (27.1%)** | **77 (32.6%)** | **26 (11.0%)** | **16 (6.8%)** | **22 (9.3%)** | **13 (5.5%)** | **236** |

**Table S3. Distribution of circadian-associated domain genes across PQ disruption categories.**

Of 236 circadian-associated domain genes identified in symbiotic-specific, aposymbiotic-specific, and core rhythmic gene sets (derived from the 239 rhythmic domain genes in Supporting Table S1, excluding 3 bZIP genes not included in the original curation), 167 (70.8%) were preferentially disrupted under photosynthesis inhibition (Abolished + Partially Disrupted + Phase Inverted categories), while only 69 (29.2%) were maintained, dampened, or amplified. Domain families traditionally implicated in eukaryotic circadian control—EF-hand $Ca^{2+}$-binding proteins (22.1% abolished, 38.4% partially disrupted), WD40 repeat proteins (39.5% abolished, 31.6% partially disrupted), and casein kinases (23.5% abolished, 41.2% partially disrupted)—showed the highest representation in disrupted categories, indicating that photosynthetic input is required to maintain the stability of core circadian-associated protein machinery.

| Domain Family | Abolished (%) | Partially Disrupted (%) | Phase Inverted (%) | Total Disrupted (%) | Maintained (%) | Reduced Amplitude (%) | Increased Amplitude (%) |
|---|---|---|---|---|---|---|---|
| WD40 repeat protein | 39.5 | 31.6 | 13.2 | **84.2** | 5.3 | 7.9 | 2.6 |
| Casein kinase | 23.5 | 41.2 | 11.8 | **76.5** | 5.9 | 17.6 | 0.0 |
| EF-hand $Ca^{2+}$-binding protein | 22.1 | 38.4 | 12.8 | **73.3** | 10.5 | 12.8 | 3.5 |
| F-box protein | 50.0 | 50.0 | 0.0 | **100.0** | 0.0 | 0.0 | 0.0 |
| PAS domain protein | 25.0 | 50.0 | 12.5 | **87.5** | 12.5 | 0.0 | 0.0 |
| Kelch repeat protein | 12.5 | 50.0 | 12.5 | **75.0** | 12.5 | 12.5 | 0.0 |
| MYB transcription factor | 22.2 | 50.0 | 11.1 | **83.3** | 0.0 | 16.7 | 0.0 |
| BTB/POZ domain protein | 0.0 | 50.0 | 25.0 | **75.0** | 25.0 | 0.0 | 0.0 |
| **ALL DOMAINS** | **27.1** | **32.6** | **11.0** | **70.8** | **6.8** | **9.3** | **5.5** |

**Table S4. Percentage breakdown of circadian-associated domain gene responses to photosynthesis inhibition.**

For each domain family, the table shows the percentage of genes falling into each disruption category. Across all domain families, 70.8% of genes were disrupted (abolished, partially disrupted, or phase-inverted), with WD40 repeat proteins (84.2%), PAS domain proteins (87.5%), F-box proteins (100.0%), and MYB transcription factors (83.3%) showing particularly high disruption rates. Conversely, only 6.8% of domain genes were fully maintained, and 5.5% showed increased amplitude under PQ treatment. This pattern demonstrates that circadian-associated protein machinery is systematically destabilized when photosynthetic energy input is removed, indicating that the post-translational regulatory infrastructure underpinning temporal organization in *P. bursaria* depends fundamentally on endosymbiotic photosynthetic activity.
